# Bioinformatics analysis of next generation sequencing data reveals novel biomarkers and signaling pathways associated with recurrent implantation failure

**DOI:** 10.1101/2025.02.13.638207

**Authors:** Basavaraj Vastrad, Chanabasayya Vastrad

## Abstract

Recurrent implantation failure (RIF) is a cases in which women have had three fruitless in vitro fertilization (IVF) bid with positive quality embryos. The RIF originates from uterine endometrium microbiota has been implicated in reproductive failure, and poor prognosis and lacks effective treatment. Efforts have been made to elucidate the molecular pathogenesis of RIF. To identify key genes and signaling pathways in RIF, the next genetation sequencing data GSE243550 was downloaded from the Gene Expression Omnibus (GEO) database. The differentially expressed genes (DEGs) between RIF and normal controls samples were identified using t-tests in the limma R bioconductor package. Using the DEGs, we further performed a series of gene ontology (GO) and REACTOME pathway enrichment analyses. Protein-protein interaction network was derived using the IMex interactome database and visualized using Cytoscape software. The most significant modules from the PPI network were selected for GO and pathway enrichment analysis. A miRNA-hub gene regulatory network and TF-hub gene regulatory network were constructed depending on key hub genes and visualized using Cytoscape software. A receiver operating characteristic curve (ROC) analysis was plotted to diagnose RIF. In total, 958 DEGs were identified, of which 479 were up regulated genes and 479 were down regulated genes. GO and REACTOME pathway enrichment analysis results revealed that the upregulated genes were mainly enriched in multicellular organismal process, membrane, small molecule binding and extracellular matrix organization, whereas downregulated genes were mainly enriched in organonitrogen compound metabolic process, intracellular anatomical structure, catalytic activity and translation. Through analyzing the PPI network, we screened hub genes APP, HSP90AA1, CAND1, CUL1, HSP90AB1, SIRT7, SRC, CDKN1A, ISG15 and RPS16 by the Cytoscape software. The regulatory network analysis revealed that microRNAs (miRNAs) include hsa-miR-574-3p and hsa-mir-208a-3p, and transcription factors (TFs) include SREBF1 and RELA might be involved in the development of RIF. Receiver operating characteristic curve analysis demonstrated that the hub genes screened for RIF were of good diagnostic significance. Overall, these results thus highlight a range of novel signaling pathways and genes that are linked to the incidence and progression of RIF, providing a list of important diagnostic and prognostic molecular markers that have the potential to aid in the clinical diagnosis and treatment of RIF.

## Introduction

Successful implantation involves the synchronized and interdependent crosstalk between the embryo and endometrium, which is the essential point for a fruitful pregnancy [1]. At present, in vitro fertilization-embryo transfer (IVF-ET) is considered the key approach for treating infertility with improvements in laboratory procedures and ovarian stimulation [2]. Recurrent implantation failure (RIF) is the failure to achieve a clinical pregnancy after transfer of at least four good-quality embryos in a minimum of three fresh or frozen cycles in a woman < 40 years of age [3]. RIF affects only 10% women undergoing IVF-ET, which places a great difficulty on the economy of health and reduces quality of life [4]. The pathological manifestations of RIF are characterized by attachment and migration process, with a negative urine or blood test for human chorionic gonadotropin (hCG) or failure to form an intrauterine gestational sac with positive hCG [5]. Regarding the pathogenesis of RIF, the most widely discussed factors include the hyperparathyroidism [6], insulin resistance [7], obesity [8], preeclampsia [9], male factor [10], endometriosis [11], polycystic ovarian syndrome [12], inflammatory factors [13], autoimmunity [14], endometrial cancer [15], endometrial receptivity [16], psychosocial consequences [17], inherited thrombophilia [18], prothrombotic genetic variants [19], chromosomal abnormalities [20], viral infections [21], hyperprolactinaemia [22], immunomodulation [23], embryo aneuploidy [24], endometrial microbiome [25], uterine fibroids [26], sperm DNA damage [27], oxidative stress [28] and Male factor infertility [29]. Studies have revealed that the progression of RIF is related to genetic factors [30]. Therefore, molecular changes in the occurrence and development of RIF and are effective ways to explore the molecular pathogenesis of RIF.

Despite improvement in diagnosis and treatment, the prognosis of RIF patients remains poor, which has become an active topic of clinical and basic research. Genetic mutations [31], epigenetic alterations [32] and aberrant molecular signaling pathways [33] are involved in the processes of RIF. In particular, the new molecular characteristics can be applied in early risk assessment, the identification of better specific biomarkers, and the improvement of clinic treatment and survival. In the treatment of RIF, early diagnosis is of great significance, so it is necessary to dig out more RIF biomarkers. New biomarkers such as KIR and LILRB [34], ERAP/HLA-C and KIR [35], KIR2DL4 [36], STAT3, IL-1β, IL-6 and TNF-α [37], and MTHFR and TS [38] have also been widely concerned and studied. Previous reports on this topic have focused on identification of potential signaling pathways such as Wnt4/β-catenin signaling pathway [39], NF-κB signaling pathway [40], JAK/STAT3 signaling pathway [41], Met/PI3K/Akt signaling pathway [42] and ADCY1/cAMP signalling pathway [43]. The identification of novel biomarkers and signaling pathways may be helpful to improve the clinical outcome of RIF patients.

With the rapid advancement of bioinformatics and next generation sequencing (NGS) technology, plenty of clinical and genetic data are disclosed, which provides rich resources for basic and clinical investigation of various diseases and disorders [44–45]. Through various public databases, we can further investigation the molecular pathogenesis of RIF and explore novel RIF biomarkers and potential therapeutic targets of RIF. At present, NGS dataset GSE243550 downloaded from Gene Expression Omnibus (GEO) (https://www.ncbi.nlm.nih.gov/geo) [46] database can be used to detect genes transcription expression levels and to provide the technical support for monitoring mRNA expression and cell function prediction in RIF. Non-biased bioinformatics analyses, including identification of DEGs, gene ontology (GO) and REACTOME pathway enrichment analyses, and protein–protein interaction (PPI) network and module analysis, and miRNA-hub gene regulatory network and TF-hub gene regulatory network analysis were conducted, and the findings were further validated by receiver operating characteristic (ROC) curve analysis. Based on the obtained results, we propose novel biomarkers as potential targets for the diagnosis and treatment of RIF

## Materials and Methods

### Next generation sequencing (NGS) data source

GSE243550 NGS dataset was downloaded from the GEO database. The GSE243550 dataset was generated utilizing the GPL24676 [Illumina NovaSeq 6000 (Homo sapiens)] platform and contained 20 RIF samples and 20 normal control samples.

### Identification of DEGs

The statistically significant DEGs were selected by moderated t test approach with limma [47] package of R bioconductor. We screened DEGs between RIF and normal control samples by utilizing limma package with a adjust p < 0.05, and a log (Fold Change) > 0.1343 for up regulated genes and log (Fold Change) < - 0.1554 for down regulated genes. We adjusted p-value to correct the false positive error caused by the multiple tests and determined it by the Benjamini & Hochberg method [48], which is one of the suitable tools to reduce the false discovery rate. The DEGs are presented as volcano plot, generated using ggplot2 package of R bioconductor. The gplot package of R bioconductor was used to construct a heatmap of the DEGs.

### GO and pathway enrichment analyses of DEGs

In order to further investigation the role of DEGs in the occurrence and advancement of RIF, the GO (http://www.geneontology.org) [49] and REACTOME pathway (https://reactome.org/) [50] enrichment analyses were carried out through online tool g:Profiler (http://biit.cs.ut.ee/gprofiler/) [51]. A p < 0.05 was marked as the threshold value. Through GO enrichment analysis, biological processes (BP), cellular components (CC) and the molecular functions (MF) associated in various genes were interpreted. REACTOME pathway enrichment analysis was conducted to explore the role of DEGs in different signaling pathway in the homo sapiens.

### Construction of the PPI network and module analysis

IMex interactome (https://www.imexconsortium.org/) [52], an open source online tool, was used to construct a PPI network, and Cytoscape (version 3.10.3) (http://www.cytoscape.org/) [53] software was implemented to complete visualization. To screen the hub genes that might be associated in RIF, we applied the Network Analyzer plug-in, using different parameters such as node degree [54], betweenness [55], stress [56] and closeness [57]. In addition, PEWCC plug-in [58] in cytoscape was used to identify the significant modules of the PPI network.

### Construction of the miRNA-hub gene regulatory network

The hub genes obtained from PPI network were uploaded to miRNet (https://www.mirnet.ca/) [59] to construct miRNA-hub gene regulatory network. The miRNA-hub gene regulatory network was constructed based on the TarBase, miRTarBase, miRecords, miRanda, miR2Disease, HMDD, PhenomiR, SM2miR, PharmacomiR, EpimiR, starBase, TransmiR, ADmiRE, and TAM 2.0 databases via the miRNet platform. The Cytoscape (version 3.10.3) [53] software was used to visualize the miRNA-hub gene regulatory network.

### Construction of the TF-hub gene regulatory network

The hub genes obtained from PPI network were uploaded to NetworkAnalyst (https://www.networkanalyst.ca/) [60] to construct TF-hub gene regulatory network. The TF-hub gene regulatory network was constructed based on the JASPAR database via the NetworkAnalyst platform. The Cytoscape (version 3.10.3) [53] software was used to visualize the TF-hub gene regulatory network.

### Receiver operating characteristic curve (ROC) analysis

To verify the sensitivity and specificity of the hub gene expression obtained from the PPI network analysis for the diagnosis of RIF, we plotted the receiver operator characteristic (ROC) curve analysis and calculated the corresponding area under the curve (AUC) of hub genes based on the RIF data. The ROC curve was created to quantitatively determine the sensitivity and specificity of hub genes using pROC package of R bioconductor [61]. AUC > 0.8 was considered the excellent diagnostic value.

## Results

### Identification of DEGs

The RIF NGS dataset GSE243550 was downloaded from the GEO database. After screening with the threshold of an adjusted p < 0.05, and a log (Fold Change) > 0.1343 for up regulated genes and log (Fold Change) < −0.1554 for down regulated genes, a total of 958 (479 up regulated and 479 down regulated genes) DEGs were screened from GSE243550 using the “limma” package in R bioconductor (Table 1), respectively. As shown in Fig.1 and Fig.2, the volcano plot and heatmap analyses were used to visualize the DEGs.

**Fig. 1.**
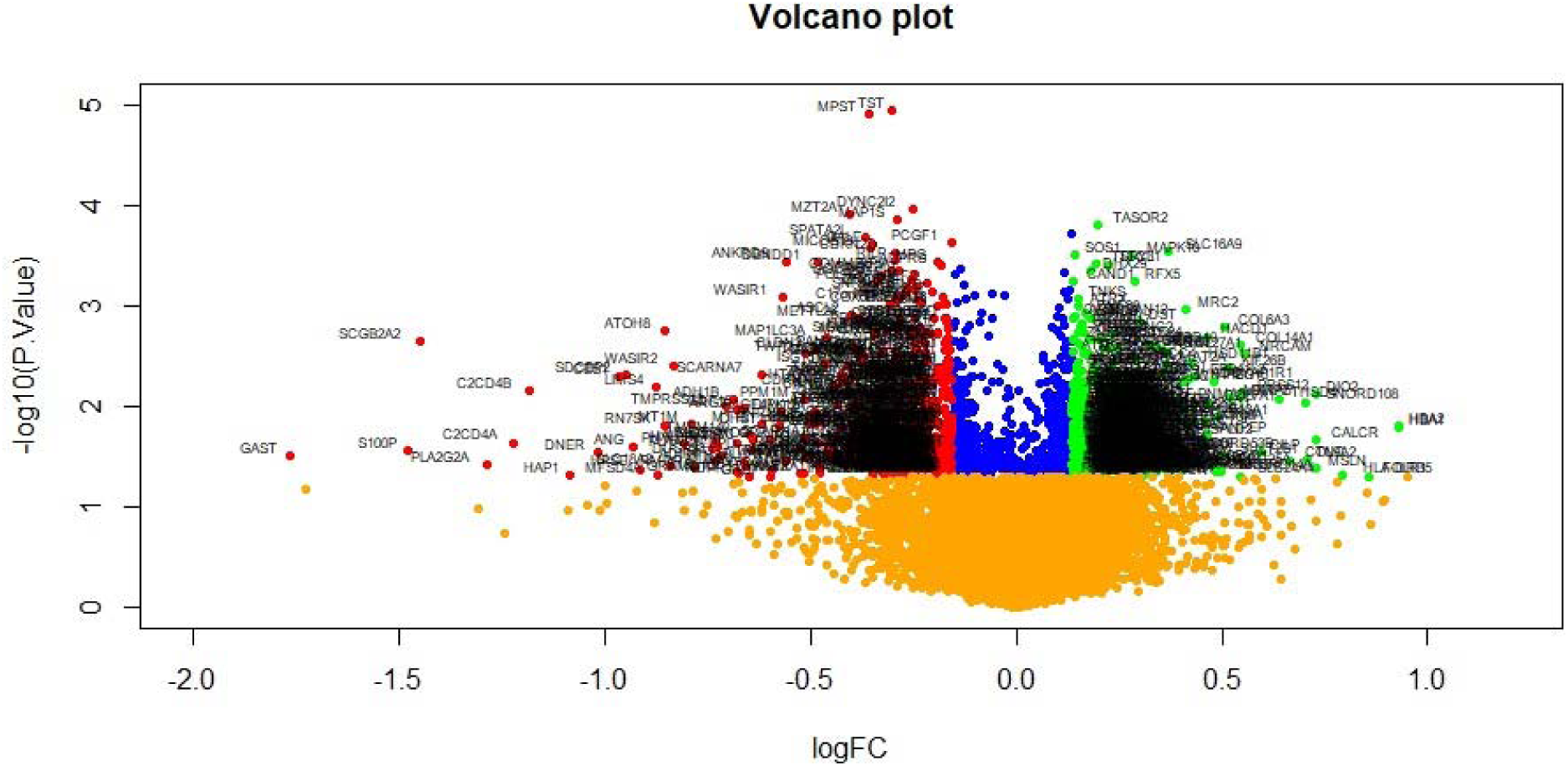
Volcano plot of differentially expressed genes. Genes with a significant change of more than two-fold were selected. Green dot represented up regulated significant genes and red dot represented down regulated significant genes.

**Fig. 2.**
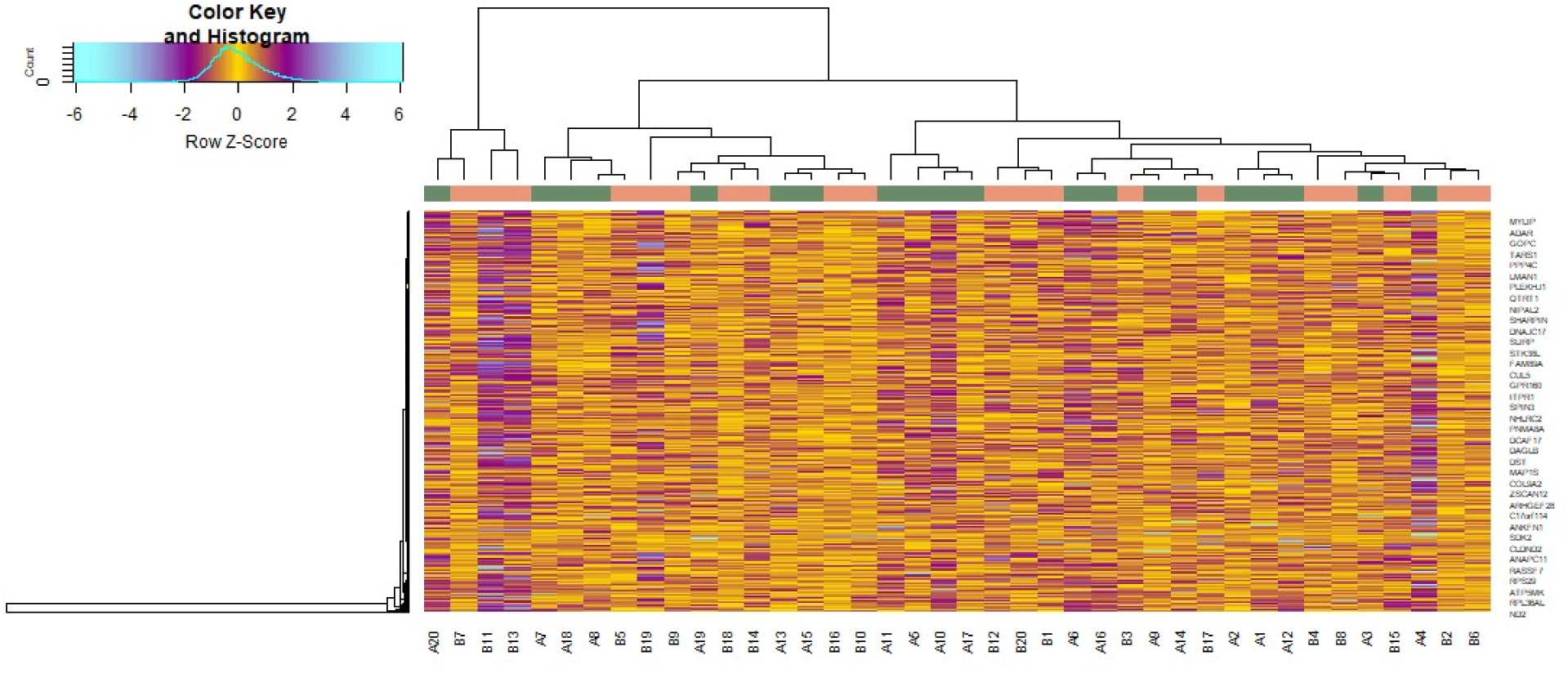
Heat map of differentially expressed genes. Legend on the top left indicate log fold change of genes. (A1 – A20 = RIF samples; B1 – B20= Normal control samples)

**Table 1.**
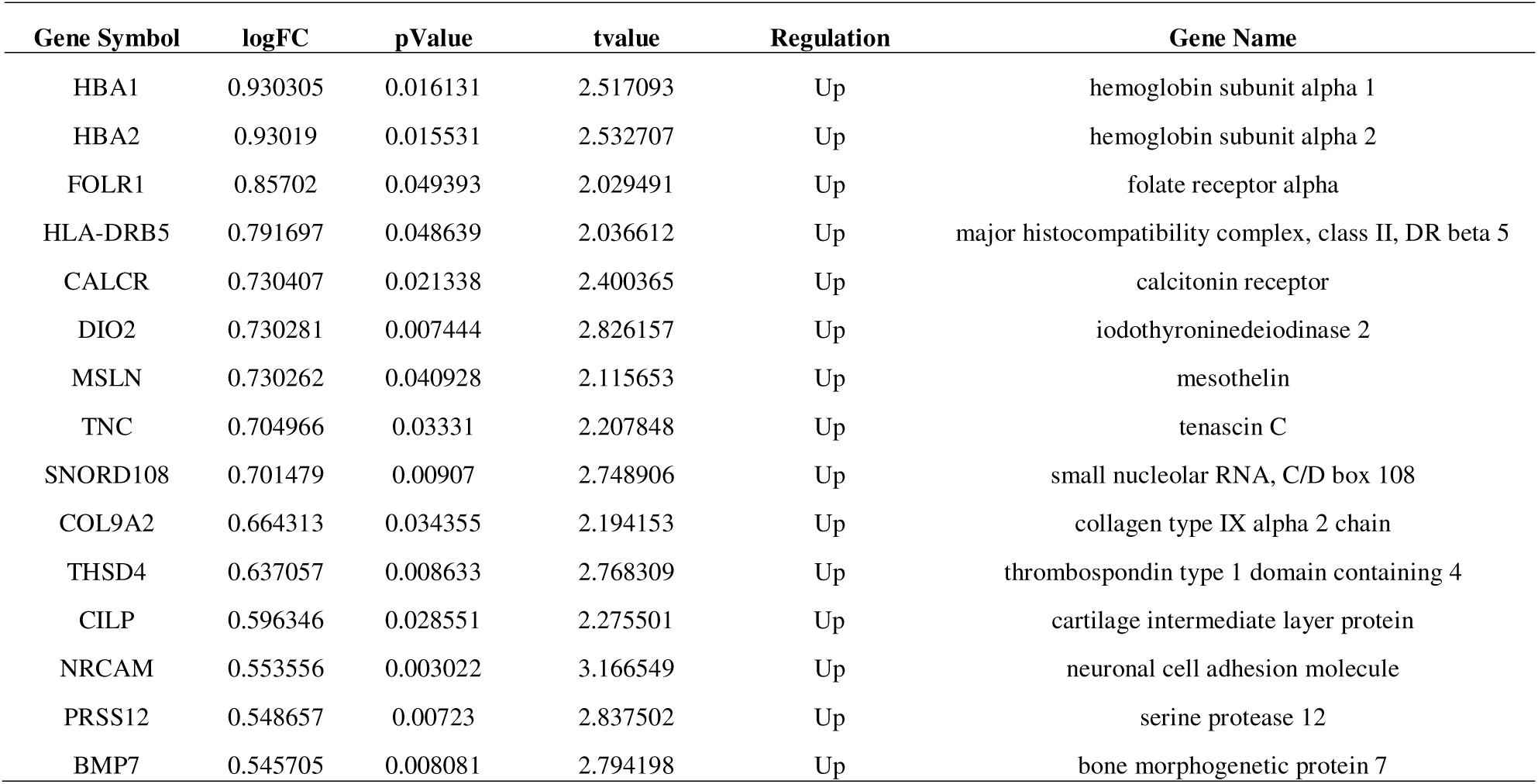
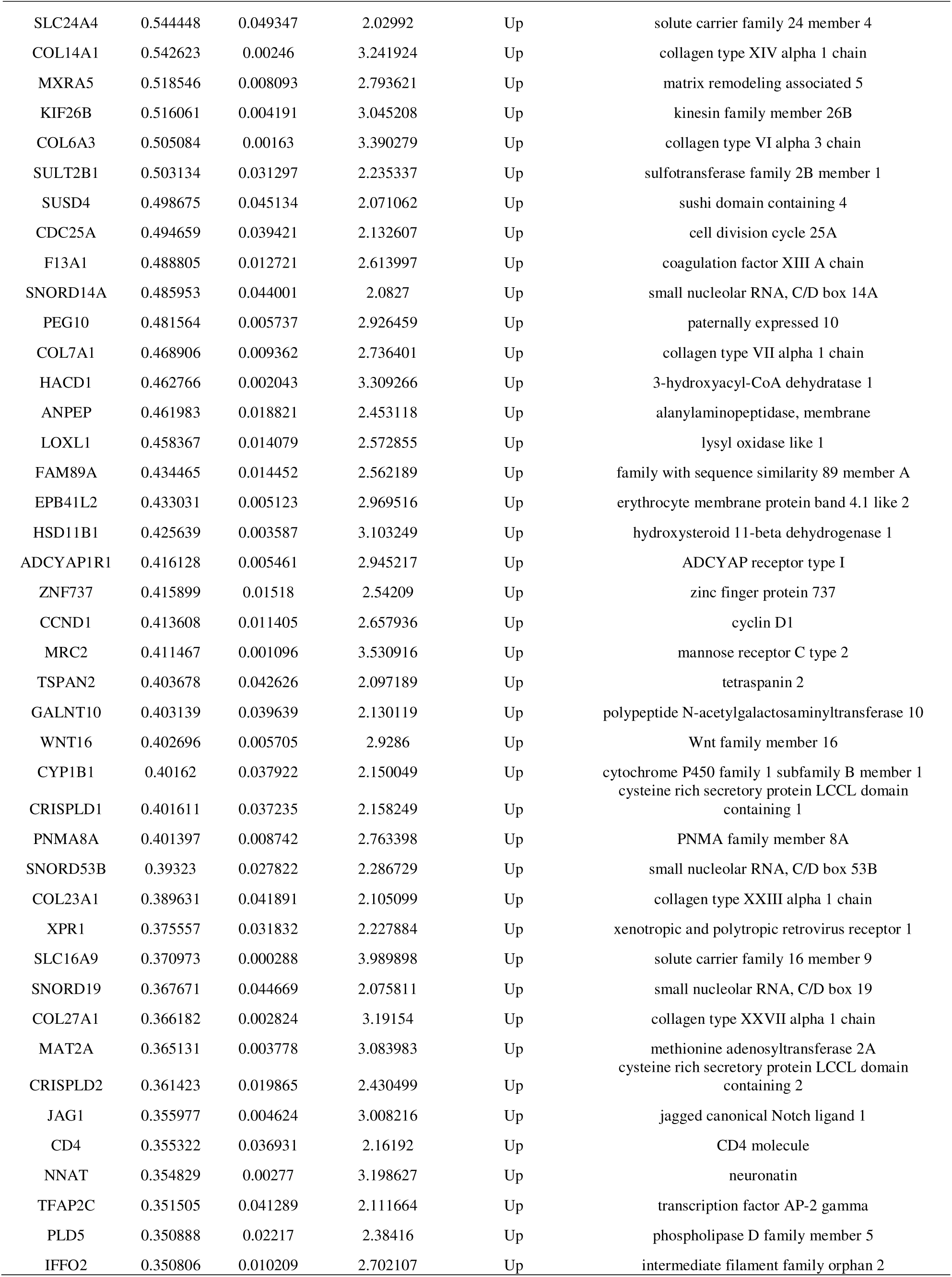
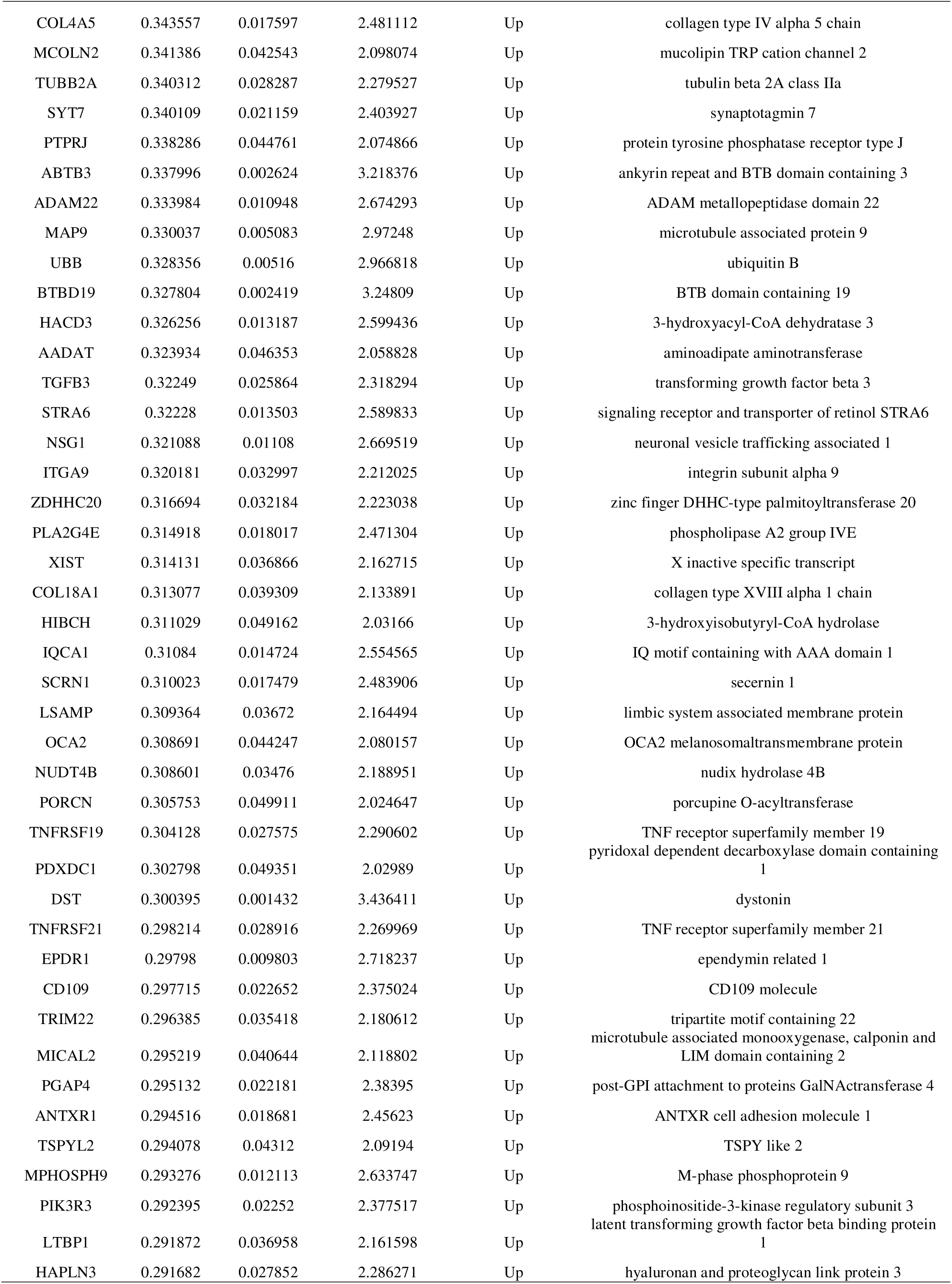
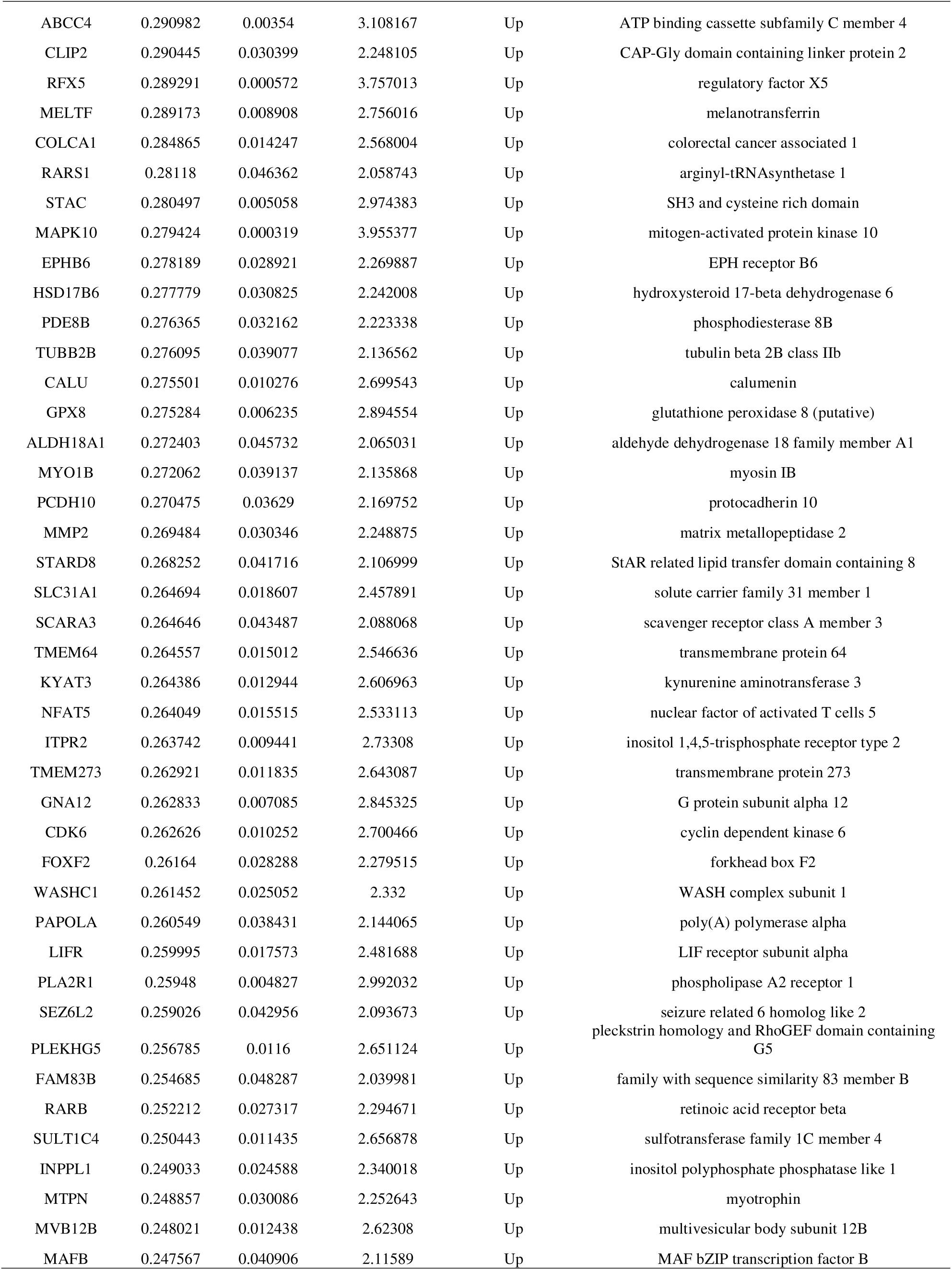
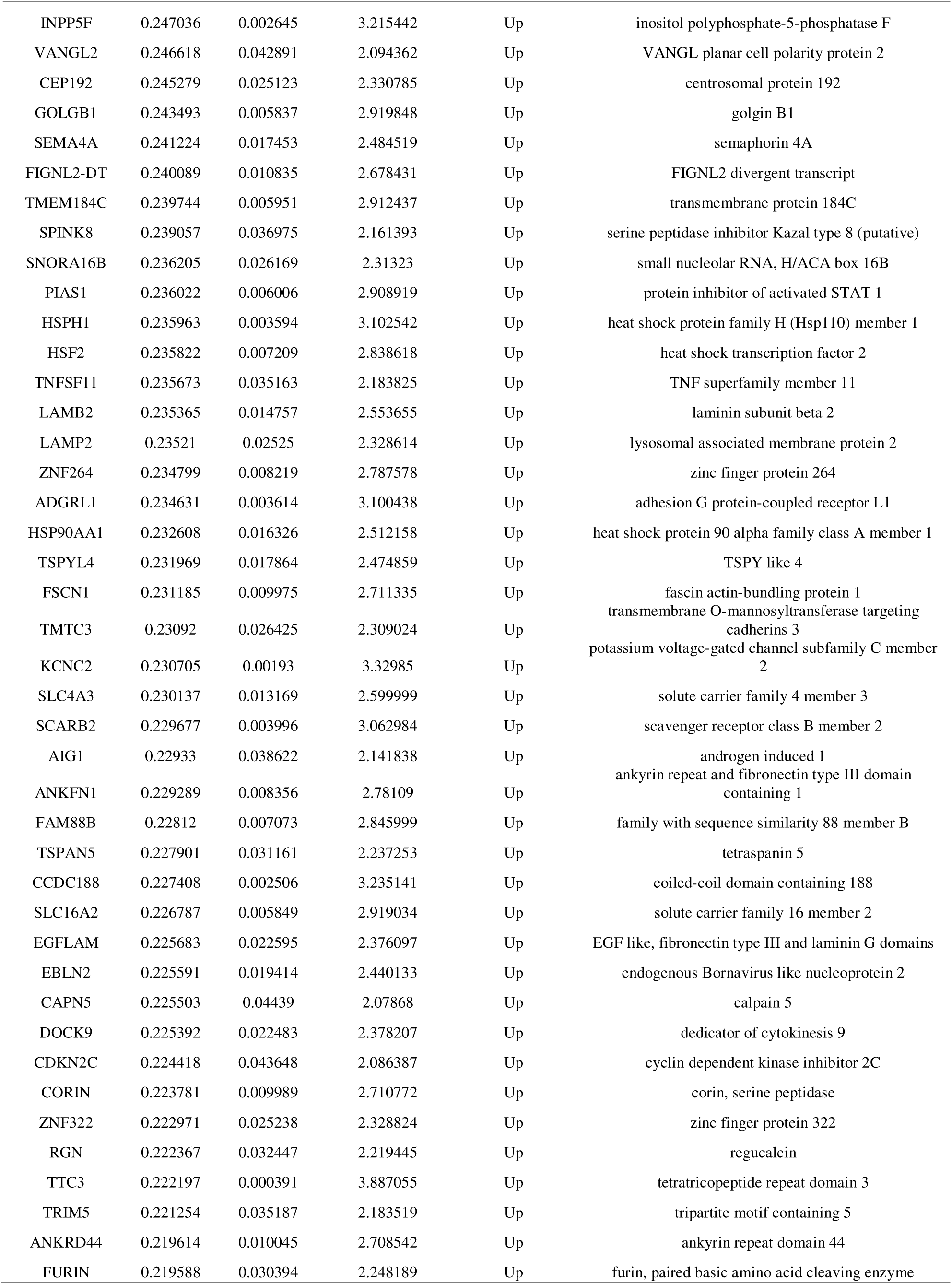
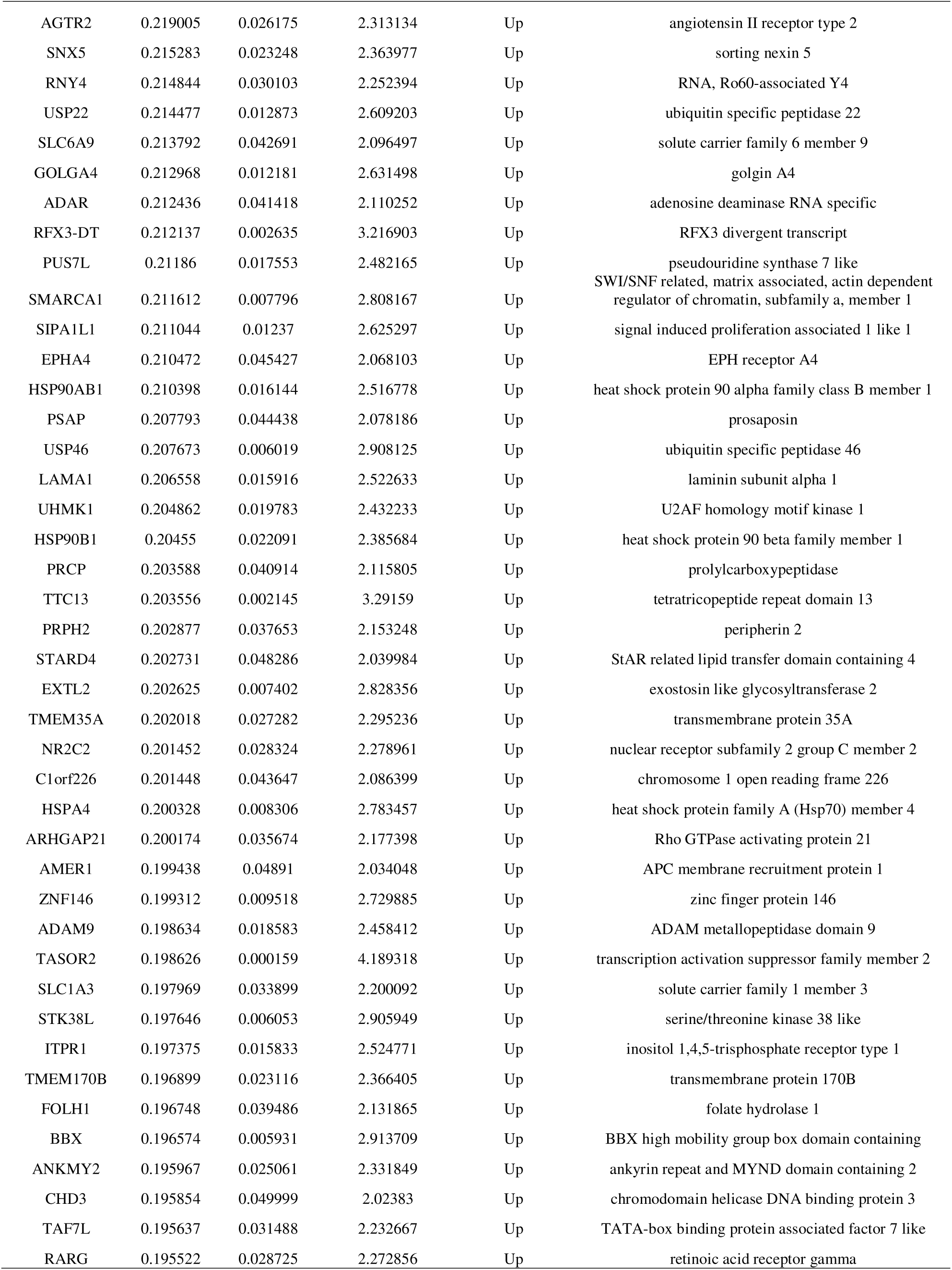
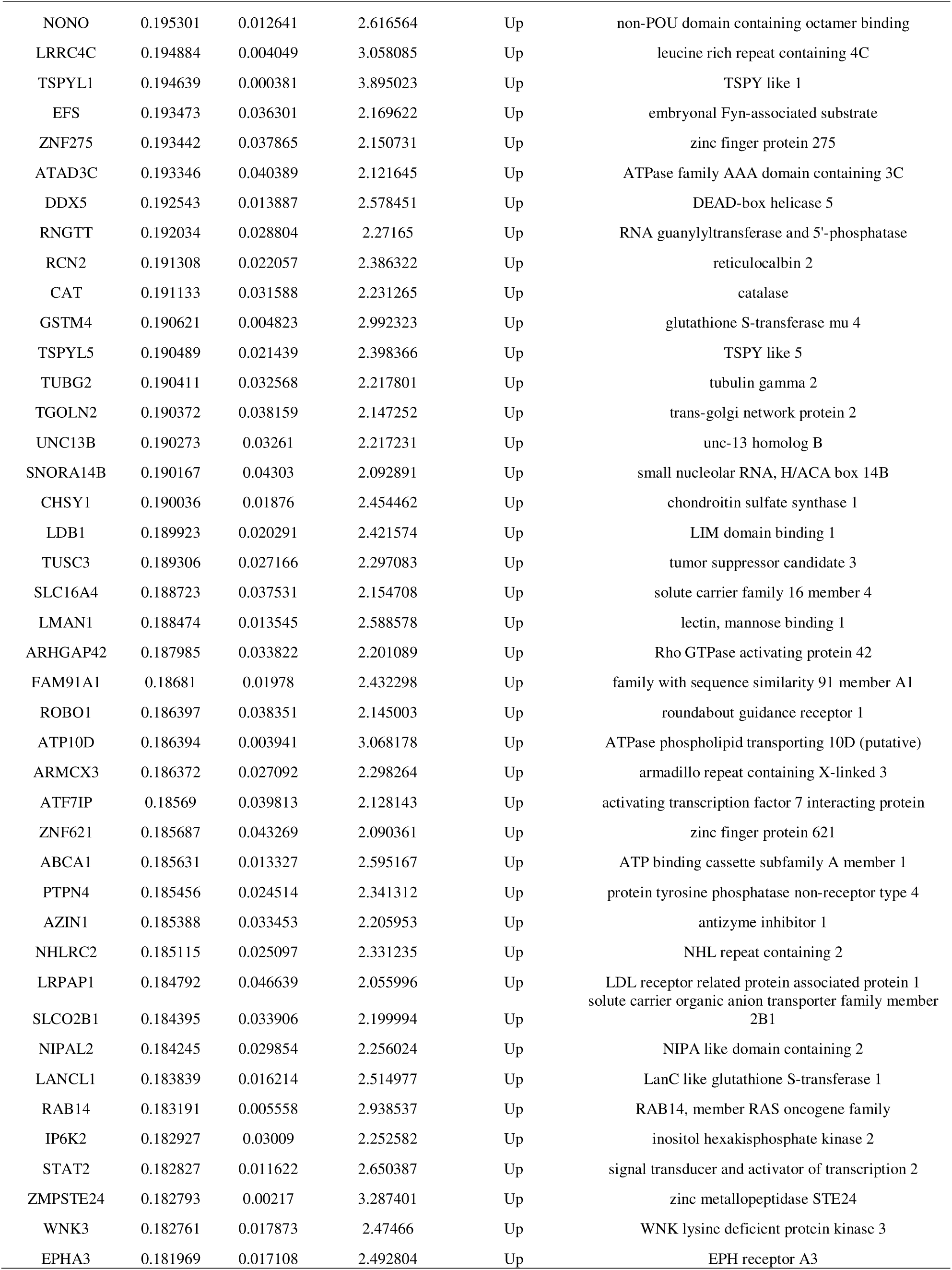
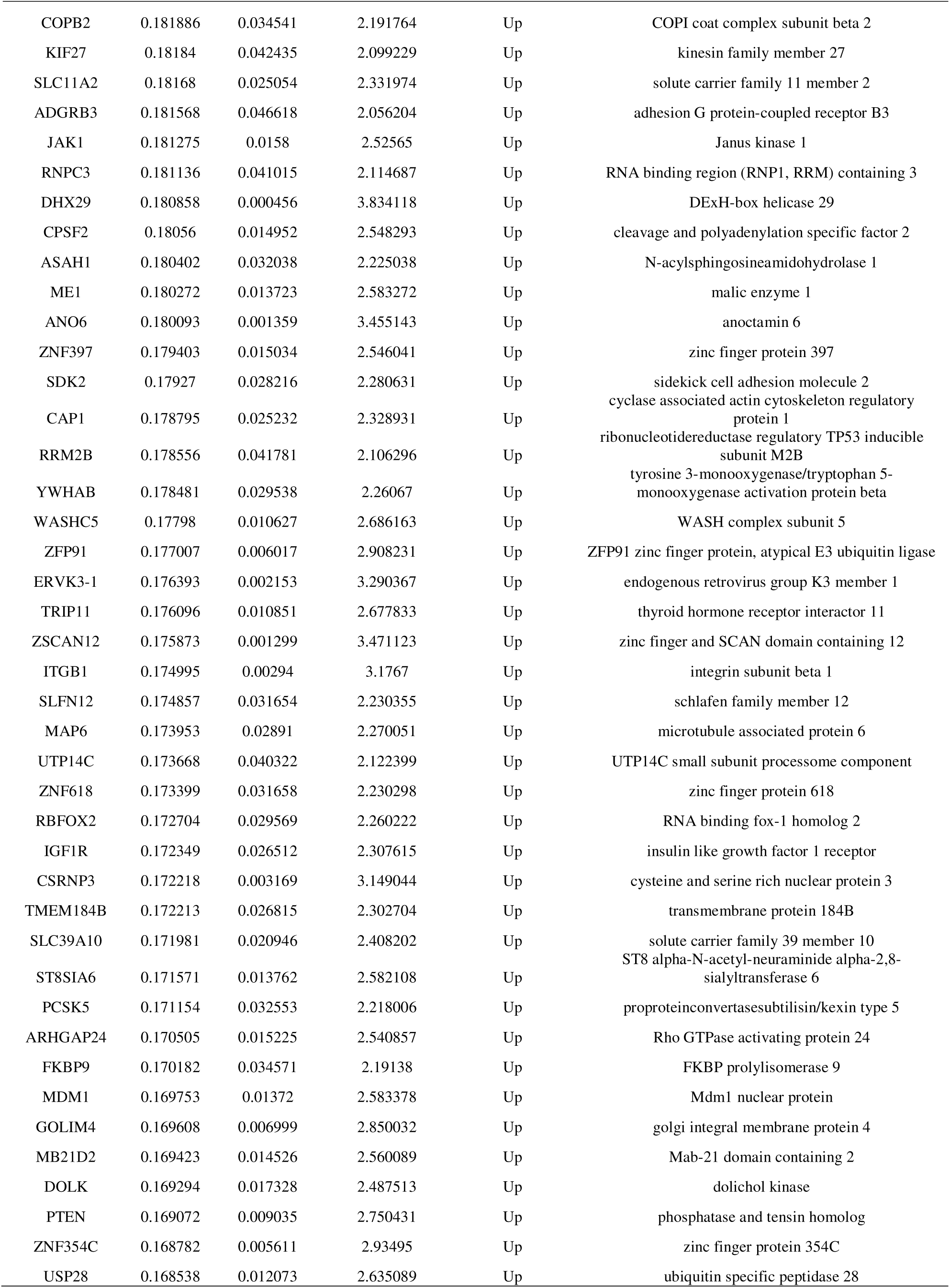
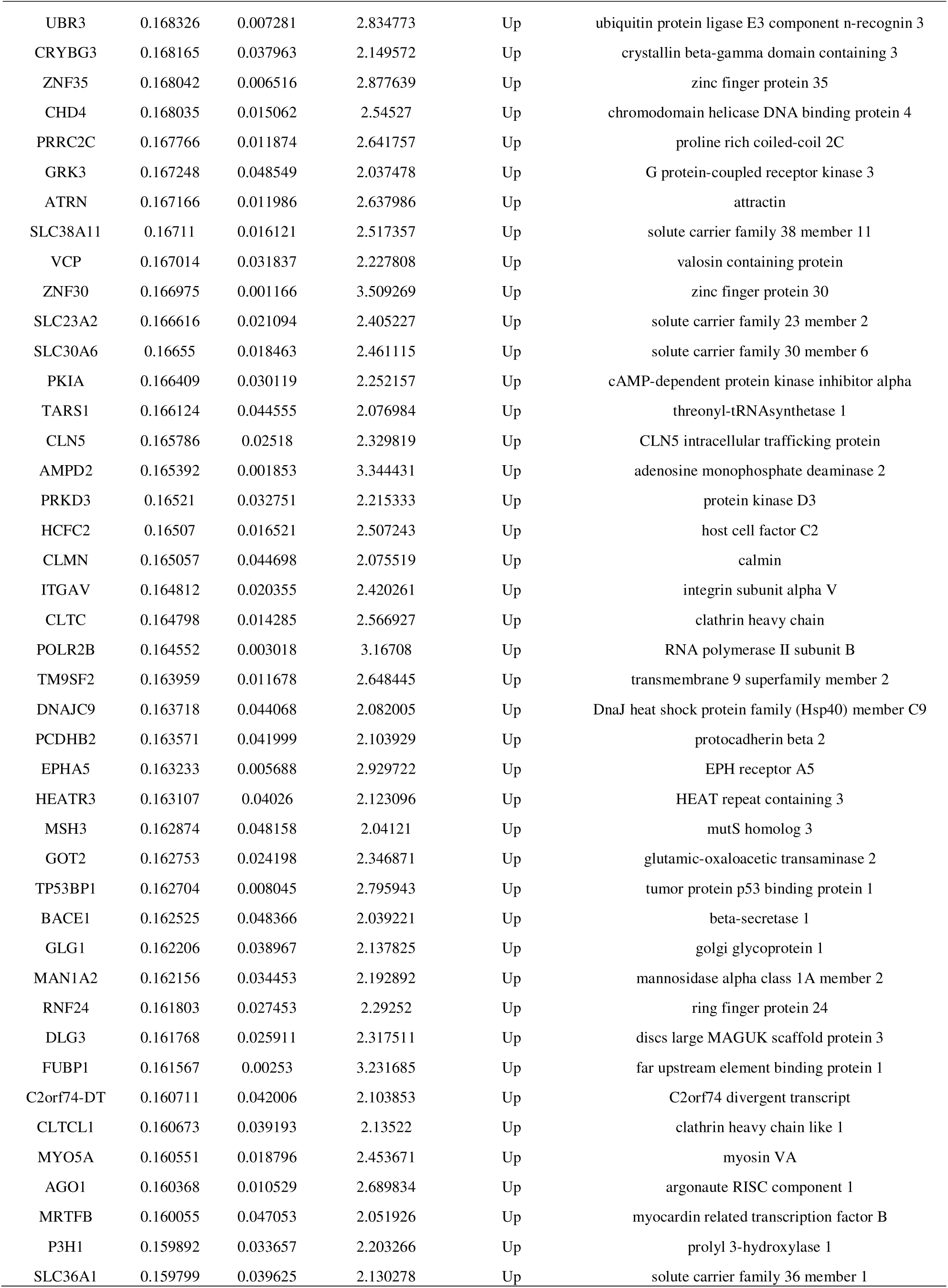
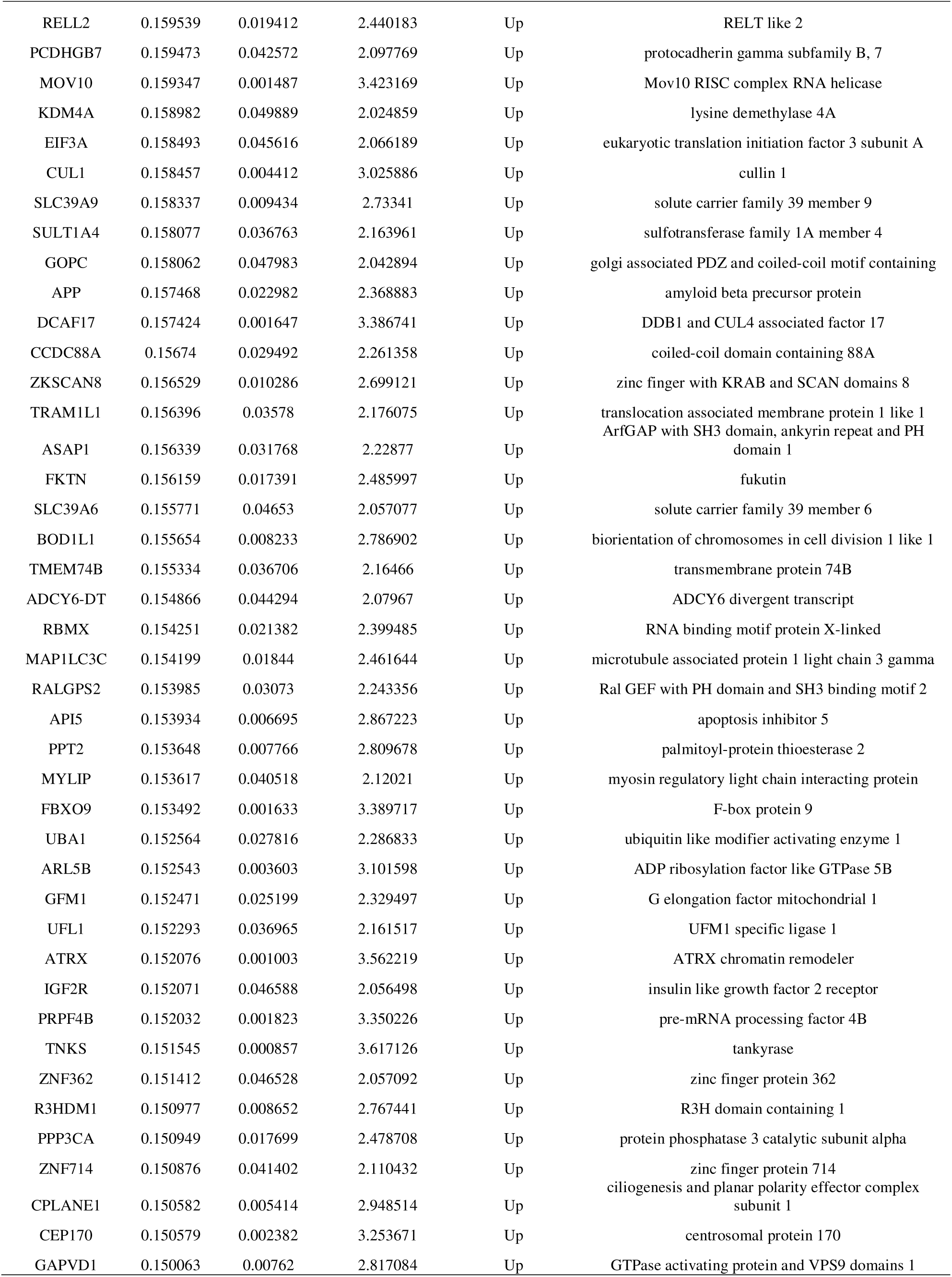
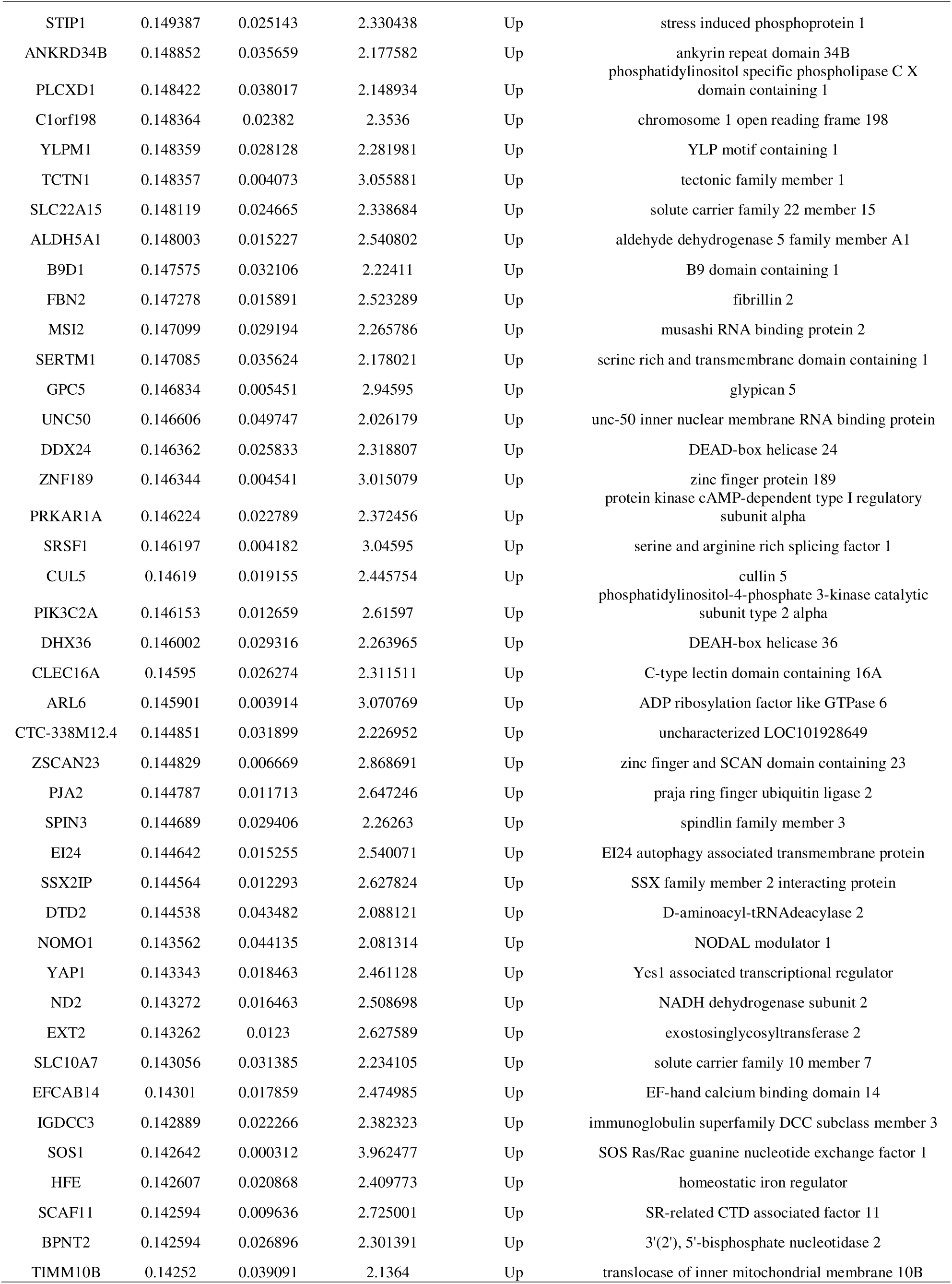
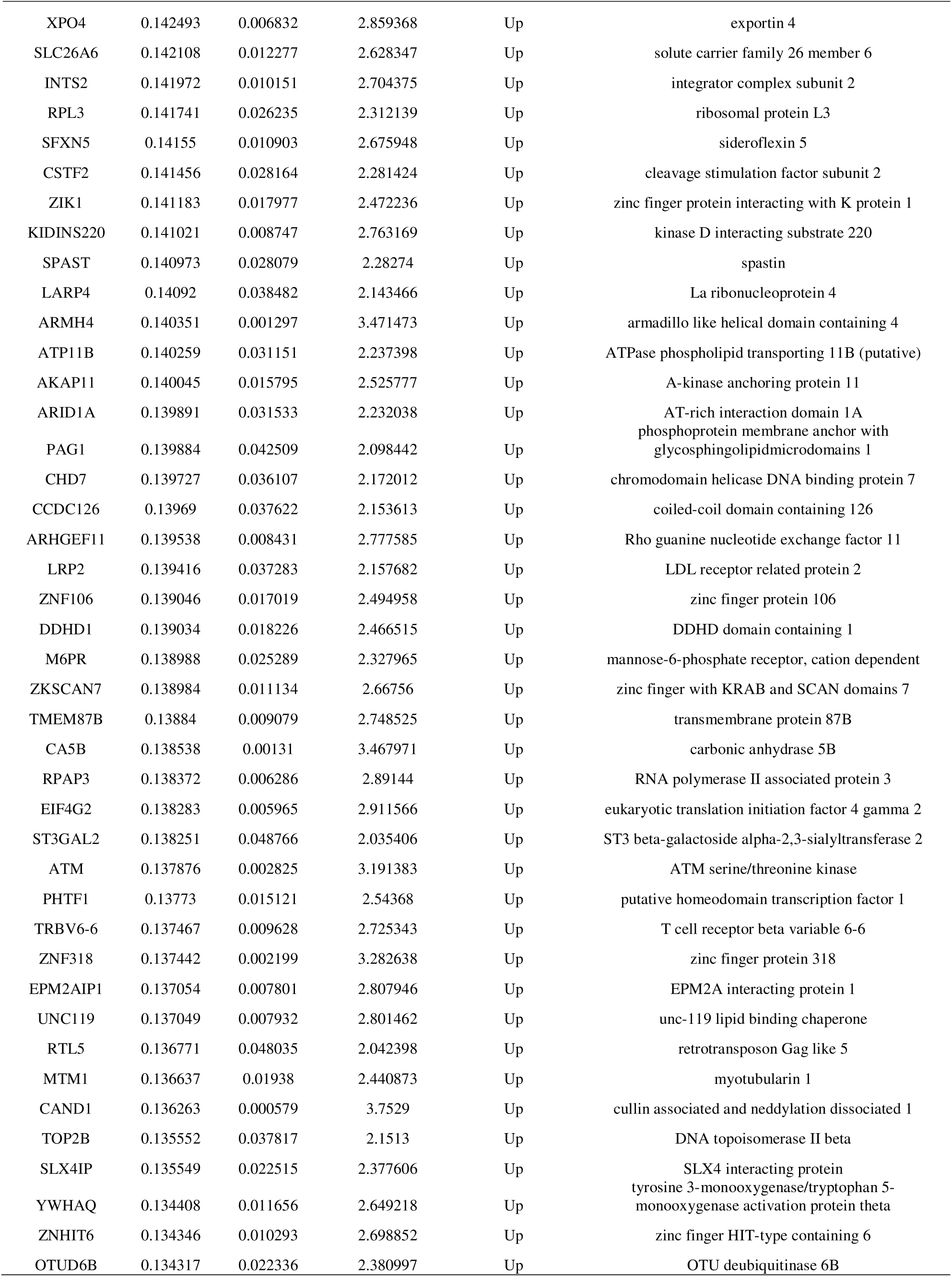
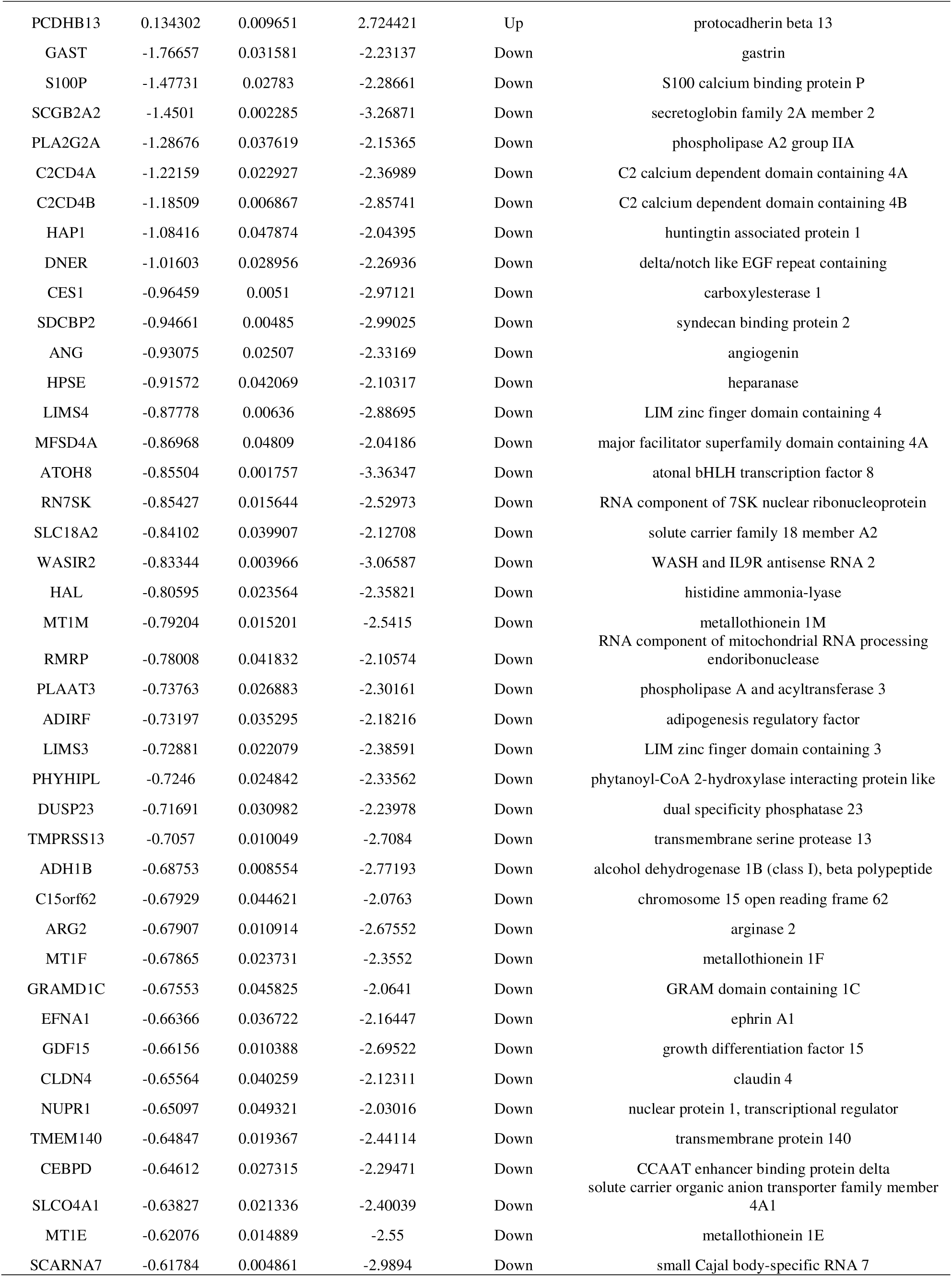
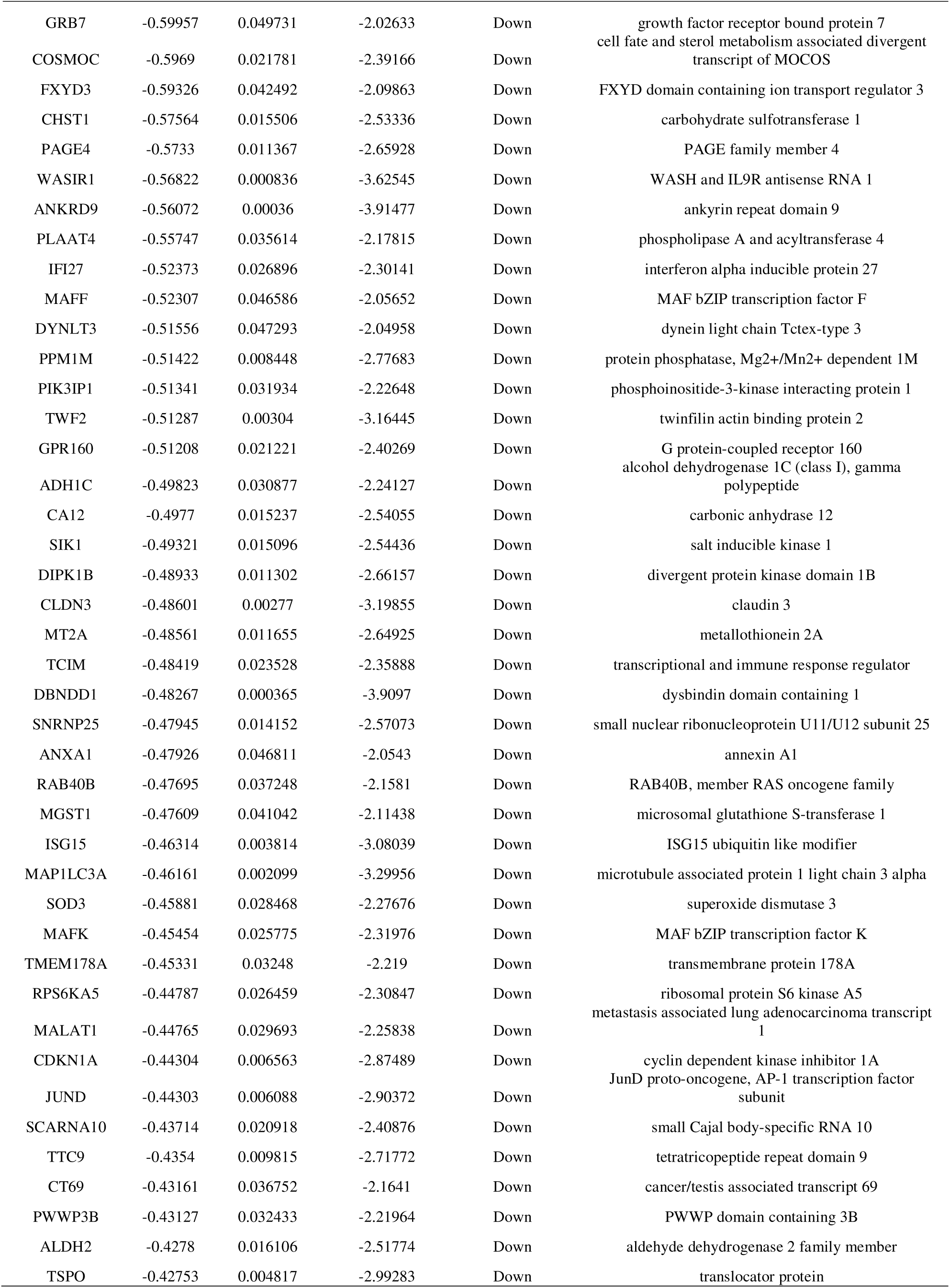
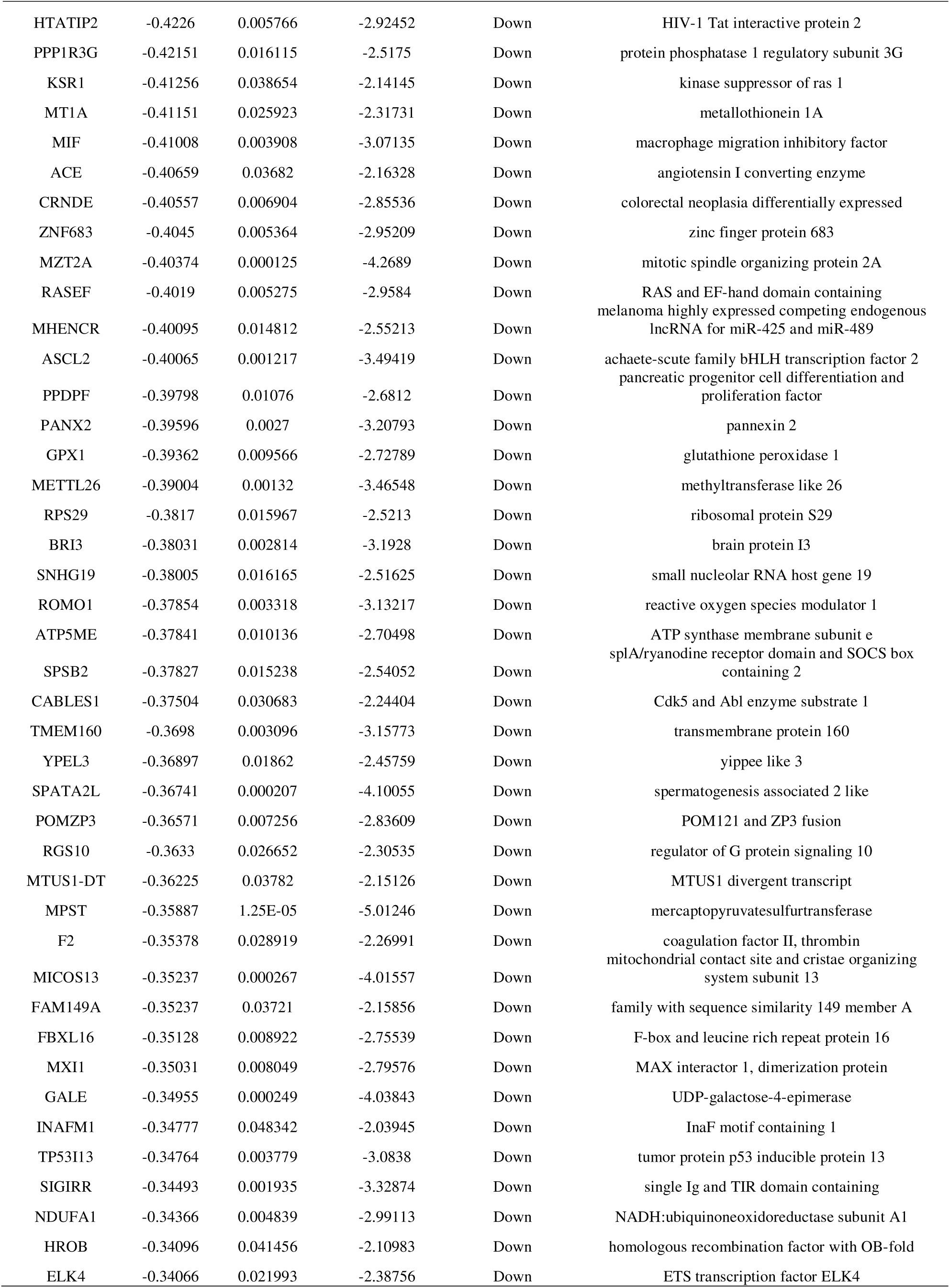
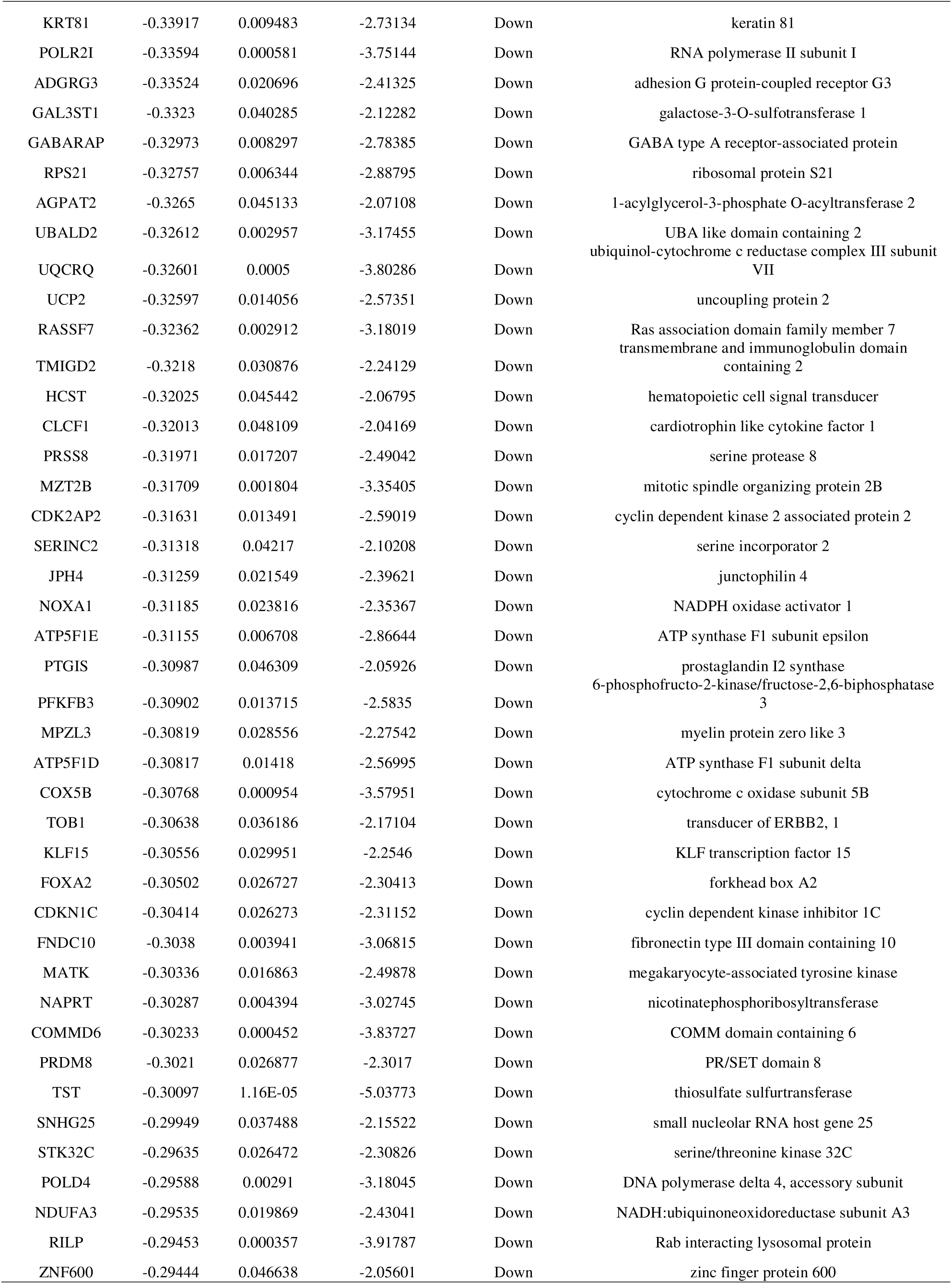
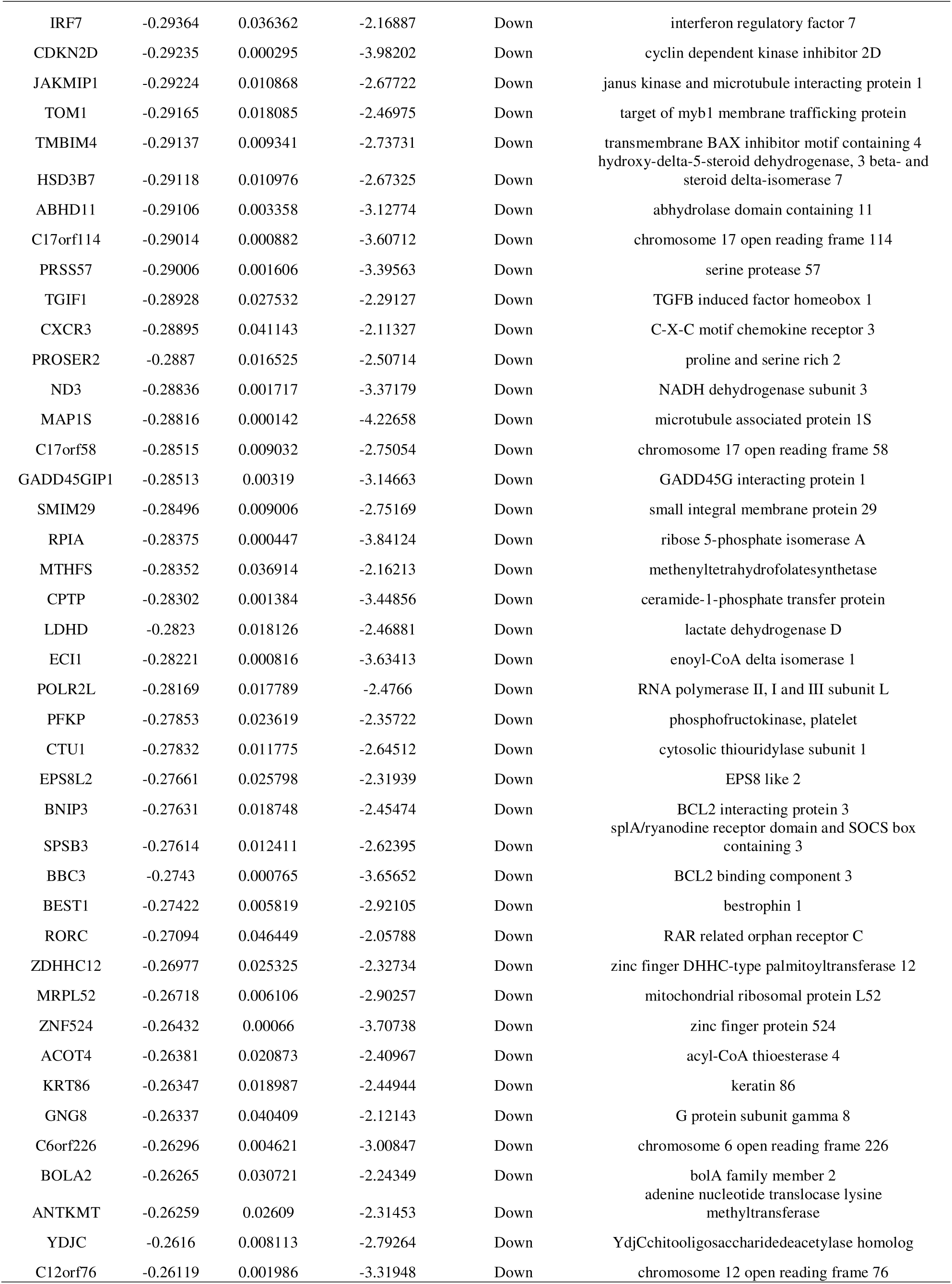
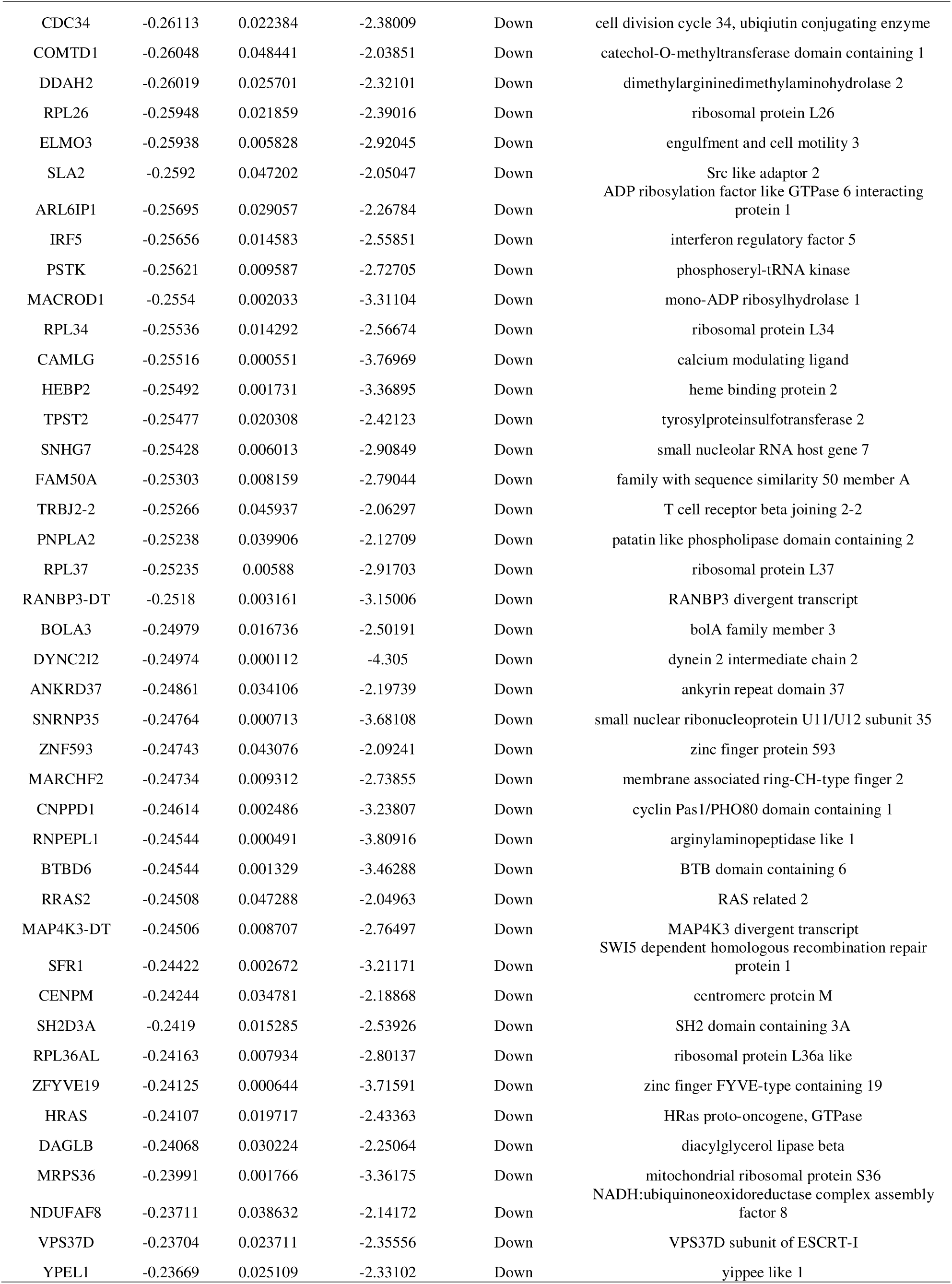
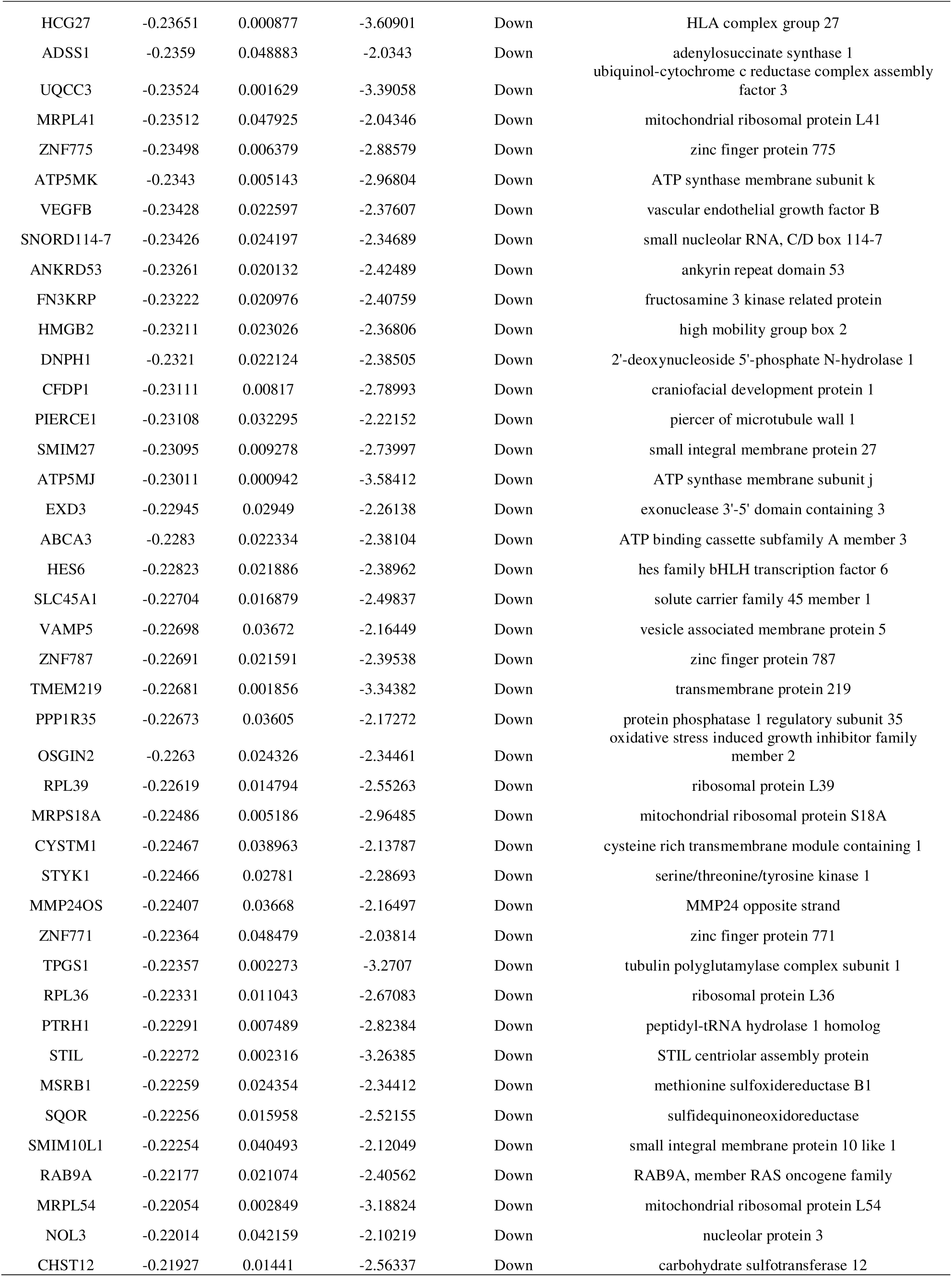
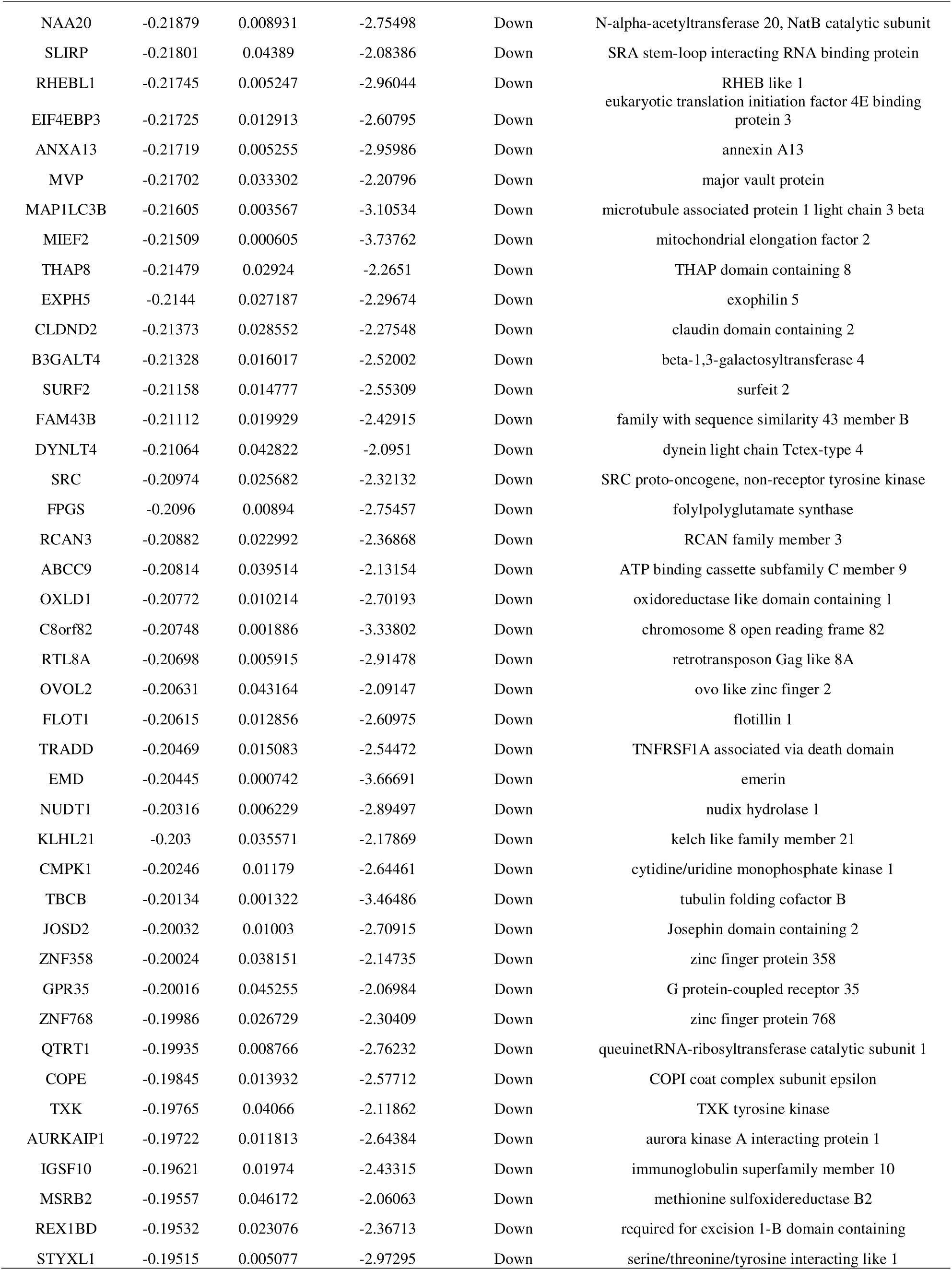
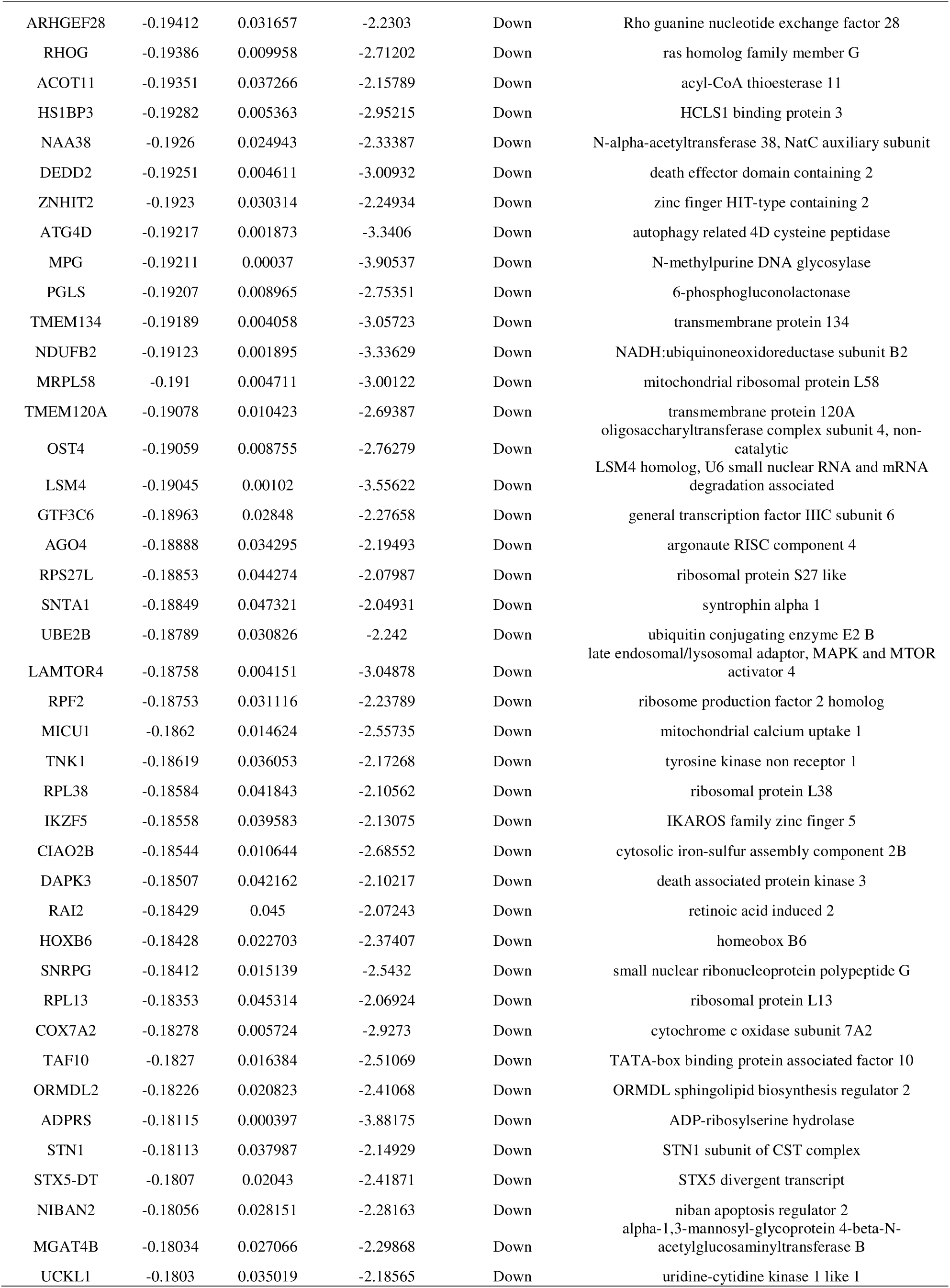
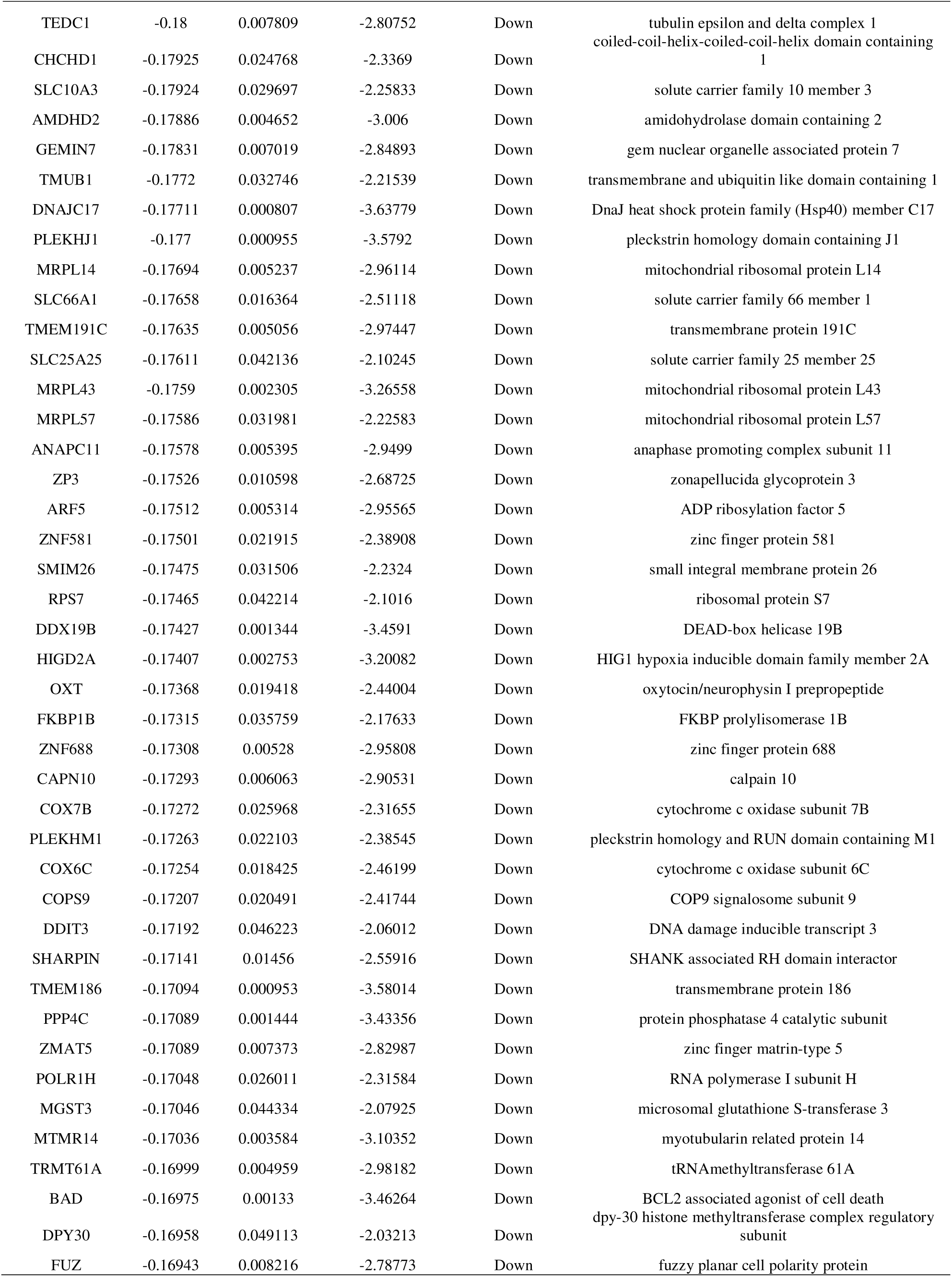
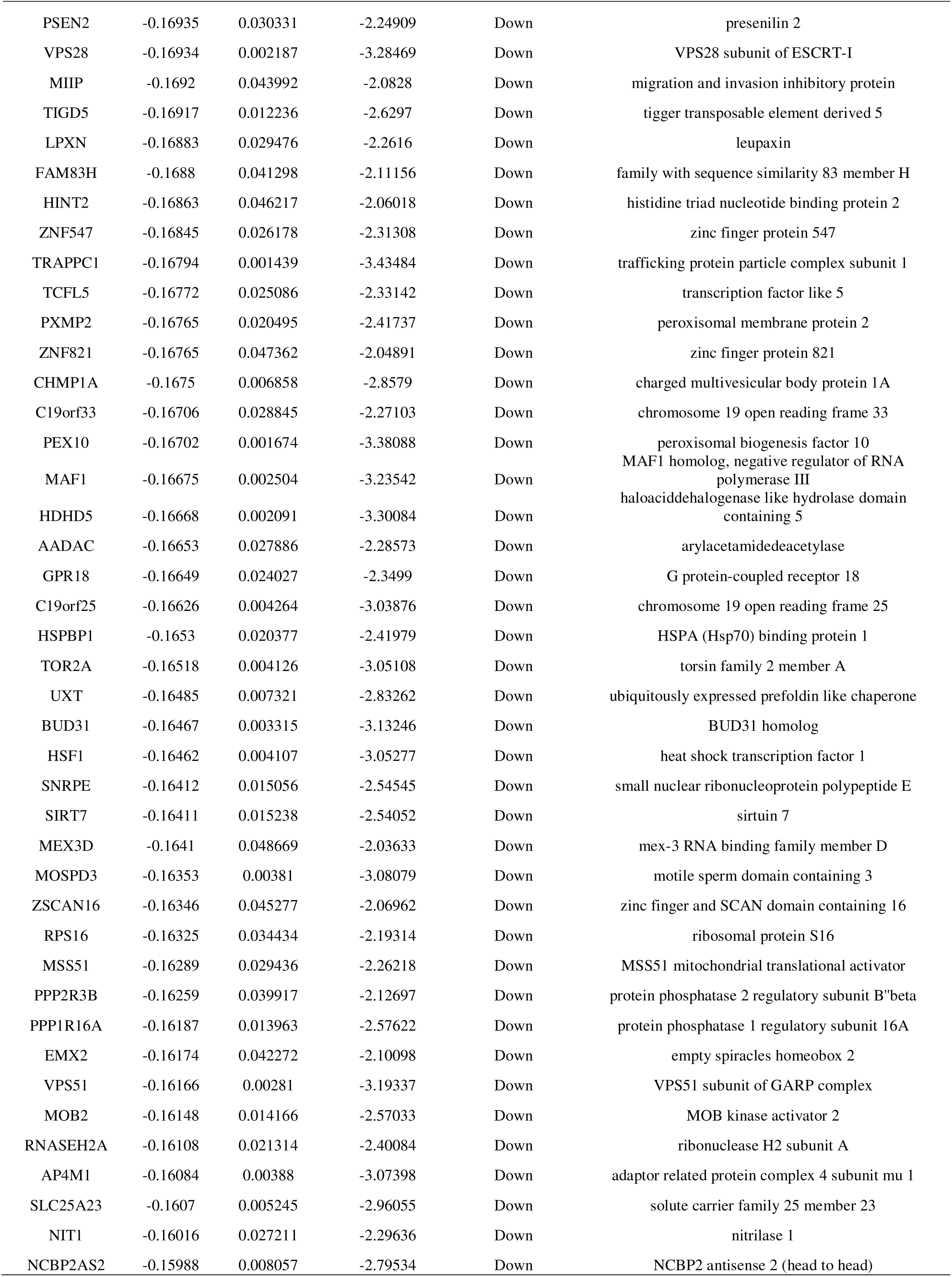
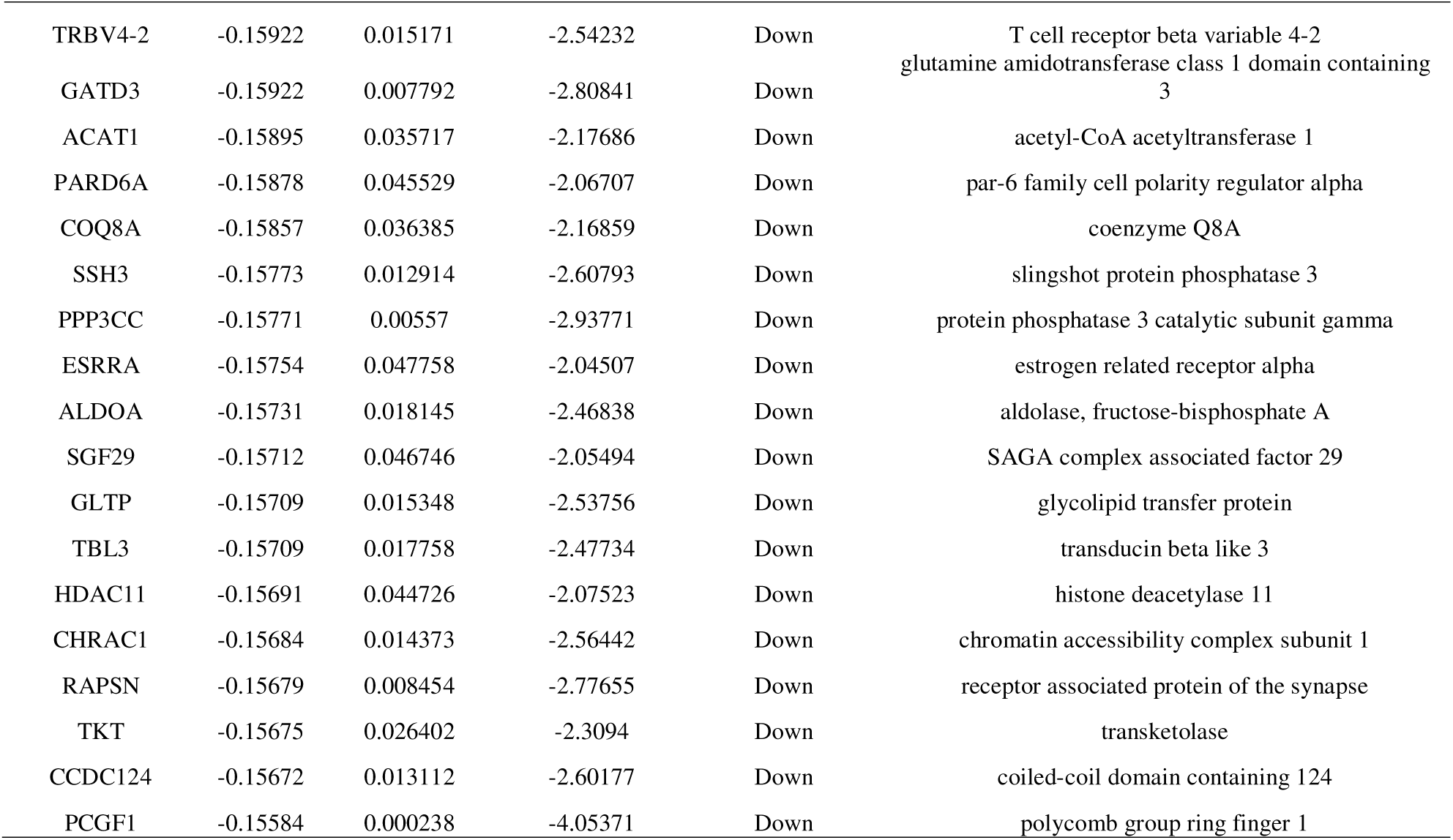
The statistical metrics for key differentially expressed genes (DEGs).

### GO and pathway enrichment analyses of DEGs

GO and REACTOME pathway enrichment analyses were performed to better understand the biological functions of the identified DEGs using the online tool g:Profiler. The top significant GO terms for each of the subsections (BP, CC, and MF) (Table 2) and top REACTOME pathways (Table 3) were clustered with the threshold of adjusted p < 0.05. In the BP, the DEGs were significantly enriched in multicellular organismal process, regulation of biological process, organonitrogen compound metabolic process and metabolic process. Under the term of CC, DEGs were principally associated with the membrane, cytoplasm, intracellular anatomical structure and cytosol. Among the MF, DEGs were mainly enriched in small molecule binding, carbohydrate derivative binding, catalytic activity and hydrolase activity. Additionally, DEGs were significantly enriched in REACTOME pathways, including extracellular matrix organization, RHO GTPase cycle, translation and Respiratory electron transport, ATP synthesis by chemiosmotic coupling, and heat production by uncoupling proteins.

**Table 2.**
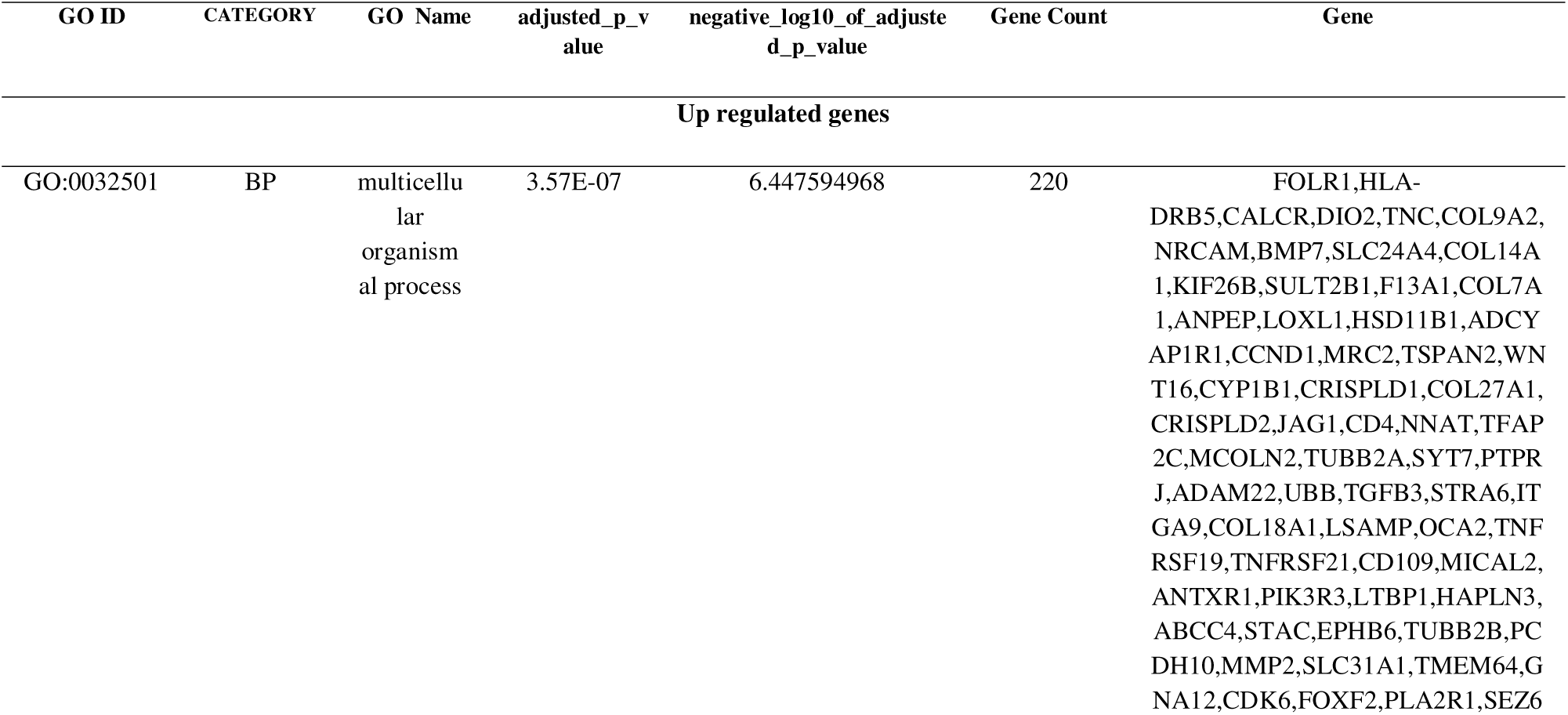
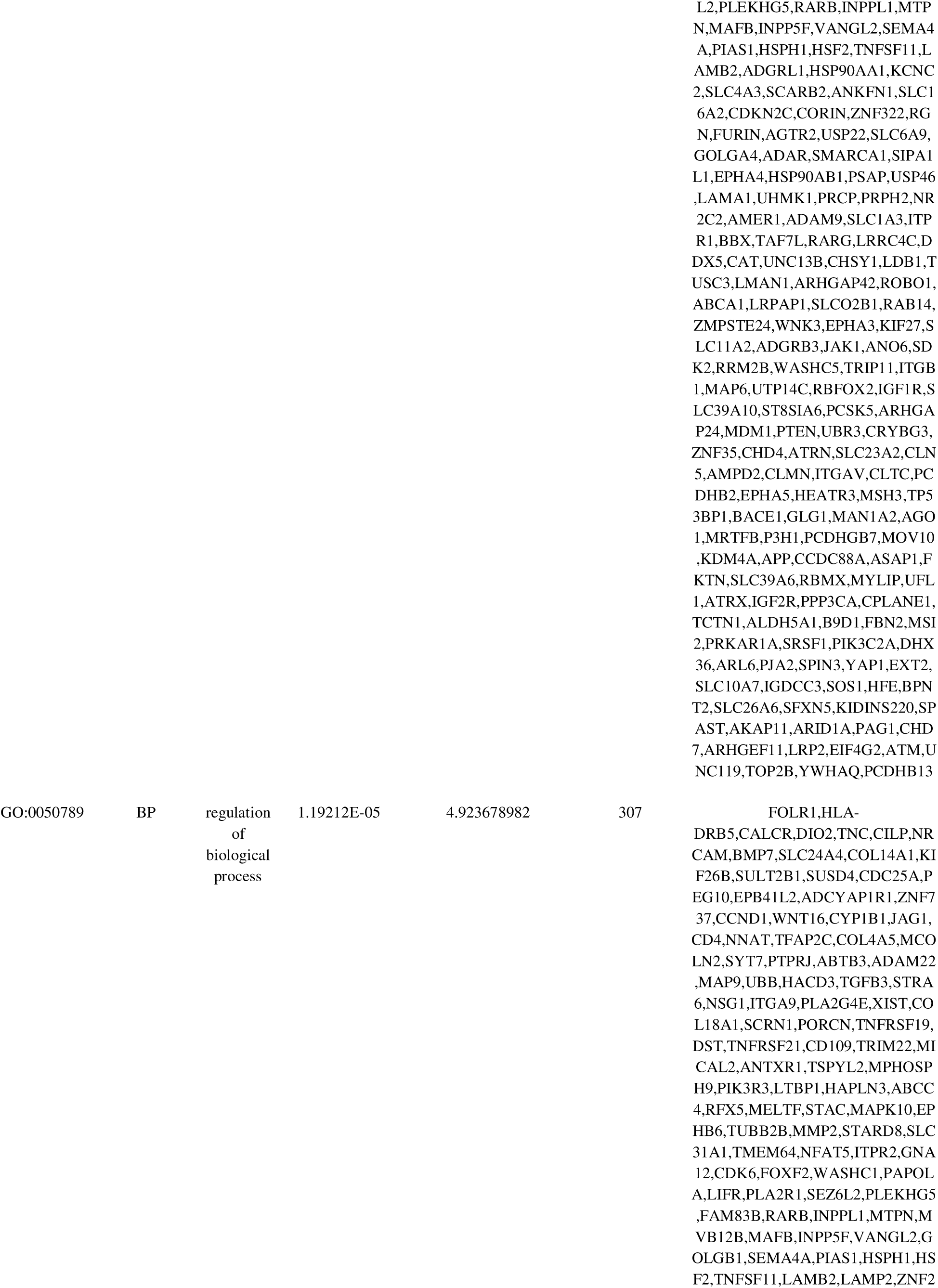
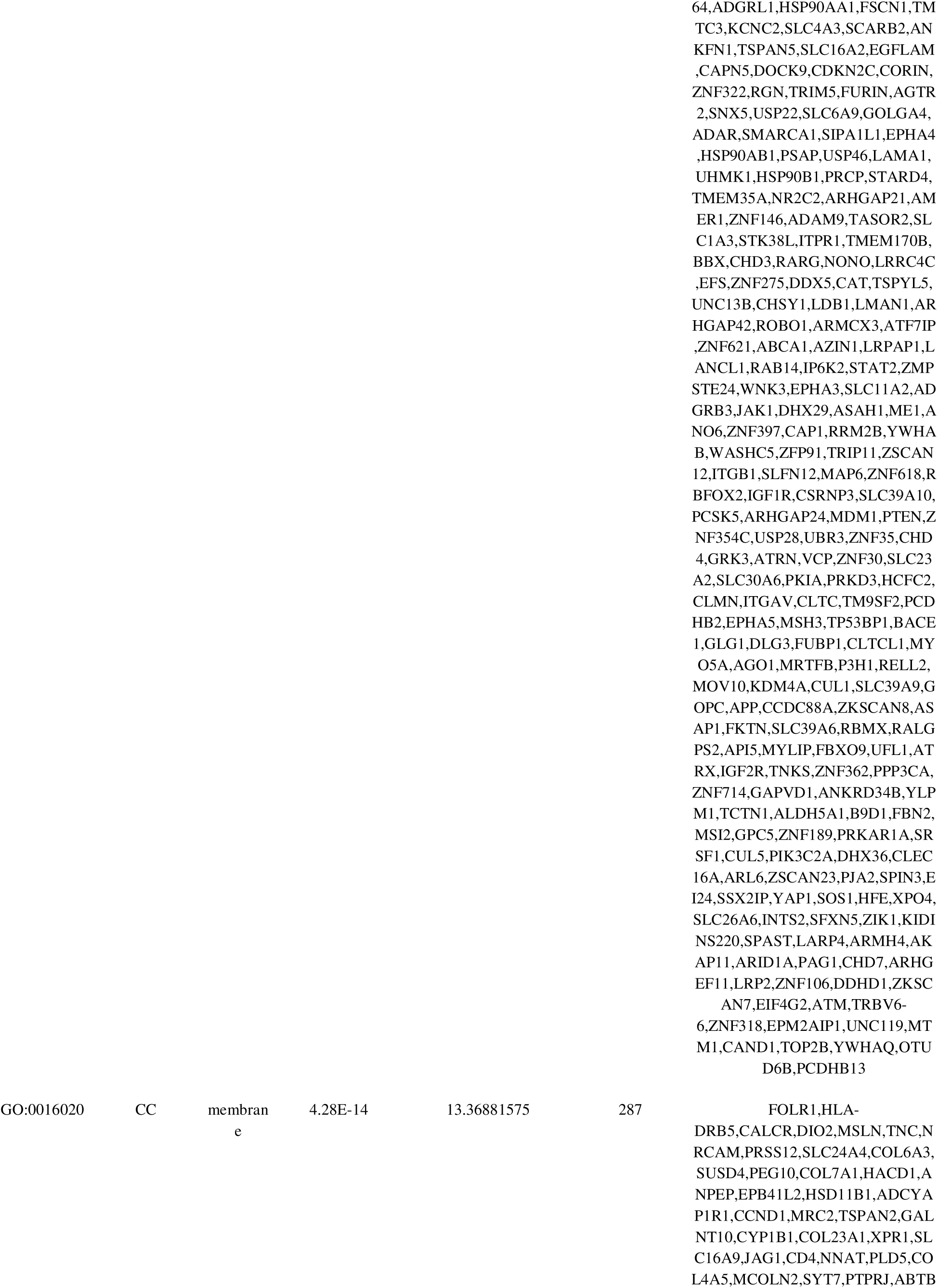
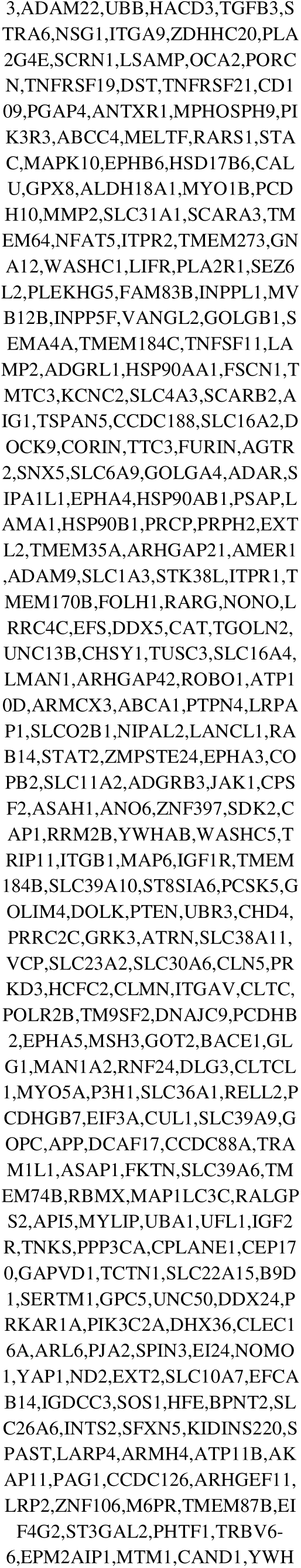
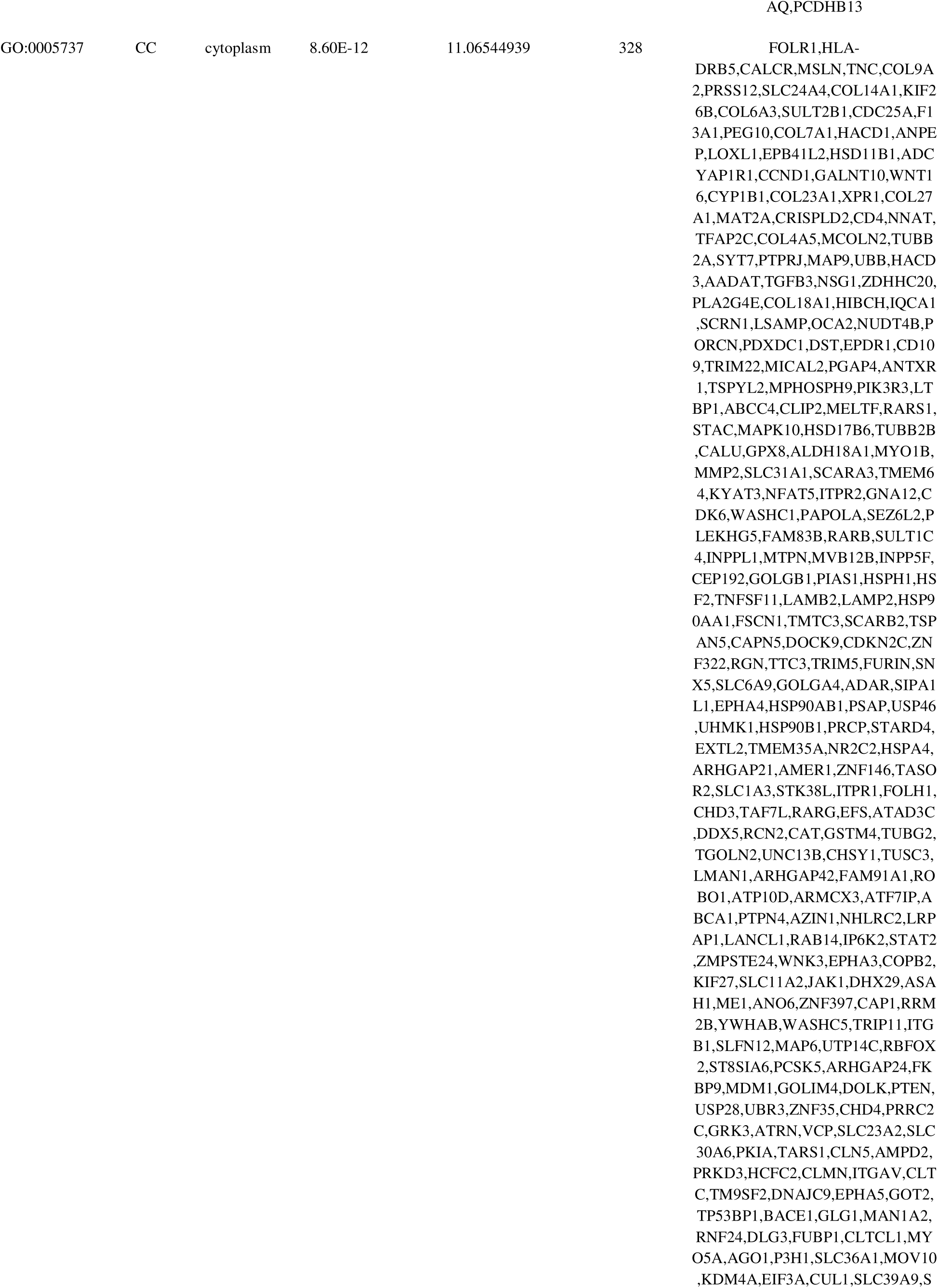
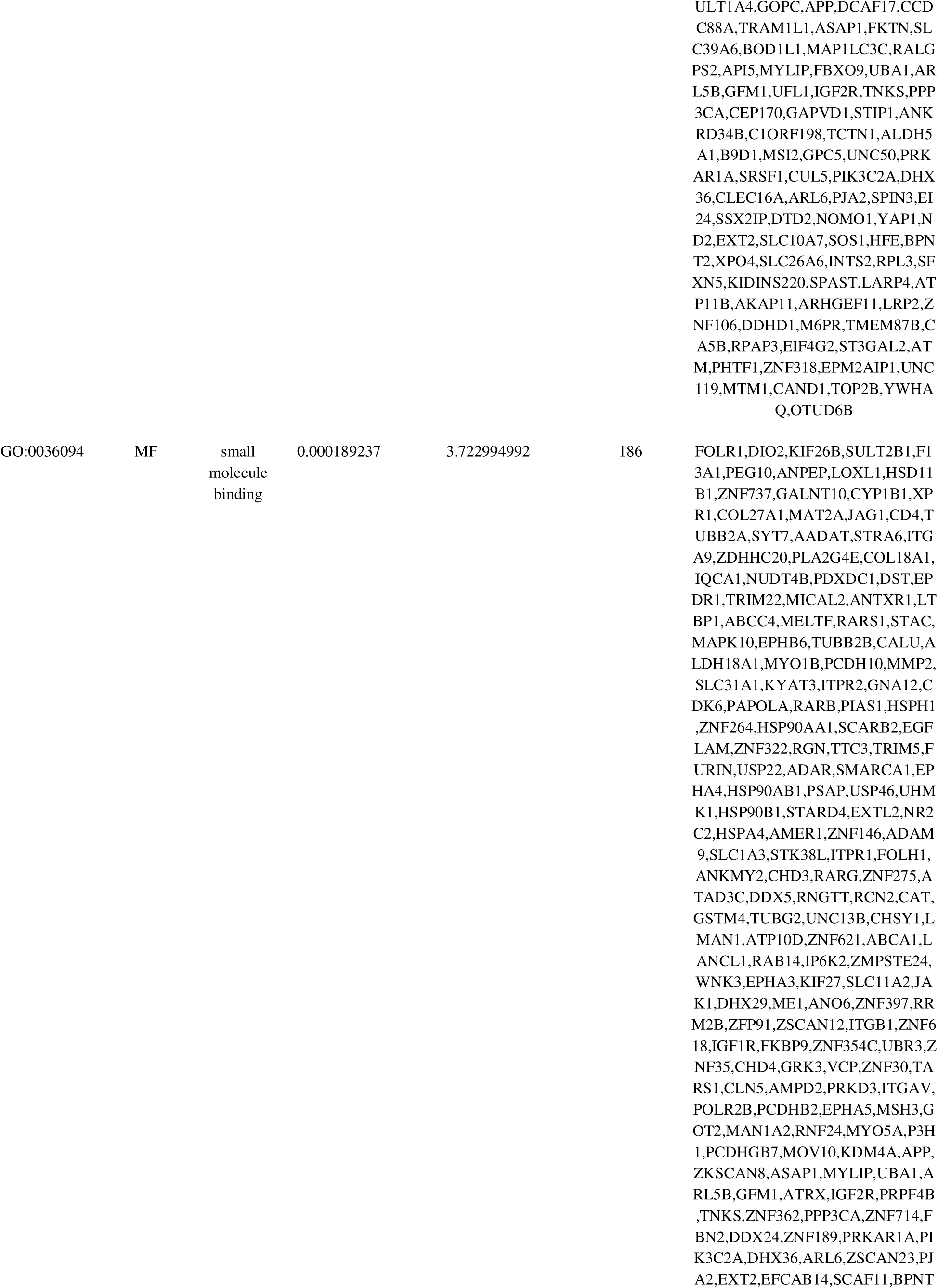
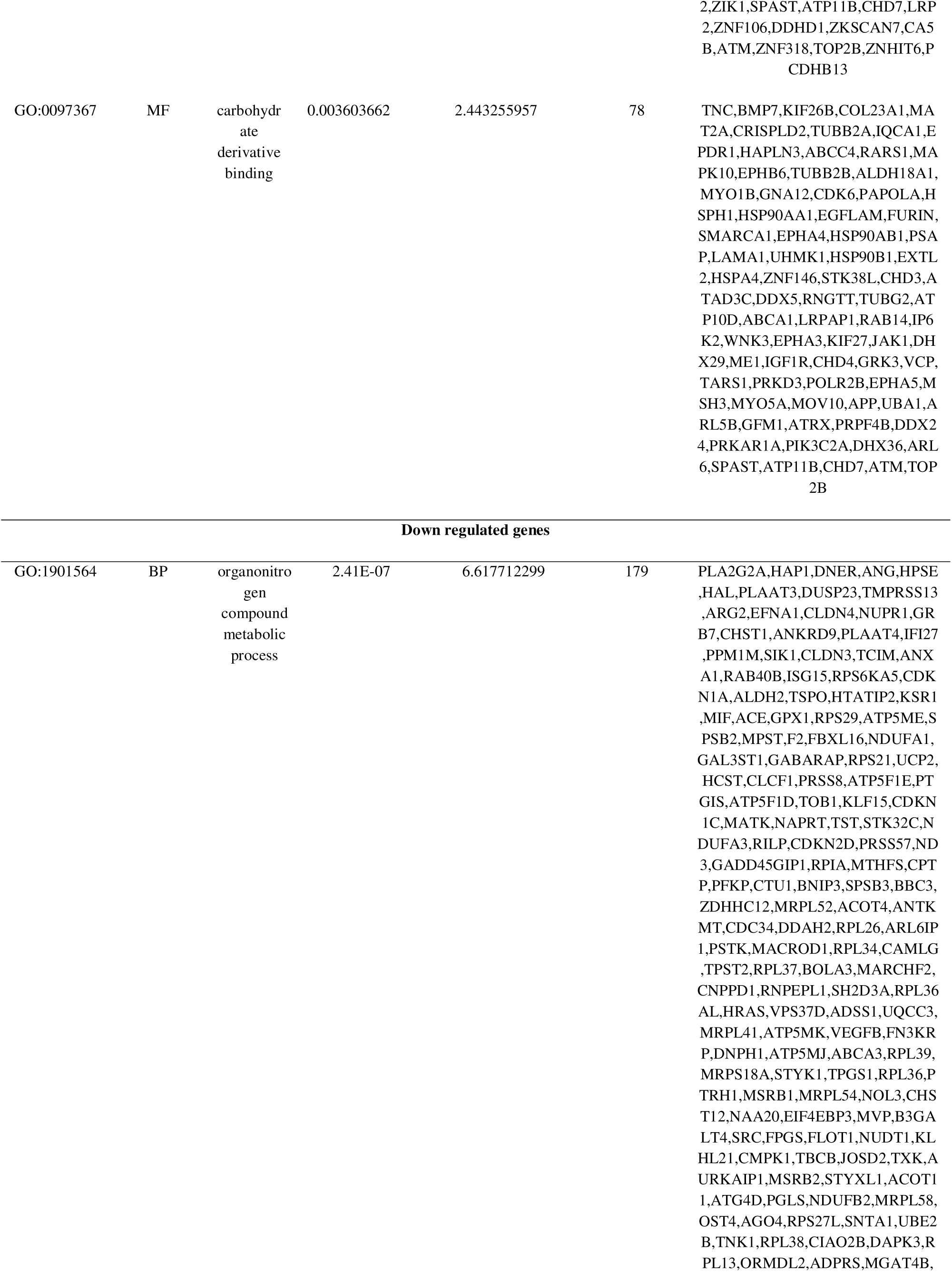
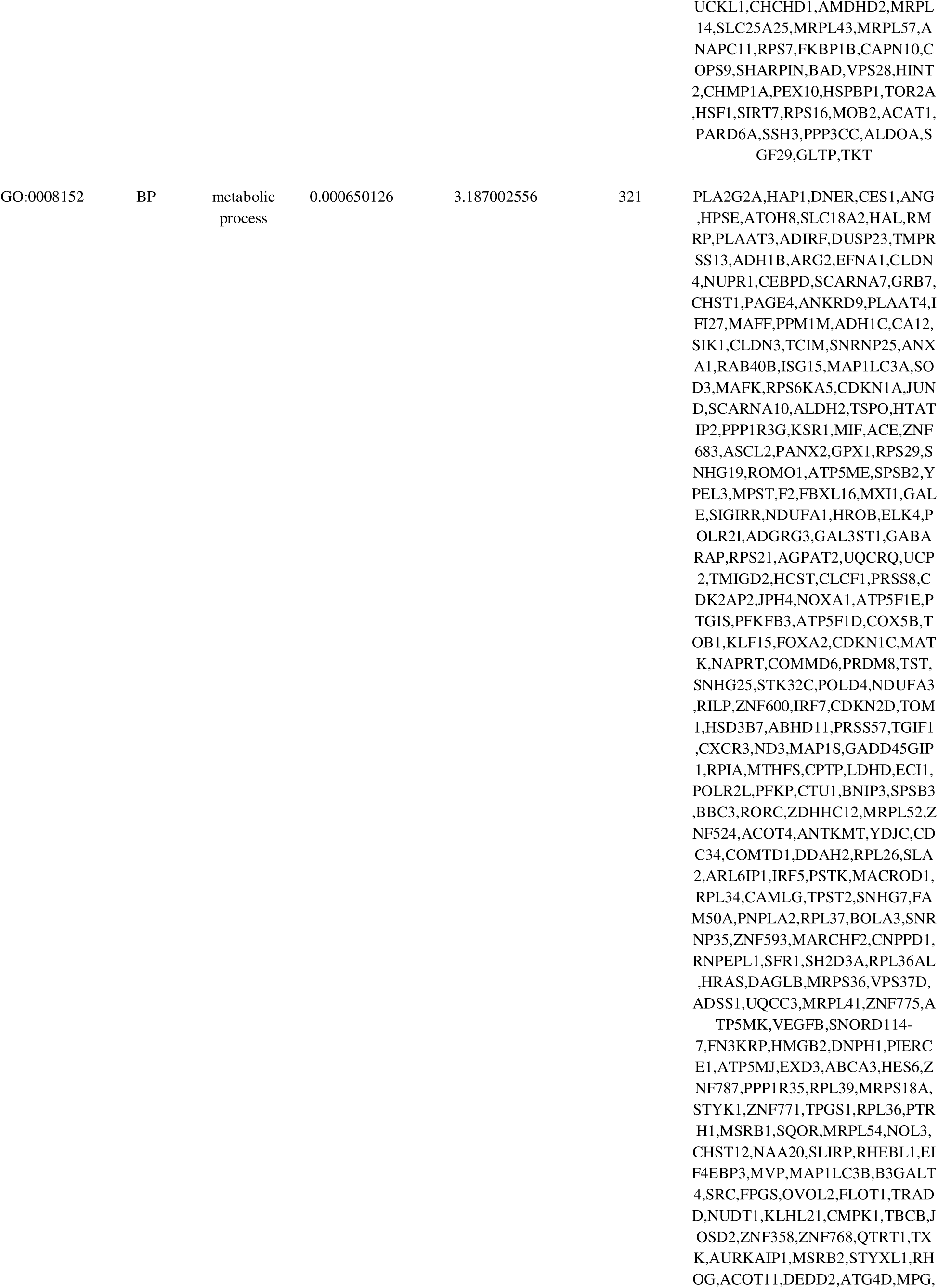
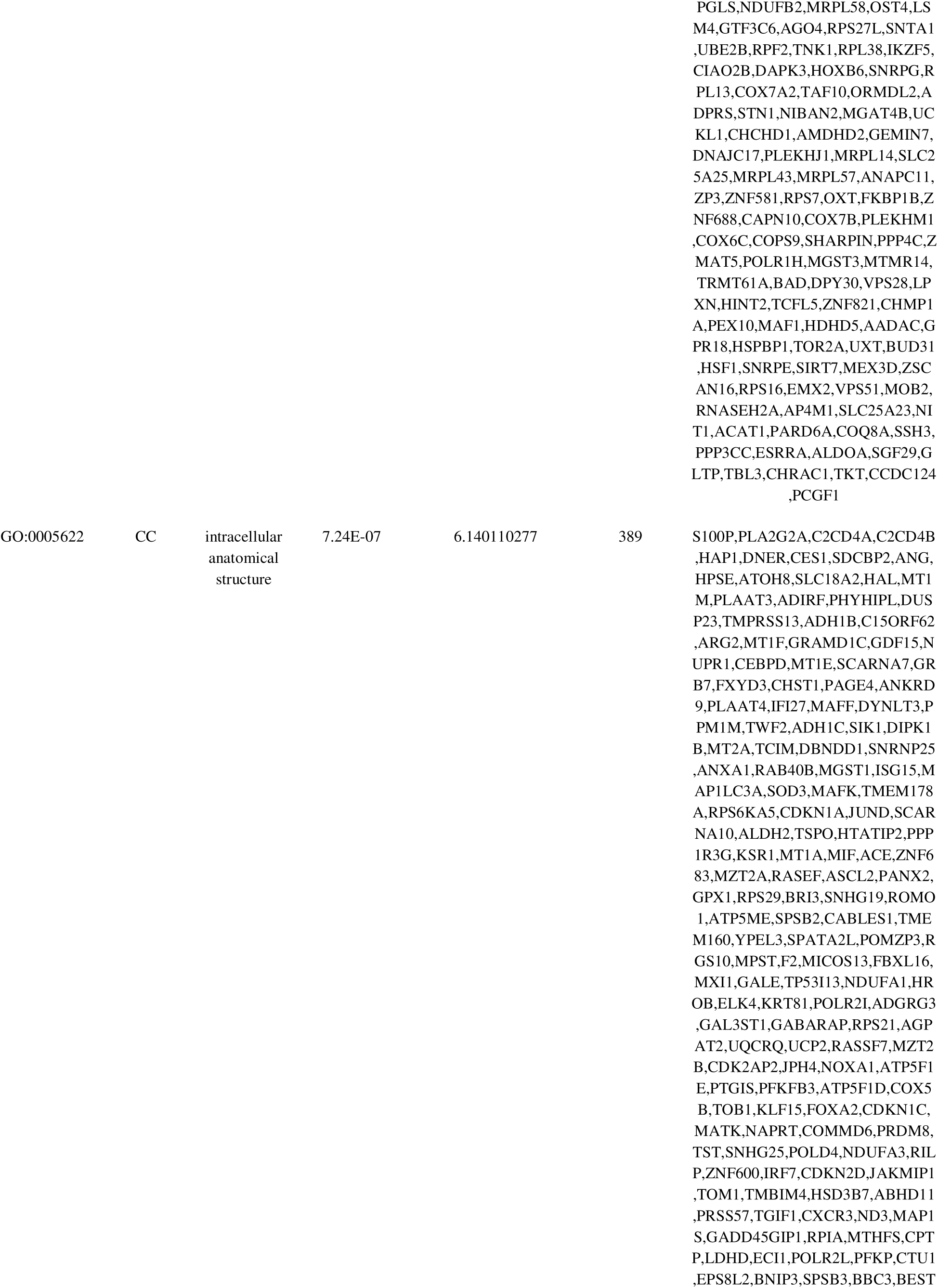
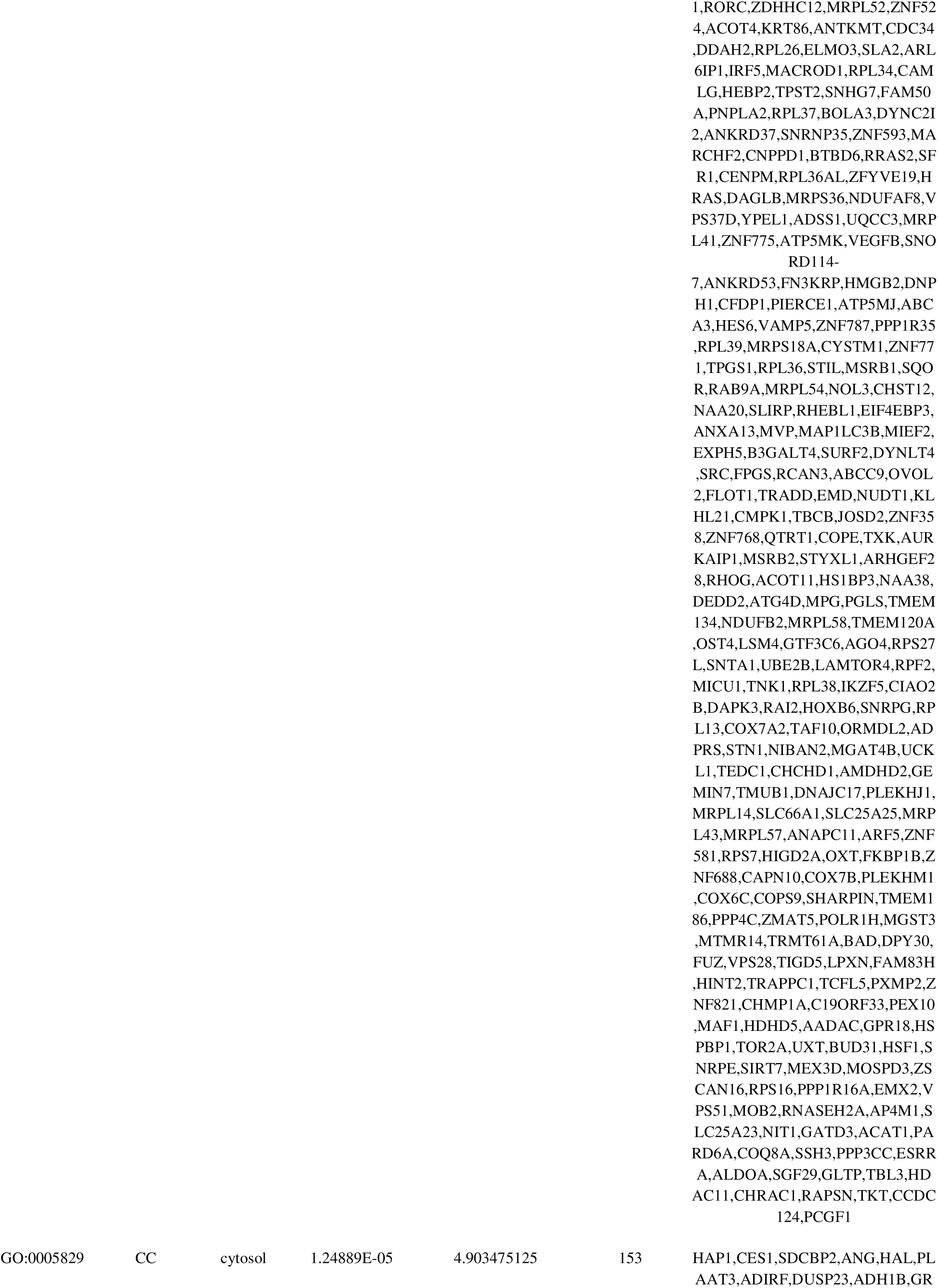
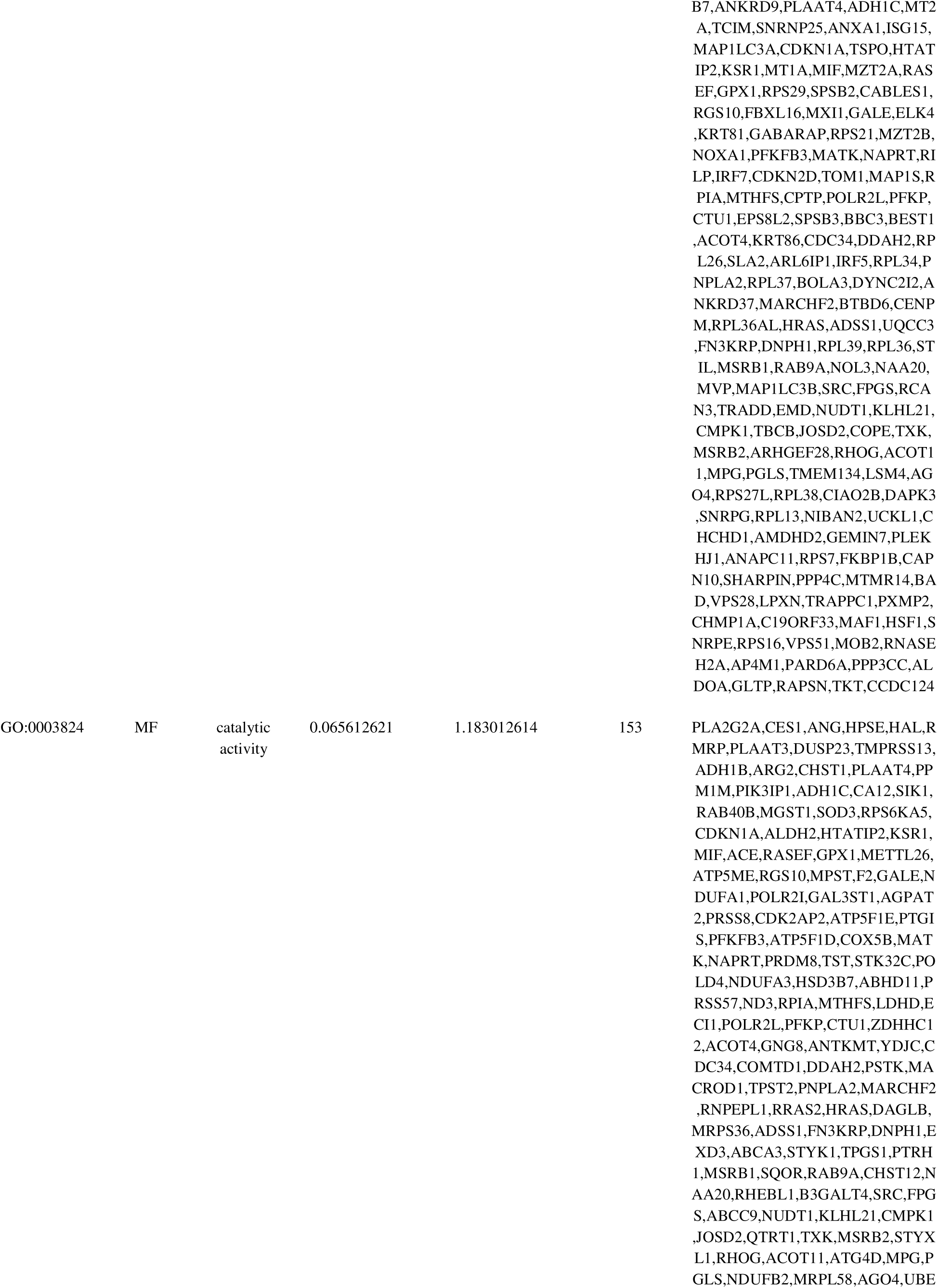
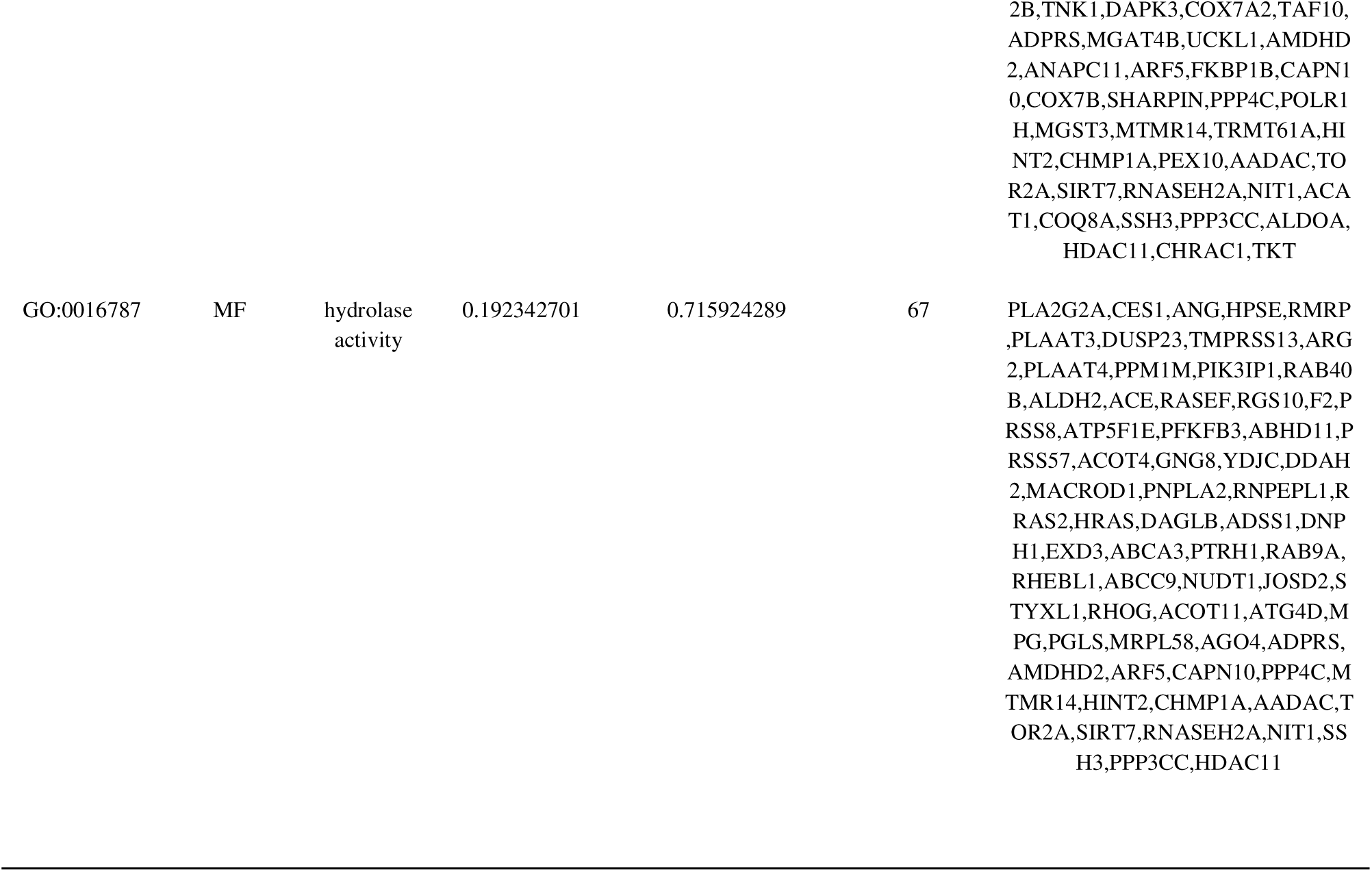
The enriched GO terms of the up and down regulated differentially expressed genes.

**Table 3.**
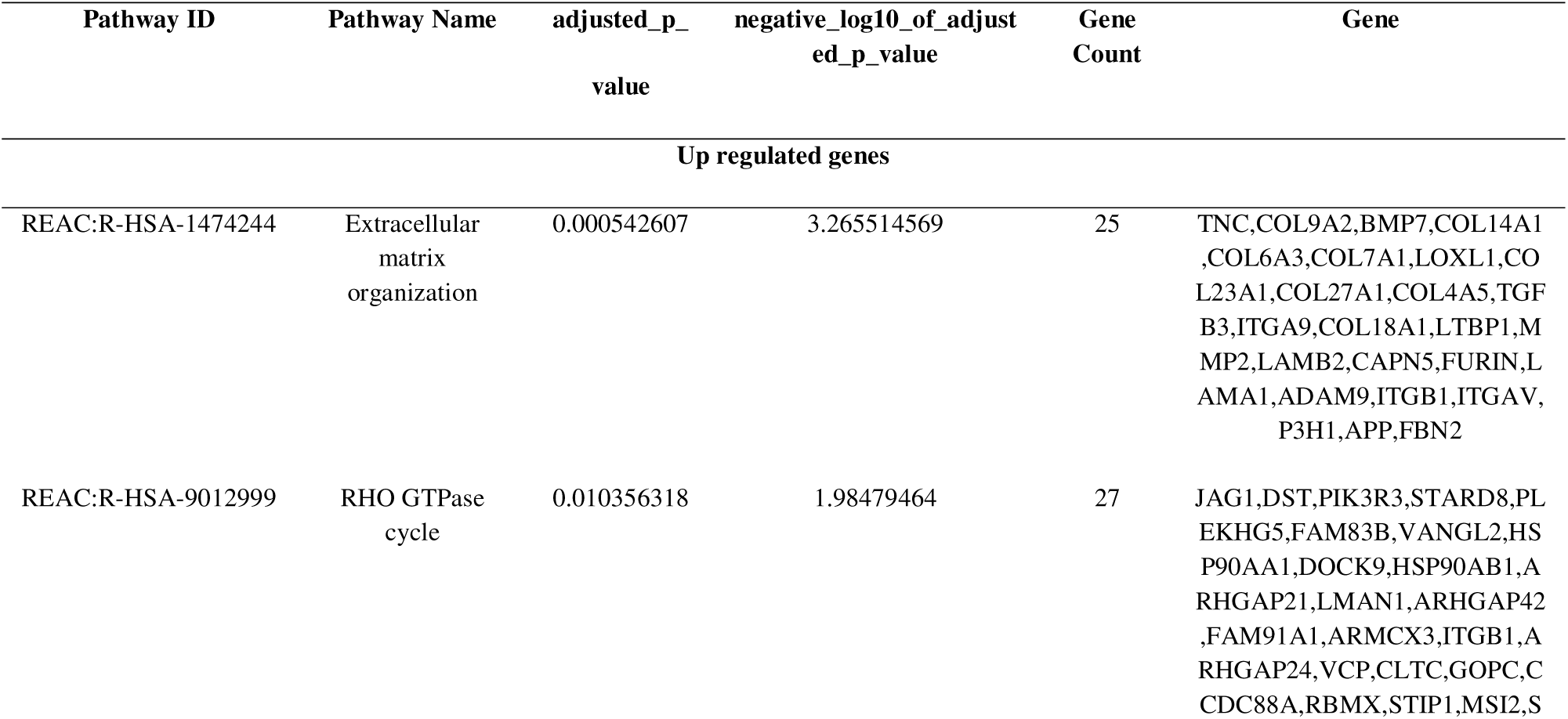
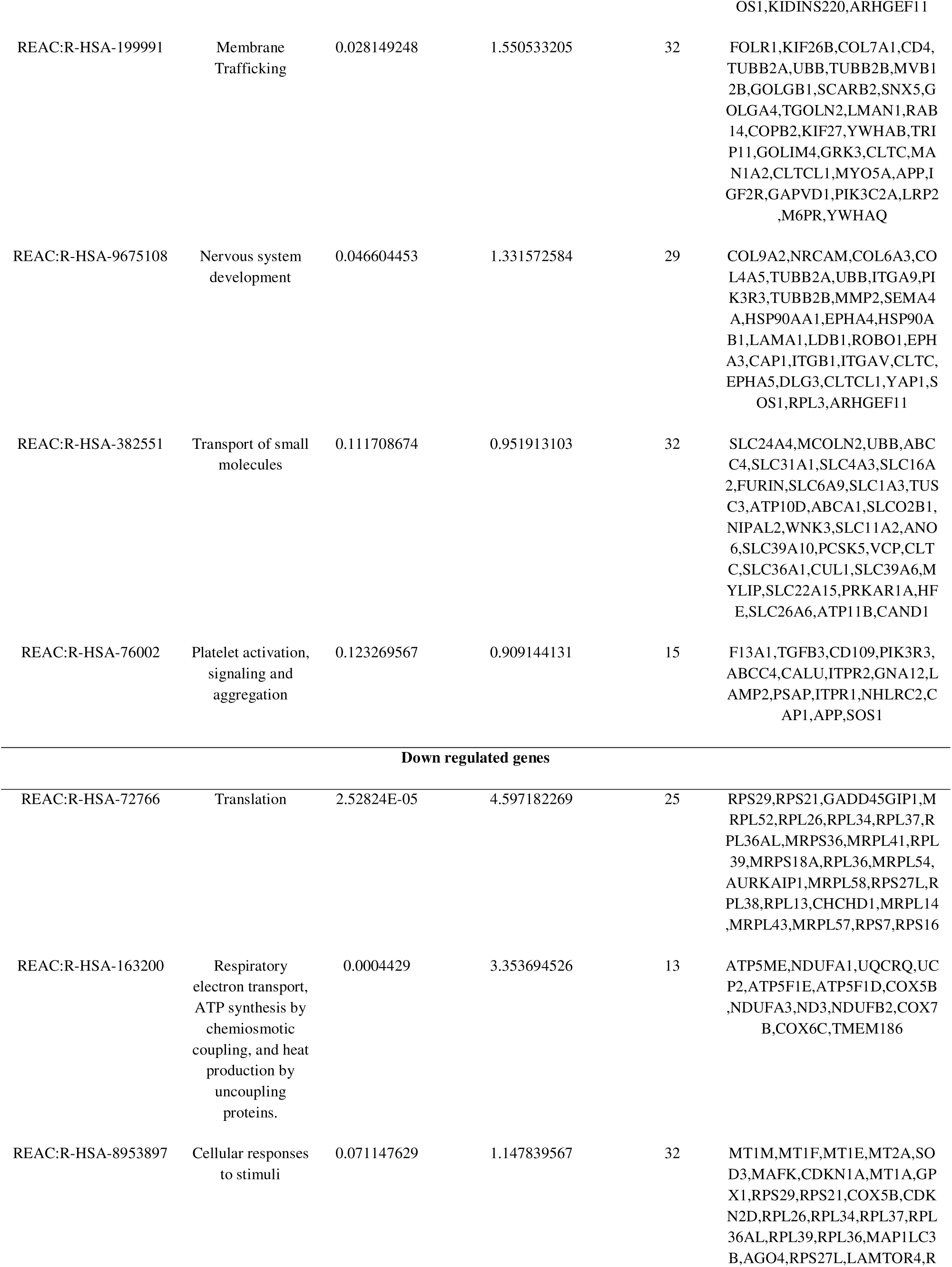
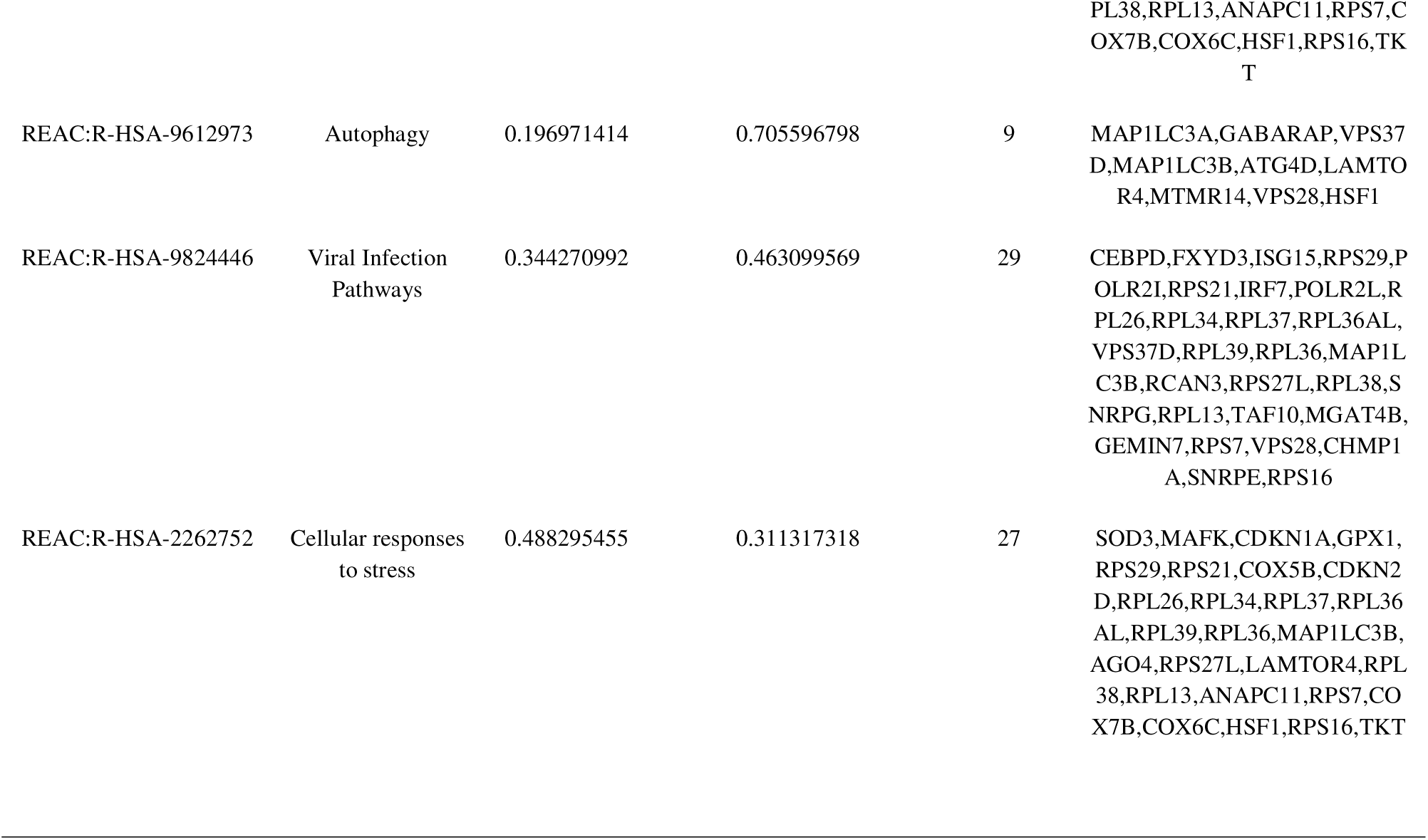
The enriched pathway terms of the up and down regulated differentially expressed genes.

### Construction of the PPI network and module analysis

The PPI network of DEGs was established by the IMex interactome database to analyze the interaction between the DEGs and visualized by Cytoscape software (Fig.3). PPI network consist of 6316 nodes and 15825 edges. To identify the hub genes from the PPI network, the Network Analyzer plug in of Cytoscape software was used. And according to the node degree, betweenness, stress and closeness algorithms, we selected the top hub genes as potential hub genes (Table 4), including APP, HSP90AA1, CAND1, CUL1, HSP90AB1, SIRT7, SRC, CDKN1A, ISG15 and RPS16. Two significant modules were identified by the PEWCC. A total of 44 nodes and 177 edges were included in Module 1 (Fig. 4A), and a total of 40 nodes and 141 edges were included in Module 2 (Fig. 4B). Module 1 was mainly enriched in regulation of biological process, membrane, membrane trafficking, RHO GTPase cycle, cytoplasm, small molecules, nervous system development, multicellular organismal process and transport of small molecules. Module 2 was mainly enriched in translation, cellular responses to stimuli, organonitrogen compound metabolic process, viral infection pathways and cellular responses to stress.

**Fig. 3.**
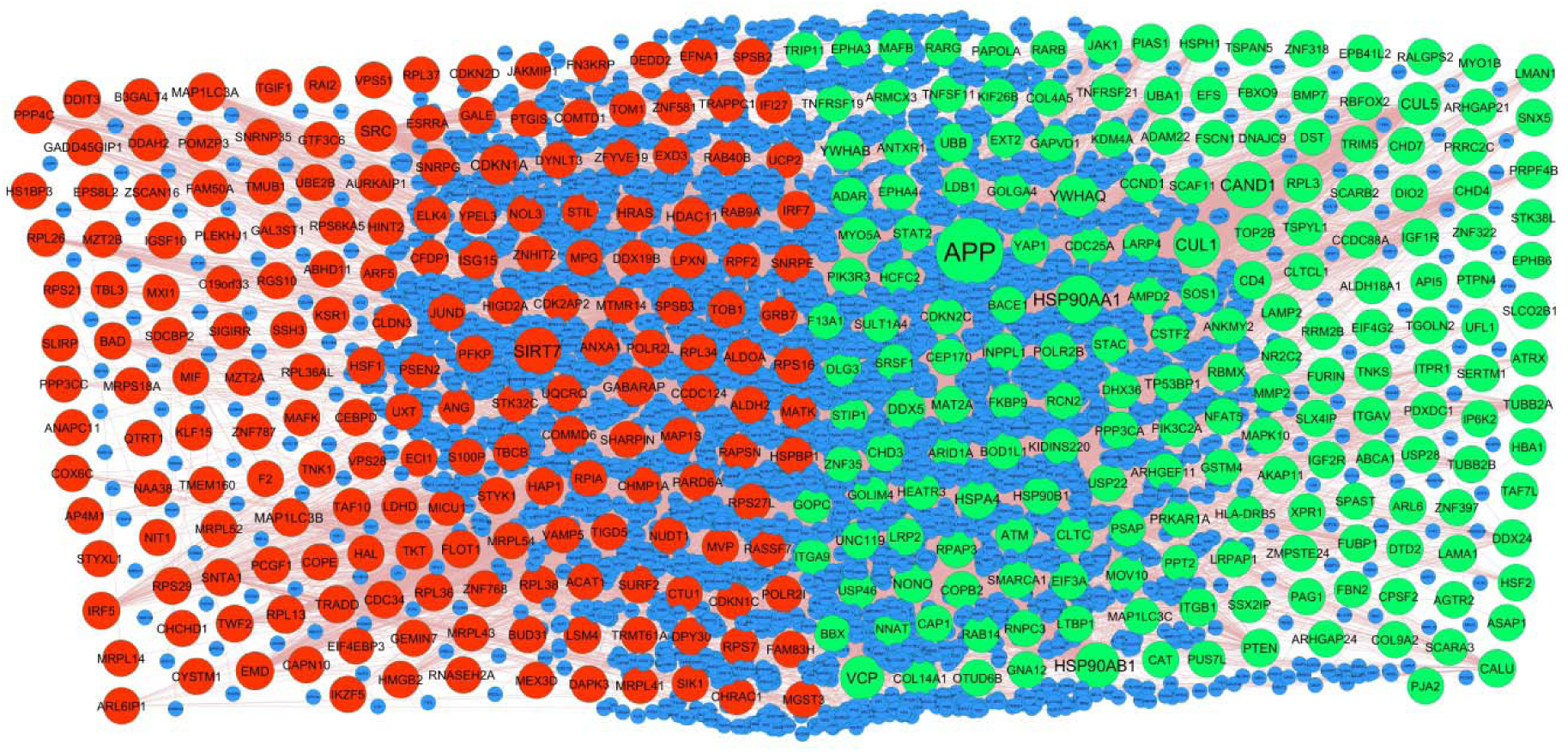
PPI network of DEGs. Up regulated genes are marked in green; down regulated genes are marked in red.

**Fig. 4.**
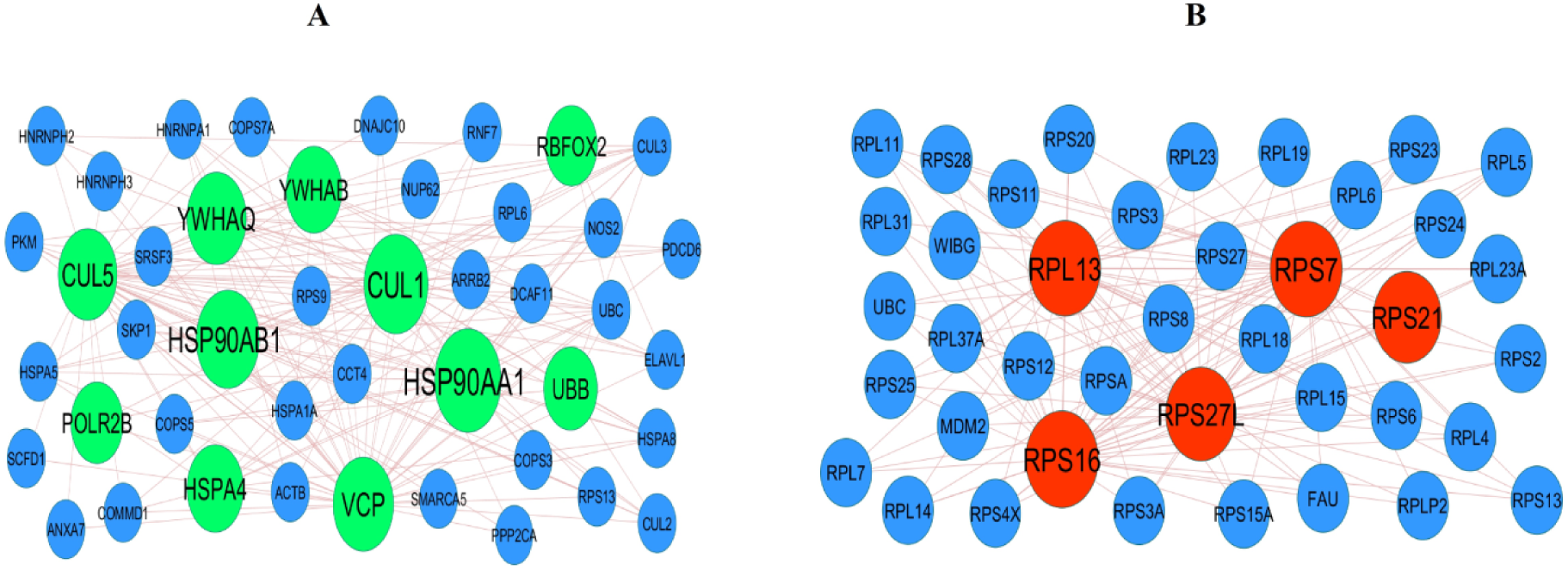
Modules selected from the PPI network. (A) The most significant module was obtained from PPI network with 44 nodes and 177 edges for up regulated genes (B) The most significant module was obtained from PPI network with 40 nodes and 141 edges for down regulated genes. Up regulated genes are marked in parrot green; down regulated genes are marked in red.

**Table 4.**
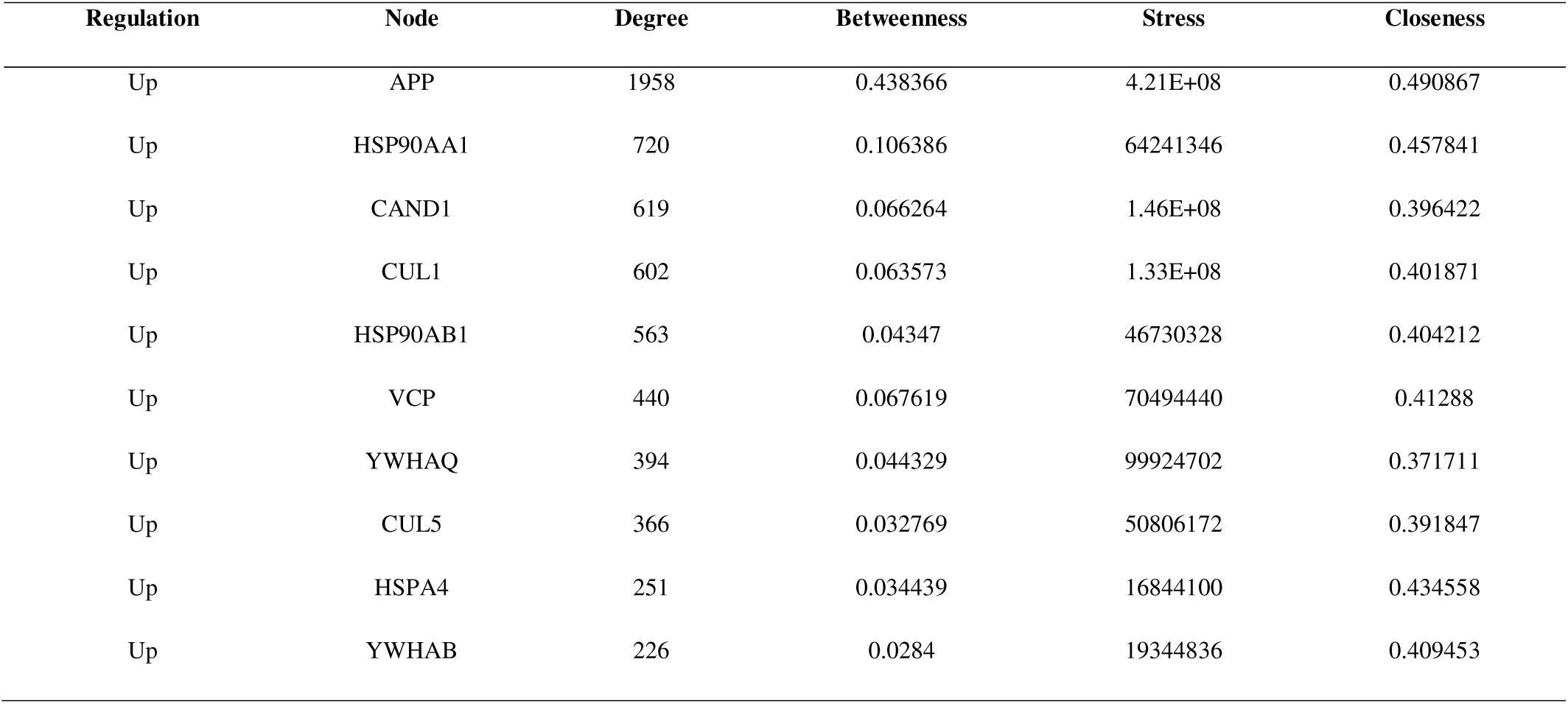
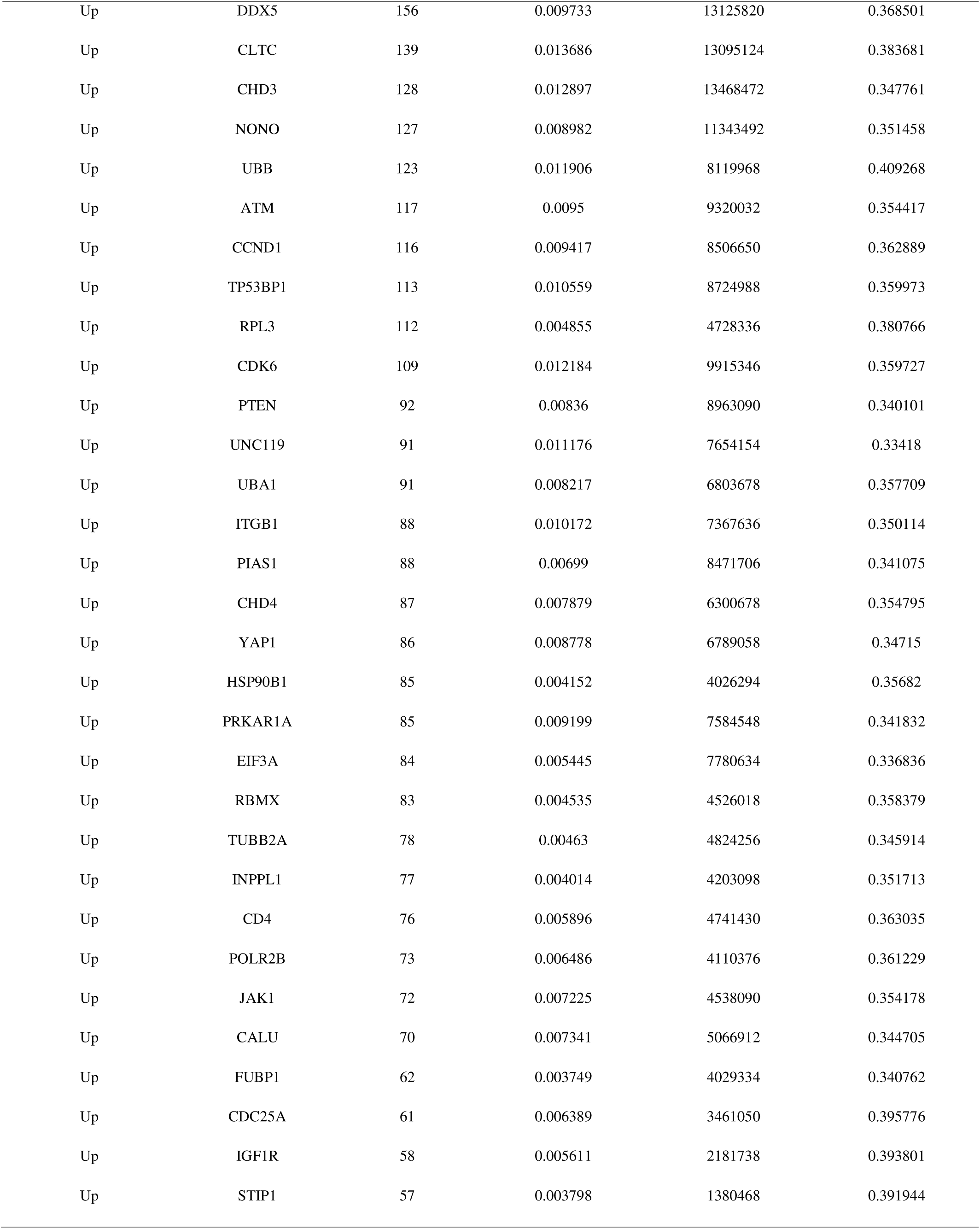
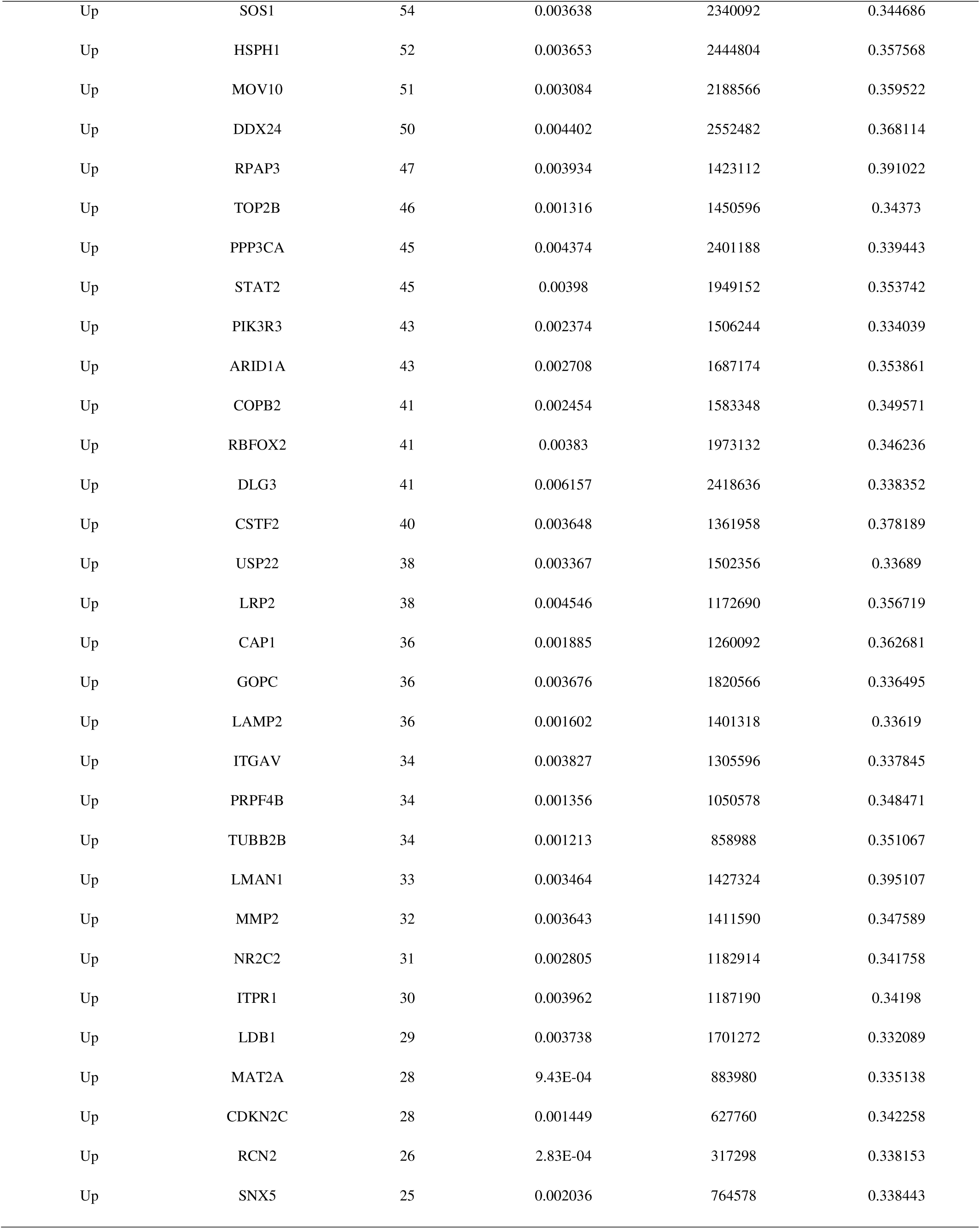
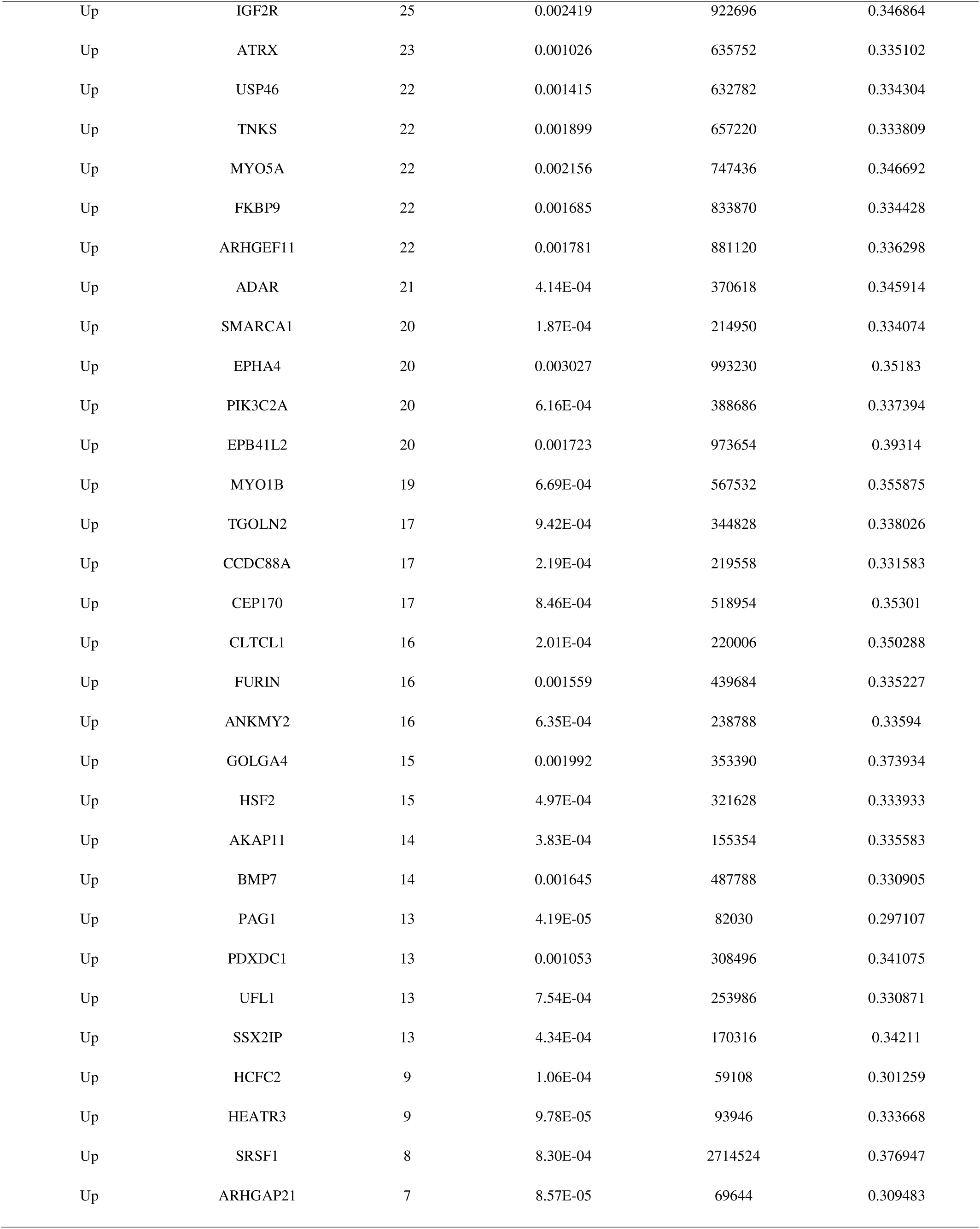
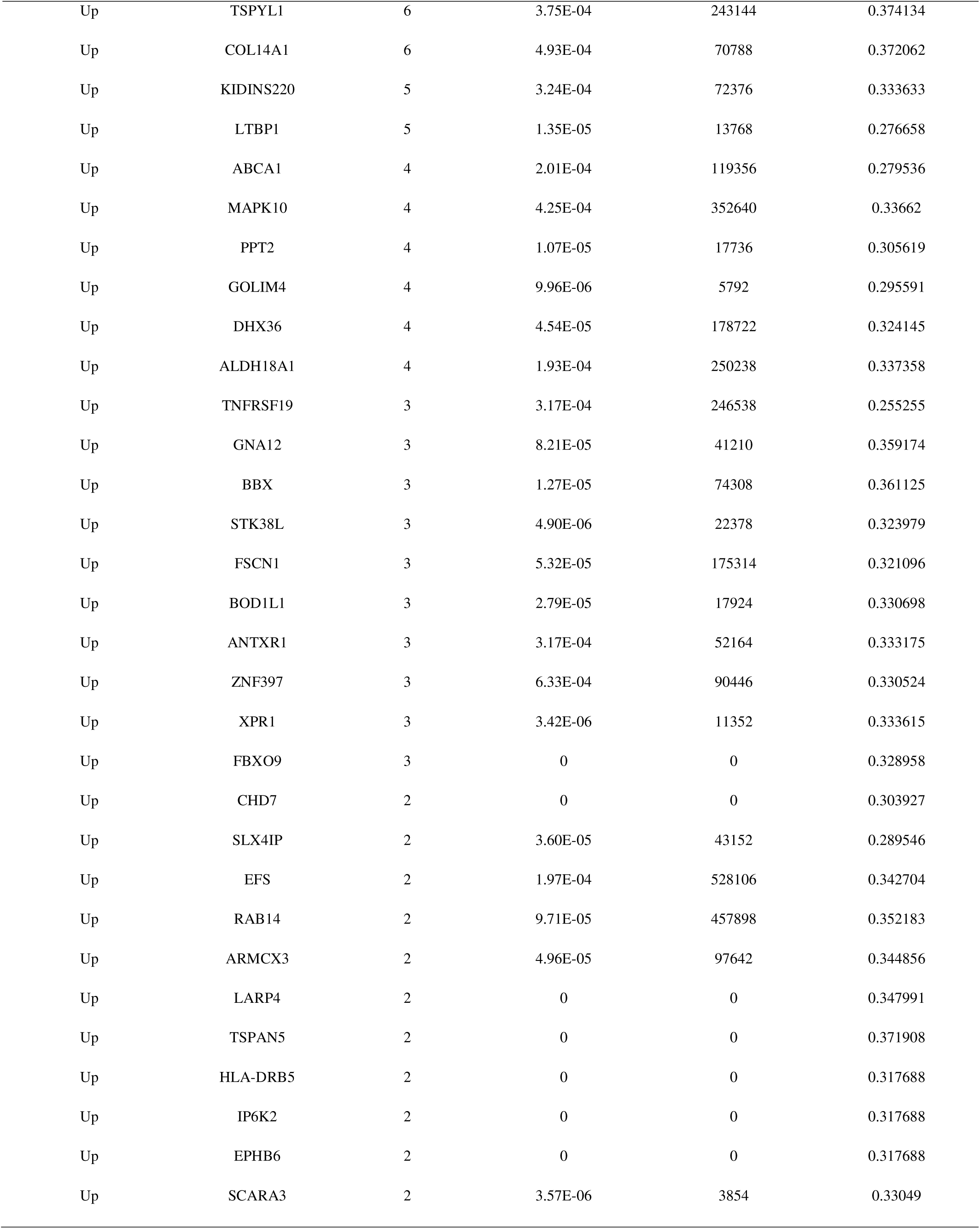
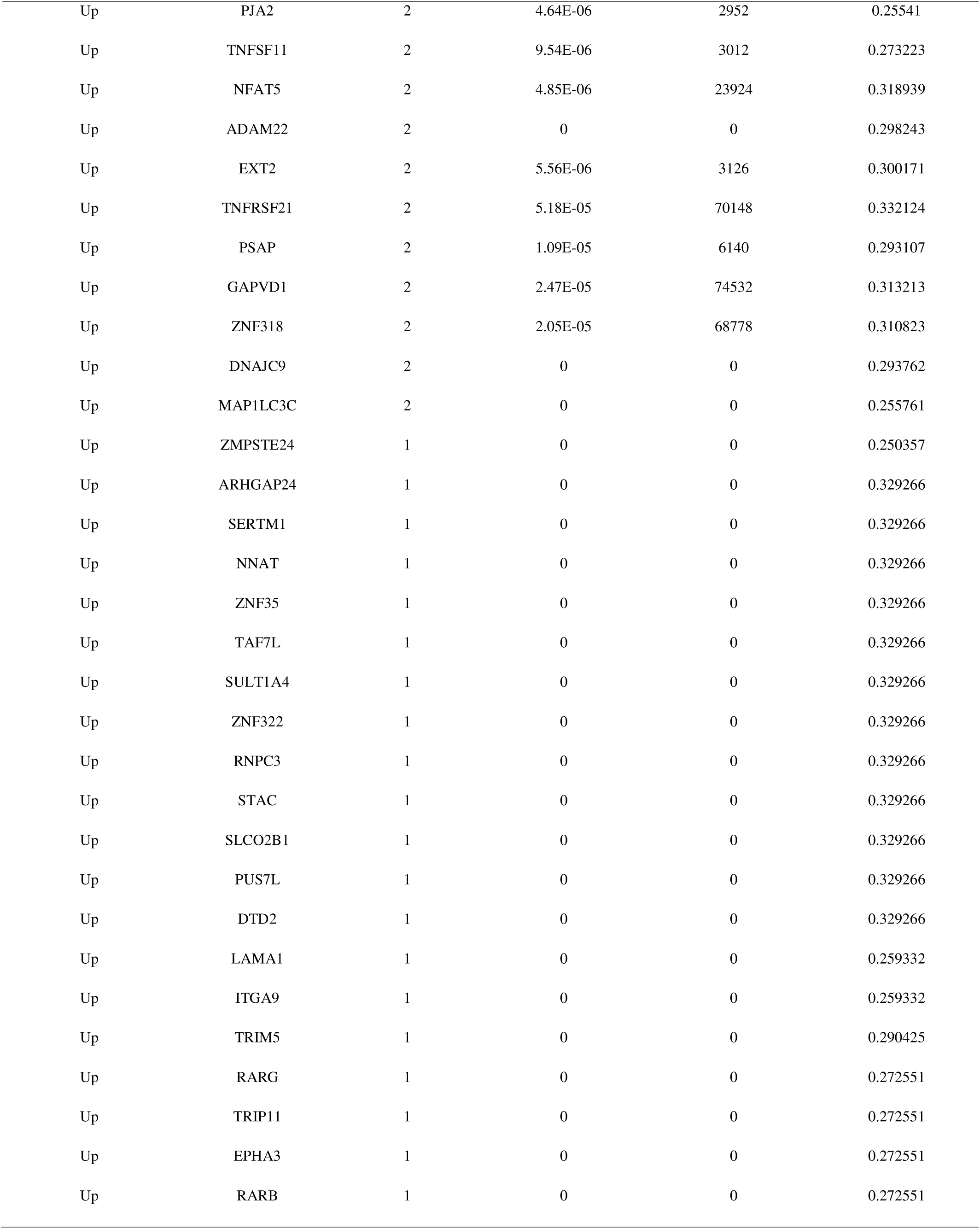
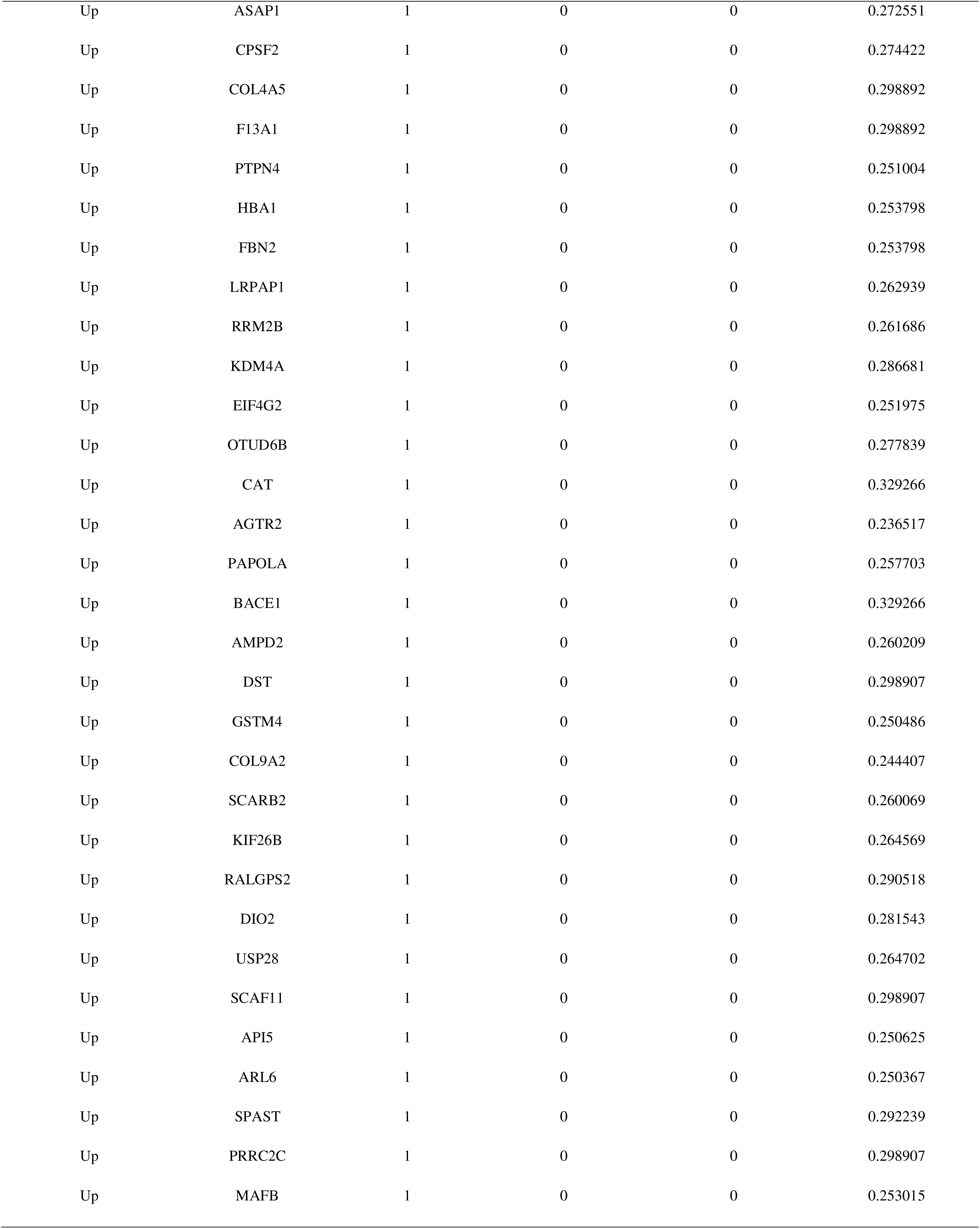
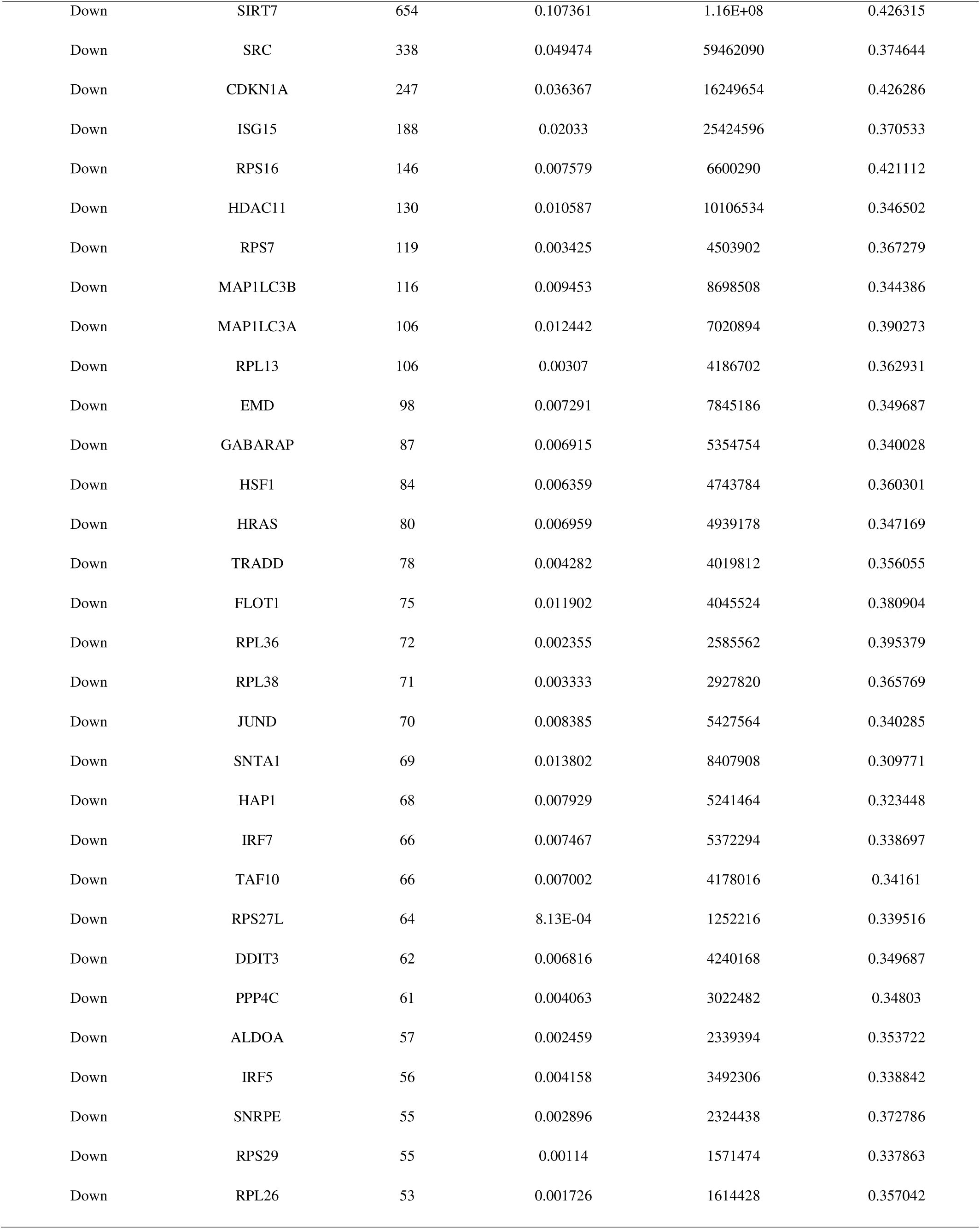
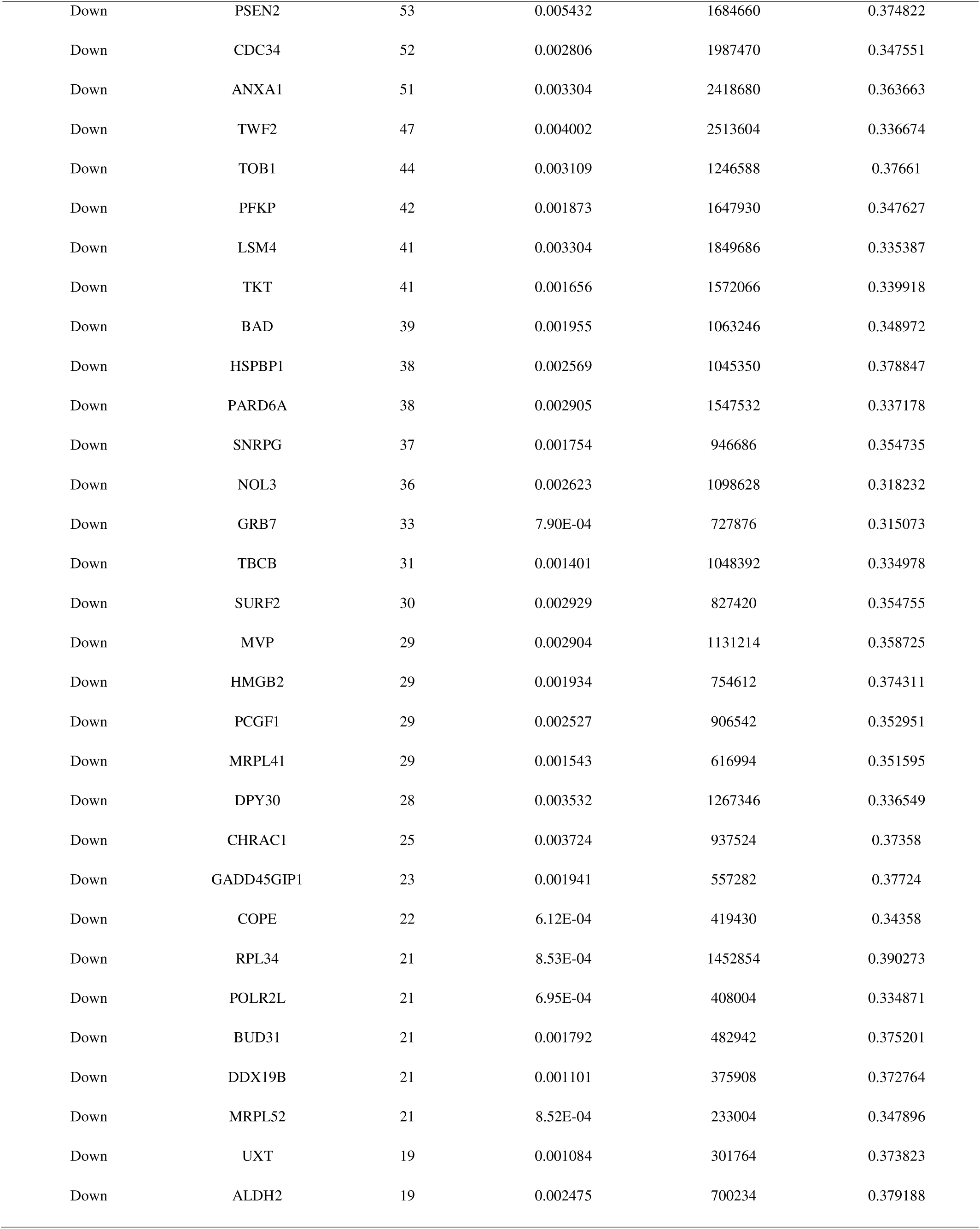
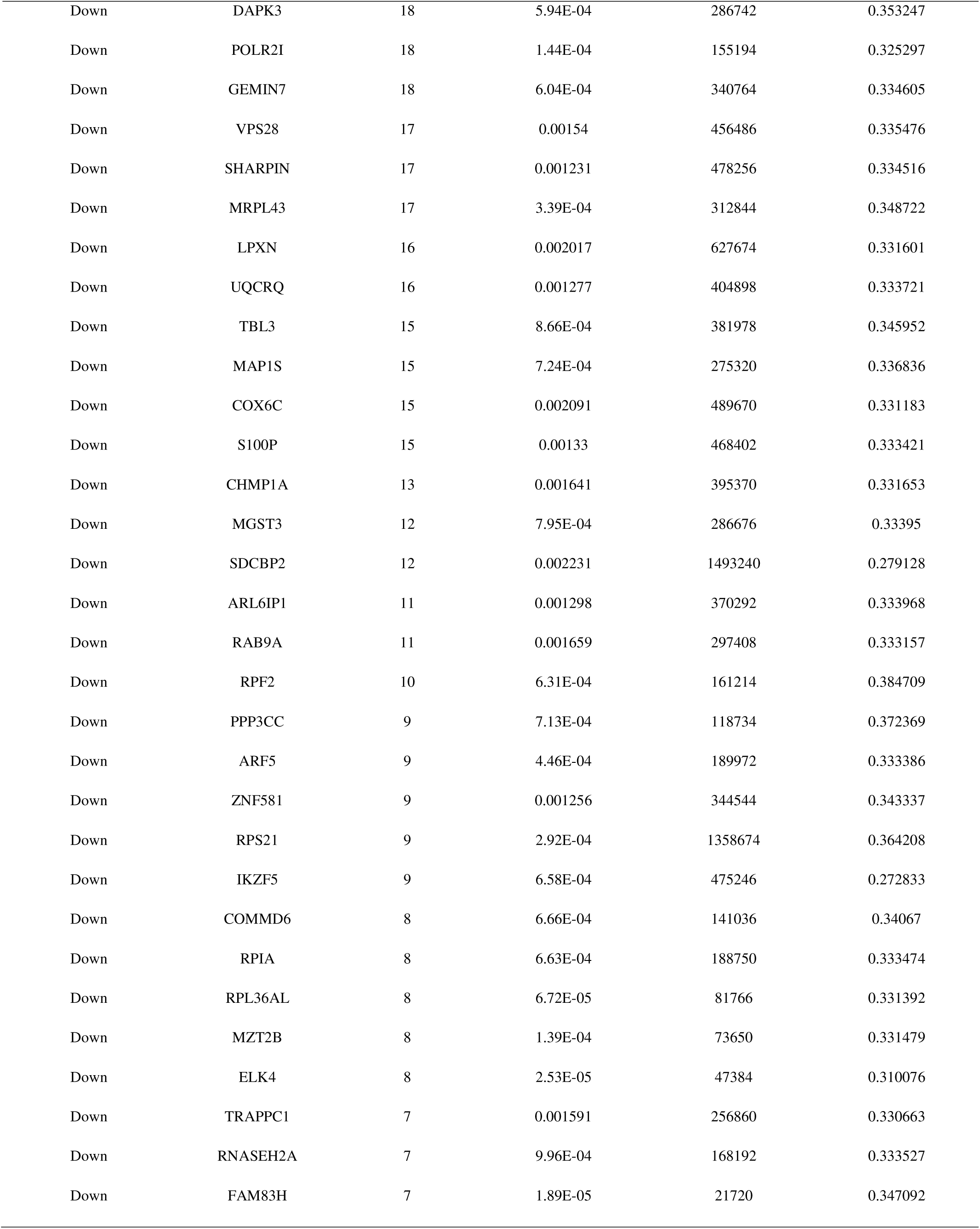
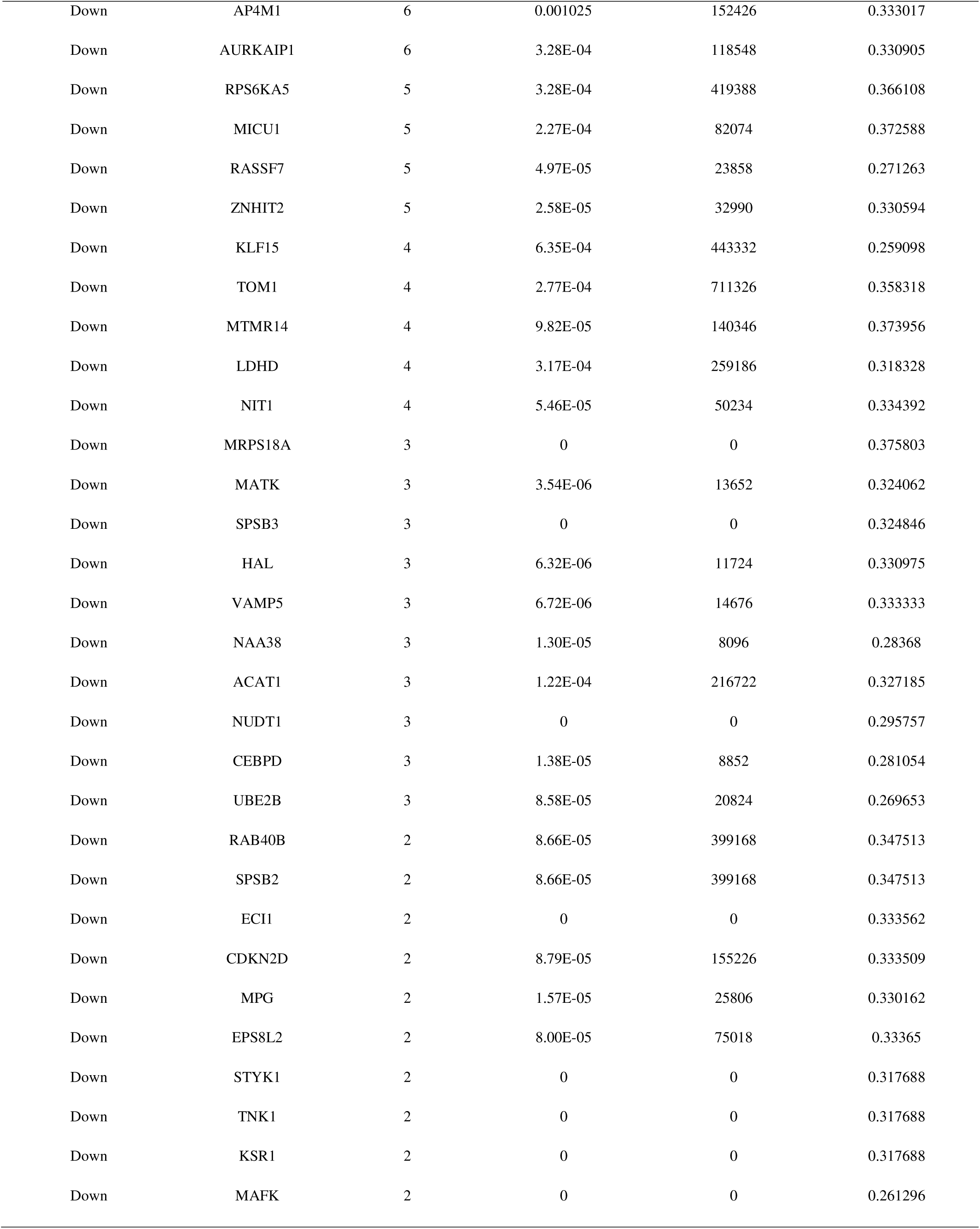
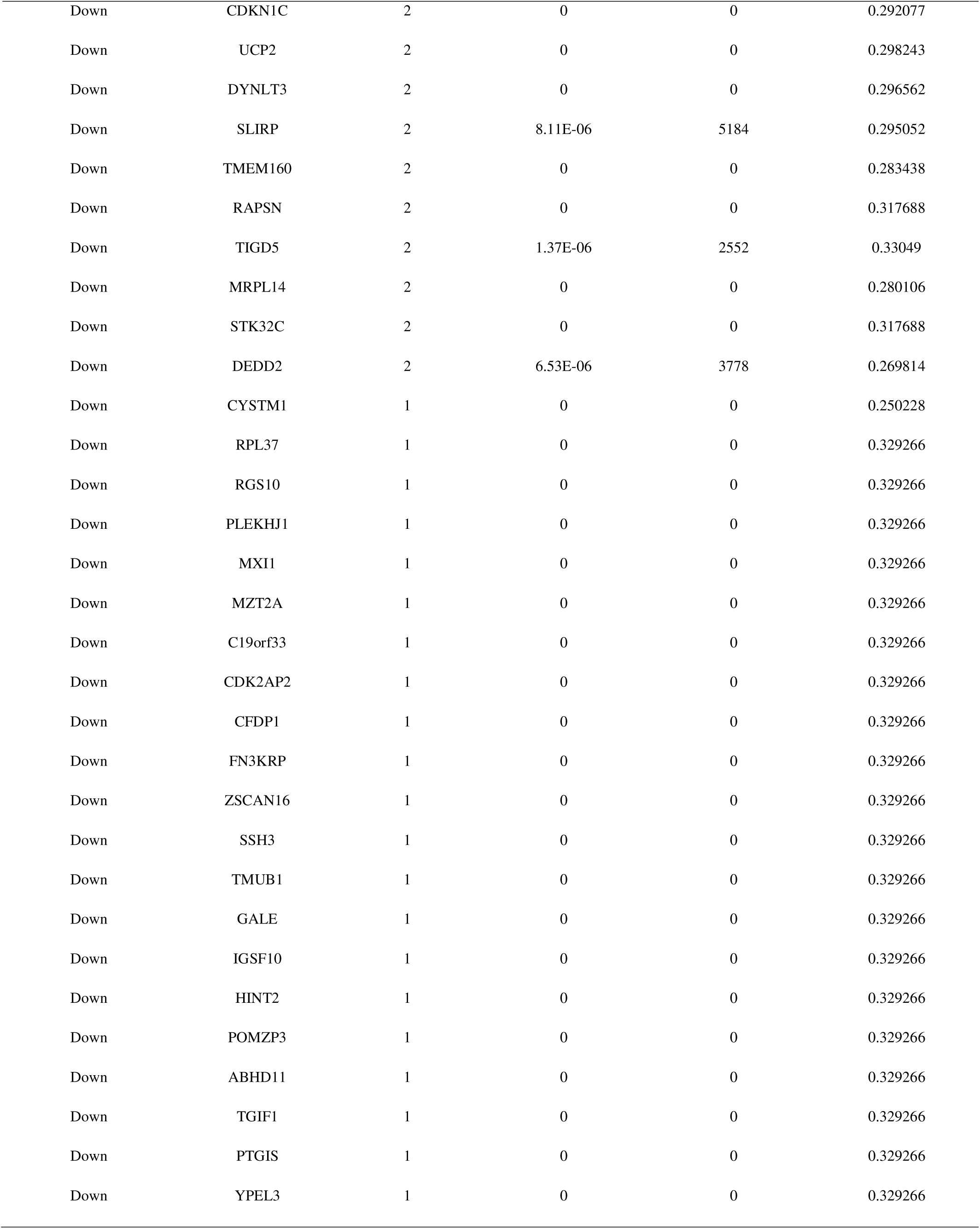
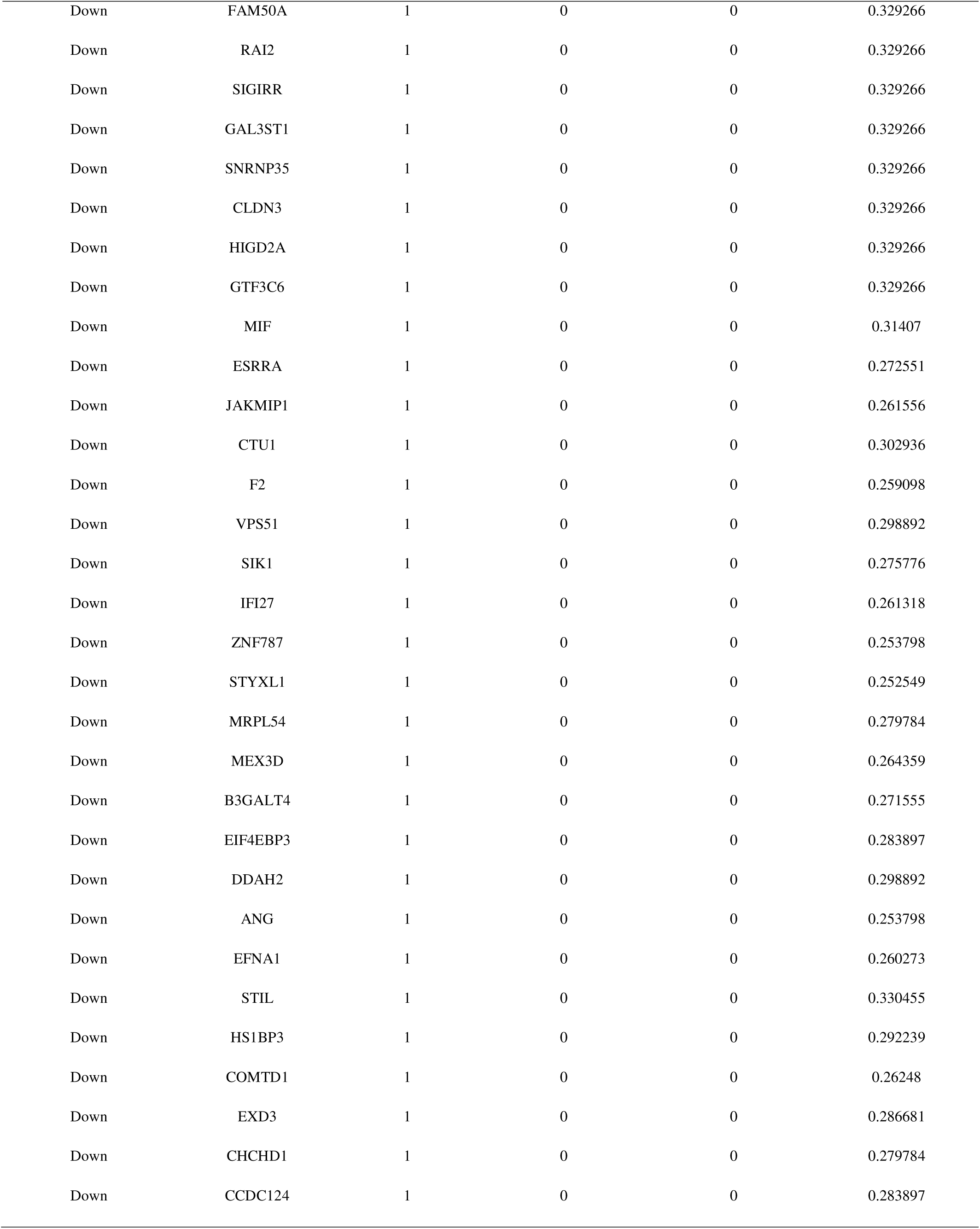
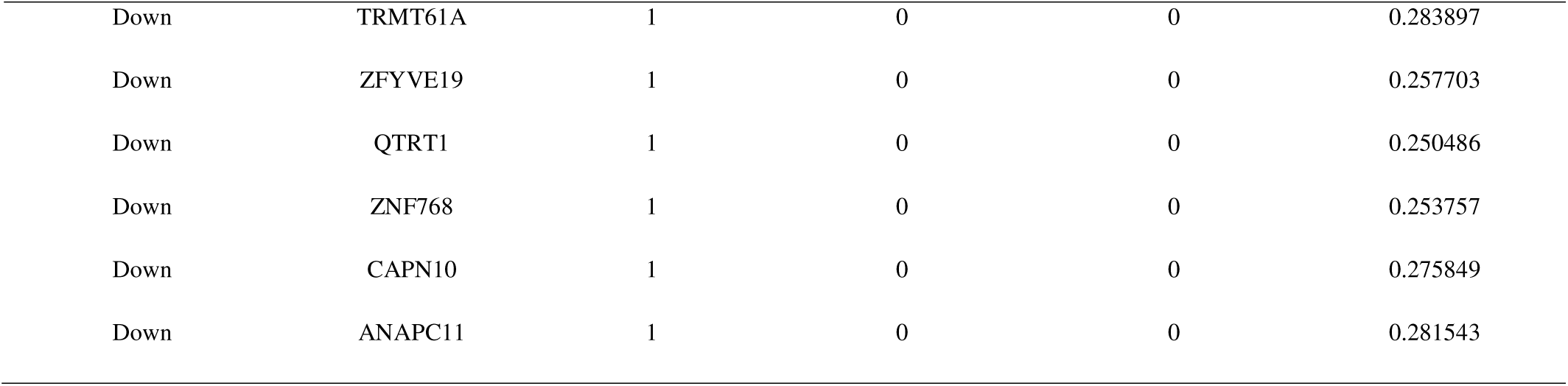
Topology table for up and down regulated genes.

| Regulation | Node | Degree | Betweenness | Stress | Closeness |
| --- | --- | --- | --- | --- | --- |
| Up | APP | 1958 | 0.438366 | 4.21E+08 | 0.490867 |
| Up | HSP90AA1 | 720 | 0.106386 | 64241346 | 0.457841 |
| Up | CAND1 | 619 | 0.066264 | 1.46E+08 | 0.396422 |
| Up | CUL1 | 602 | 0.063573 | 1.33E+08 | 0.401871 |
| Up | HSP90AB1 | 563 | 0.04347 | 46730328 | 0.404212 |
| Up | VCP | 440 | 0.067619 | 70494440 | 0.41288 |
| Up | YWHAQ | 394 | 0.044329 | 99924702 | 0.371711 |
| Up | CUL5 | 366 | 0.032769 | 50806172 | 0.391847 |
| Up | HSPA4 | 251 | 0.034439 | 16844100 | 0.434558 |
| Up | YWHAB | 226 | 0.0284 | 19344836 | 0.409453 |
| Up | DDX5 | 156 | 0.009733 | 13125820 | 0.368501 |
| Up | CLTC | 139 | 0.013686 | 13095124 | 0.383681 |
| Up | CHD3 | 128 | 0.012897 | 13468472 | 0.347761 |
| Up | NONO | 127 | 0.008982 | 11343492 | 0.351458 |
| Up | UBB | 123 | 0.011906 | 8119968 | 0.409268 |
| Up | ATM | 117 | 0.0095 | 9320032 | 0.354417 |
| Up | CCND1 | 116 | 0.009417 | 8506650 | 0.362889 |
| Up | TP53BP1 | 113 | 0.010559 | 8724988 | 0.359973 |
| Up | RPL3 | 112 | 0.004855 | 4728336 | 0.380766 |
| Up | CDK6 | 109 | 0.012184 | 9915346 | 0.359727 |
| Up | PTEN | 92 | 0.00836 | 8963090 | 0.340101 |
| Up | UNC119 | 91 | 0.011176 | 7654154 | 0.33418 |
| Up | UBA1 | 91 | 0.008217 | 6803678 | 0.357709 |
| Up | ITGB1 | 88 | 0.010172 | 7367636 | 0.350114 |
| Up | PIAS1 | 88 | 0.00699 | 8471706 | 0.341075 |
| Up | CHD4 | 87 | 0.007879 | 6300678 | 0.354795 |
| Up | YAP1 | 86 | 0.008778 | 6789058 | 0.34715 |
| Up | HSP90B1 | 85 | 0.004152 | 4026294 | 0.35682 |
| Up | PRKAR1A | 85 | 0.009199 | 7584548 | 0.341832 |
| Up | EIF3A | 84 | 0.005445 | 7780634 | 0.336836 |
| Up | RBMX | 83 | 0.004535 | 4526018 | 0.358379 |
| Up | TUBB2A | 78 | 0.00463 | 4824256 | 0.345914 |
| Up | INPPL1 | 77 | 0.004014 | 4203098 | 0.351713 |
| Up | CD4 | 76 | 0.005896 | 4741430 | 0.363035 |
| Up | POLR2B | 73 | 0.006486 | 4110376 | 0.361229 |
| Up | JAK1 | 72 | 0.007225 | 4538090 | 0.354178 |
| Up | CALU | 70 | 0.007341 | 5066912 | 0.344705 |
| Up | FUBP1 | 62 | 0.003749 | 4029334 | 0.340762 |
| Up | CDC25A | 61 | 0.006389 | 3461050 | 0.395776 |
| Up | IGF1R | 58 | 0.005611 | 2181738 | 0.393801 |
| Up | STIP1 | 57 | 0.003798 | 1380468 | 0.391944 |
| Up | SOS1 | 54 | 0.003638 | 2340092 | 0.344686 |
| Up | HSPH1 | 52 | 0.003653 | 2444804 | 0.357568 |
| Up | MOV10 | 51 | 0.003084 | 2188566 | 0.359522 |
| Up | DDX24 | 50 | 0.004402 | 2552482 | 0.368114 |
| Up | RPAP3 | 47 | 0.003934 | 1423112 | 0.391022 |
| Up | TOP2B | 46 | 0.001316 | 1450596 | 0.34373 |
| Up | PPP3CA | 45 | 0.004374 | 2401188 | 0.339443 |
| Up | STAT2 | 45 | 0.00398 | 1949152 | 0.353742 |
| Up | PIK3R3 | 43 | 0.002374 | 1506244 | 0.334039 |
| Up | ARID1A | 43 | 0.002708 | 1687174 | 0.353861 |
| Up | COPB2 | 41 | 0.002454 | 1583348 | 0.349571 |
| Up | RBFOX2 | 41 | 0.00383 | 1973132 | 0.346236 |
| Up | DLG3 | 41 | 0.006157 | 2418636 | 0.338352 |
| Up | CSTF2 | 40 | 0.003648 | 1361958 | 0.378189 |
| Up | USP22 | 38 | 0.003367 | 1502356 | 0.33689 |
| Up | LRP2 | 38 | 0.004546 | 1172690 | 0.356719 |
| Up | CAP1 | 36 | 0.001885 | 1260092 | 0.362681 |
| Up | GOPC | 36 | 0.003676 | 1820566 | 0.336495 |
| Up | LAMP2 | 36 | 0.001602 | 1401318 | 0.33619 |
| Up | ITGAV | 34 | 0.003827 | 1305596 | 0.337845 |
| Up | PRPF4B | 34 | 0.001356 | 1050578 | 0.348471 |
| Up | TUBB2B | 34 | 0.001213 | 858988 | 0.351067 |
| Up | LMAN1 | 33 | 0.003464 | 1427324 | 0.395107 |
| Up | MMP2 | 32 | 0.003643 | 1411590 | 0.347589 |
| Up | NR2C2 | 31 | 0.002805 | 1182914 | 0.341758 |
| Up | ITPR1 | 30 | 0.003962 | 1187190 | 0.34198 |
| Up | LDB1 | 29 | 0.003738 | 1701272 | 0.332089 |
| Up | MAT2A | 28 | 9.43E-04 | 883980 | 0.335138 |
| Up | CDKN2C | 28 | 0.001449 | 627760 | 0.342258 |
| Up | RCN2 | 26 | 2.83E-04 | 317298 | 0.338153 |
| Up | SNX5 | 25 | 0.002036 | 764578 | 0.338443 |
| Up | IGF2R | 25 | 0.002419 | 922696 | 0.346864 |
| Up | ATRX | 23 | 0.001026 | 635752 | 0.335102 |
| Up | USP46 | 22 | 0.001415 | 632782 | 0.334304 |
| Up | TNKS | 22 | 0.001899 | 657220 | 0.333809 |
| Up | MYO5A | 22 | 0.002156 | 747436 | 0.346692 |
| Up | FKBP9 | 22 | 0.001685 | 833870 | 0.334428 |
| Up | ARHGEF11 | 22 | 0.001781 | 881120 | 0.336298 |
| Up | ADAR | 21 | 4.14E-04 | 370618 | 0.345914 |
| Up | SMARCA1 | 20 | 1.87E-04 | 214950 | 0.334074 |
| Up | EPHA4 | 20 | 0.003027 | 993230 | 0.35183 |
| Up | PIK3C2A | 20 | 6.16E-04 | 388686 | 0.337394 |
| Up | EPB41L2 | 20 | 0.001723 | 973654 | 0.39314 |
| Up | MYO1B | 19 | 6.69E-04 | 567532 | 0.355875 |
| Up | TGOLN2 | 17 | 9.42E-04 | 344828 | 0.338026 |
| Up | CCDC88A | 17 | 2.19E-04 | 219558 | 0.331583 |
| Up | CEP170 | 17 | 8.46E-04 | 518954 | 0.35301 |
| Up | CLTCL1 | 16 | 2.01E-04 | 220006 | 0.350288 |
| Up | FURIN | 16 | 0.001559 | 439684 | 0.335227 |
| Up | ANKMY2 | 16 | 6.35E-04 | 238788 | 0.33594 |
| Up | GOLGA4 | 15 | 0.001992 | 353390 | 0.373934 |
| Up | HSF2 | 15 | 4.97E-04 | 321628 | 0.333933 |
| Up | AKAP11 | 14 | 3.83E-04 | 155354 | 0.335583 |
| Up | BMP7 | 14 | 0.001645 | 487788 | 0.330905 |
| Up | PAG1 | 13 | 4.19E-05 | 82030 | 0.297107 |
| Up | PDXDC1 | 13 | 0.001053 | 308496 | 0.341075 |
| Up | UFL1 | 13 | 7.54E-04 | 253986 | 0.330871 |
| Up | SSX2IP | 13 | 4.34E-04 | 170316 | 0.34211 |
| Up | HCFC2 | 9 | 1.06E-04 | 59108 | 0.301259 |
| Up | HEATR3 | 9 | 9.78E-05 | 93946 | 0.333668 |
| Up | SRSF1 | 8 | 8.30E-04 | 2714524 | 0.376947 |
| Up | ARHGAP21 | 7 | 8.57E-05 | 69644 | 0.309483 |
| Up | TSPYL1 | 6 | 3.75E-04 | 243144 | 0.374134 |
| Up | COL14A1 | 6 | 4.93E-04 | 70788 | 0.372062 |
| Up | KIDINS220 | 5 | 3.24E-04 | 72376 | 0.333633 |
| Up | LTBP1 | 5 | 1.35E-05 | 13768 | 0.276658 |
| Up | ABCA1 | 4 | 2.01E-04 | 119356 | 0.279536 |
| Up | MAPK10 | 4 | 4.25E-04 | 352640 | 0.33662 |
| Up | PPT2 | 4 | 1.07E-05 | 17736 | 0.305619 |
| Up | GOLIM4 | 4 | 9.96E-06 | 5792 | 0.295591 |
| Up | DHX36 | 4 | 4.54E-05 | 178722 | 0.324145 |
| Up | ALDH18A1 | 4 | 1.93E-04 | 250238 | 0.337358 |
| Up | TNFRSF19 | 3 | 3.17E-04 | 246538 | 0.255255 |
| Up | GNA12 | 3 | 8.21E-05 | 41210 | 0.359174 |
| Up | BBX | 3 | 1.27E-05 | 74308 | 0.361125 |
| Up | STK38L | 3 | 4.90E-06 | 22378 | 0.323979 |
| Up | FSCN1 | 3 | 5.32E-05 | 175314 | 0.321096 |
| Up | BOD1L1 | 3 | 2.79E-05 | 17924 | 0.330698 |
| Up | ANTXR1 | 3 | 3.17E-04 | 52164 | 0.333175 |
| Up | ZNF397 | 3 | 6.33E-04 | 90446 | 0.330524 |
| Up | XPR1 | 3 | 3.42E-06 | 11352 | 0.333615 |
| Up | FBXO9 | 3 | 0 | 0 | 0.328958 |
| Up | CHD7 | 2 | 0 | 0 | 0.303927 |
| Up | SLX4IP | 2 | 3.60E-05 | 43152 | 0.289546 |
| Up | EFS | 2 | 1.97E-04 | 528106 | 0.342704 |
| Up | RAB14 | 2 | 9.71E-05 | 457898 | 0.352183 |
| Up | ARMCX3 | 2 | 4.96E-05 | 97642 | 0.344856 |
| Up | LARP4 | 2 | 0 | 0 | 0.347991 |
| Up | TSPAN5 | 2 | 0 | 0 | 0.371908 |
| Up | HLA-DRB5 | 2 | 0 | 0 | 0.317688 |
| Up | IP6K2 | 2 | 0 | 0 | 0.317688 |
| Up | EPHB6 | 2 | 0 | 0 | 0.317688 |
| Up | SCARA3 | 2 | 3.57E-06 | 3854 | 0.33049 |
| Up | PJA2 | 2 | 4.64E-06 | 2952 | 0.25541 |
| Up | TNFSF11 | 2 | 9.54E-06 | 3012 | 0.273223 |
| Up | NFAT5 | 2 | 4.85E-06 | 23924 | 0.318939 |
| Up | ADAM22 | 2 | 0 | 0 | 0.298243 |
| Up | EXT2 | 2 | 5.56E-06 | 3126 | 0.300171 |
| Up | TNFRSF21 | 2 | 5.18E-05 | 70148 | 0.332124 |
| Up | PSAP | 2 | 1.09E-05 | 6140 | 0.293107 |
| Up | GAPVD1 | 2 | 2.47E-05 | 74532 | 0.313213 |
| Up | ZNF318 | 2 | 2.05E-05 | 68778 | 0.310823 |
| Up | DNAJC9 | 2 | 0 | 0 | 0.293762 |
| Up | MAP1LC3C | 2 | 0 | 0 | 0.255761 |
| Up | ZMPSTE24 | 1 | 0 | 0 | 0.250357 |
| Up | ARHGAP24 | 1 | 0 | 0 | 0.329266 |
| Up | SERTM1 | 1 | 0 | 0 | 0.329266 |
| Up | NNAT | 1 | 0 | 0 | 0.329266 |
| Up | ZNF35 | 1 | 0 | 0 | 0.329266 |
| Up | TAF7L | 1 | 0 | 0 | 0.329266 |
| Up | SULT1A4 | 1 | 0 | 0 | 0.329266 |
| Up | ZNF322 | 1 | 0 | 0 | 0.329266 |
| Up | RNPC3 | 1 | 0 | 0 | 0.329266 |
| Up | STAC | 1 | 0 | 0 | 0.329266 |
| Up | SLCO2B1 | 1 | 0 | 0 | 0.329266 |
| Up | PUS7L | 1 | 0 | 0 | 0.329266 |
| Up | DTD2 | 1 | 0 | 0 | 0.329266 |
| Up | LAMA1 | 1 | 0 | 0 | 0.259332 |
| Up | ITGA9 | 1 | 0 | 0 | 0.259332 |
| Up | TRIM5 | 1 | 0 | 0 | 0.290425 |
| Up | RARG | 1 | 0 | 0 | 0.272551 |
| Up | TRIP11 | 1 | 0 | 0 | 0.272551 |
| Up | EPHA3 | 1 | 0 | 0 | 0.272551 |
| Up | RARB | 1 | 0 | 0 | 0.272551 |
| Up | ASAP1 | 1 | 0 | 0 | 0.272551 |
| Up | CPSF2 | 1 | 0 | 0 | 0.274422 |
| Up | COL4A5 | 1 | 0 | 0 | 0.298892 |
| Up | F13A1 | 1 | 0 | 0 | 0.298892 |
| Up | PTPN4 | 1 | 0 | 0 | 0.251004 |
| Up | HBA1 | 1 | 0 | 0 | 0.253798 |
| Up | FBN2 | 1 | 0 | 0 | 0.253798 |
| Up | LRPAP1 | 1 | 0 | 0 | 0.262939 |
| Up | RRM2B | 1 | 0 | 0 | 0.261686 |
| Up | KDM4A | 1 | 0 | 0 | 0.286681 |
| Up | EIF4G2 | 1 | 0 | 0 | 0.251975 |
| Up | OTUD6B | 1 | 0 | 0 | 0.277839 |
| Up | CAT | 1 | 0 | 0 | 0.329266 |
| Up | AGTR2 | 1 | 0 | 0 | 0.236517 |
| Up | PAPOLA | 1 | 0 | 0 | 0.257703 |
| Up | BACE1 | 1 | 0 | 0 | 0.329266 |
| Up | AMPD2 | 1 | 0 | 0 | 0.260209 |
| Up | DST | 1 | 0 | 0 | 0.298907 |
| Up | GSTM4 | 1 | 0 | 0 | 0.250486 |
| Up | COL9A2 | 1 | 0 | 0 | 0.244407 |
| Up | SCARB2 | 1 | 0 | 0 | 0.260069 |
| Up | KIF26B | 1 | 0 | 0 | 0.264569 |
| Up | RALGPS2 | 1 | 0 | 0 | 0.290518 |
| Up | DIO2 | 1 | 0 | 0 | 0.281543 |
| Up | USP28 | 1 | 0 | 0 | 0.264702 |
| Up | SCAF11 | 1 | 0 | 0 | 0.298907 |
| Up | API5 | 1 | 0 | 0 | 0.250625 |
| Up | ARL6 | 1 | 0 | 0 | 0.250367 |
| Up | SPAST | 1 | 0 | 0 | 0.292239 |
| Up | PRRC2C | 1 | 0 | 0 | 0.298907 |
| Up | MAFB | 1 | 0 | 0 | 0.253015 |
| Down | SIRT7 | 654 | 0.107361 | 1.16E+08 | 0.426315 |
| Down | SRC | 338 | 0.049474 | 59462090 | 0.374644 |
| Down | CDKN1A | 247 | 0.036367 | 16249654 | 0.426286 |
| Down | ISG15 | 188 | 0.02033 | 25424596 | 0.370533 |
| Down | RPS16 | 146 | 0.007579 | 6600290 | 0.421112 |
| Down | HDAC11 | 130 | 0.010587 | 10106534 | 0.346502 |
| Down | RPS7 | 119 | 0.003425 | 4503902 | 0.367279 |
| Down | MAP1LC3B | 116 | 0.009453 | 8698508 | 0.344386 |
| Down | MAP1LC3A | 106 | 0.012442 | 7020894 | 0.390273 |
| Down | RPL13 | 106 | 0.00307 | 4186702 | 0.362931 |
| Down | EMD | 98 | 0.007291 | 7845186 | 0.349687 |
| Down | GABARAP | 87 | 0.006915 | 5354754 | 0.340028 |
| Down | HSF1 | 84 | 0.006359 | 4743784 | 0.360301 |
| Down | HRAS | 80 | 0.006959 | 4939178 | 0.347169 |
| Down | TRADD | 78 | 0.004282 | 4019812 | 0.356055 |
| Down | FLOT1 | 75 | 0.011902 | 4045524 | 0.380904 |
| Down | RPL36 | 72 | 0.002355 | 2585562 | 0.395379 |
| Down | RPL38 | 71 | 0.003333 | 2927820 | 0.365769 |
| Down | JUND | 70 | 0.008385 | 5427564 | 0.340285 |
| Down | SNTA1 | 69 | 0.013802 | 8407908 | 0.309771 |
| Down | HAP1 | 68 | 0.007929 | 5241464 | 0.323448 |
| Down | IRF7 | 66 | 0.007467 | 5372294 | 0.338697 |
| Down | TAF10 | 66 | 0.007002 | 4178016 | 0.34161 |
| Down | RPS27L | 64 | 8.13E-04 | 1252216 | 0.339516 |
| Down | DDIT3 | 62 | 0.006816 | 4240168 | 0.349687 |
| Down | PPP4C | 61 | 0.004063 | 3022482 | 0.34803 |
| Down | ALDOA | 57 | 0.002459 | 2339394 | 0.353722 |
| Down | IRF5 | 56 | 0.004158 | 3492306 | 0.338842 |
| Down | SNRPE | 55 | 0.002896 | 2324438 | 0.372786 |
| Down | RPS29 | 55 | 0.00114 | 1571474 | 0.337863 |
| Down | RPL26 | 53 | 0.001726 | 1614428 | 0.357042 |
| Down | PSEN2 | 53 | 0.005432 | 1684660 | 0.374822 |
| Down | CDC34 | 52 | 0.002806 | 1987470 | 0.347551 |
| Down | ANXA1 | 51 | 0.003304 | 2418680 | 0.363663 |
| Down | TWF2 | 47 | 0.004002 | 2513604 | 0.336674 |
| Down | TOB1 | 44 | 0.003109 | 1246588 | 0.37661 |
| Down | PFKP | 42 | 0.001873 | 1647930 | 0.347627 |
| Down | LSM4 | 41 | 0.003304 | 1849686 | 0.335387 |
| Down | TKT | 41 | 0.001656 | 1572066 | 0.339918 |
| Down | BAD | 39 | 0.001955 | 1063246 | 0.348972 |
| Down | HSPBP1 | 38 | 0.002569 | 1045350 | 0.378847 |
| Down | PARD6A | 38 | 0.002905 | 1547532 | 0.337178 |
| Down | SNRPG | 37 | 0.001754 | 946686 | 0.354735 |
| Down | NOL3 | 36 | 0.002623 | 1098628 | 0.318232 |
| Down | GRB7 | 33 | 7.90E-04 | 727876 | 0.315073 |
| Down | TBCB | 31 | 0.001401 | 1048392 | 0.334978 |
| Down | SURF2 | 30 | 0.002929 | 827420 | 0.354755 |
| Down | MVP | 29 | 0.002904 | 1131214 | 0.358725 |
| Down | HMGB2 | 29 | 0.001934 | 754612 | 0.374311 |
| Down | PCGF1 | 29 | 0.002527 | 906542 | 0.352951 |
| Down | MRPL41 | 29 | 0.001543 | 616994 | 0.351595 |
| Down | DPY30 | 28 | 0.003532 | 1267346 | 0.336549 |
| Down | CHRA1 | 25 | 0.003724 | 937524 | 0.37358 |
| Down | GADD45GIP1 | 23 | 0.001941 | 557282 | 0.37724 |
| Down | COPE | 22 | 6.12E-04 | 419430 | 0.34358 |
| Down | RPL34 | 21 | 8.53E-04 | 1452854 | 0.390273 |
| Down | POLR2L | 21 | 6.95E-04 | 408004 | 0.334871 |
| Down | BUD31 | 21 | 0.001792 | 482942 | 0.375201 |
| Down | DDX19B | 21 | 0.001101 | 375908 | 0.372764 |
| Down | MRPL52 | 21 | 8.52E-04 | 233004 | 0.347896 |
| Down | UXT | 19 | 0.001084 | 301764 | 0.373823 |
| Down | ALDH2 | 19 | 0.002475 | 700234 | 0.379188 |
| Down | DAPK3 | 18 | 5.94E-04 | 286742 | 0.353247 |
| Down | POLR2I | 18 | 1.44E-04 | 155194 | 0.325297 |
| Down | GEMIN7 | 18 | 6.04E-04 | 340764 | 0.334605 |
| Down | VPS28 | 17 | 0.00154 | 456486 | 0.335476 |
| Down | SHARPIN | 17 | 0.001231 | 478256 | 0.334516 |
| Down | MRPL43 | 17 | 3.39E-04 | 312844 | 0.348722 |
| Down | LPXN | 16 | 0.002017 | 627674 | 0.331601 |
| Down | UQCRQ | 16 | 0.001277 | 404898 | 0.333721 |
| Down | TBL3 | 15 | 8.66E-04 | 381978 | 0.345952 |
| Down | MAP1S | 15 | 7.24E-04 | 275320 | 0.336836 |
| Down | COX6C | 15 | 0.002091 | 489670 | 0.331183 |
| Down | S100P | 15 | 0.00133 | 468402 | 0.333421 |
| Down | CHMP1A | 13 | 0.001641 | 395370 | 0.331653 |
| Down | MGST3 | 12 | 7.95E-04 | 286676 | 0.33395 |
| Down | SDCBP2 | 12 | 0.002231 | 1493240 | 0.279128 |
| Down | ARL6IP1 | 11 | 0.001298 | 370292 | 0.333968 |
| Down | RAB9A | 11 | 0.001659 | 297408 | 0.333157 |
| Down | RPF2 | 10 | 6.31E-04 | 161214 | 0.384709 |
| Down | PPP3CC | 9 | 7.13E-04 | 118734 | 0.372369 |
| Down | ARF5 | 9 | 4.46E-04 | 189972 | 0.333386 |
| Down | ZNF581 | 9 | 0.001256 | 344544 | 0.343337 |
| Down | RPS21 | 9 | 2.92E-04 | 1358674 | 0.364208 |
| Down | IKZF5 | 9 | 6.58E-04 | 475246 | 0.272833 |
| Down | COMMD6 | 8 | 6.66E-04 | 141036 | 0.34067 |
| Down | RPIA | 8 | 6.63E-04 | 188750 | 0.333474 |
| Down | RPL36AL | 8 | 6.72E-05 | 81766 | 0.331392 |
| Down | MZT2B | 8 | 1.39E-04 | 73650 | 0.331479 |
| Down | ELK4 | 8 | 2.53E-05 | 47384 | 0.310076 |
| Down | TRAPPC1 | 7 | 0.001591 | 256860 | 0.330663 |
| Down | RNASEH2A | 7 | 9.96E-04 | 168192 | 0.333527 |
| Down | FAM83H | 7 | 1.89E-05 | 21720 | 0.347092 |
| Down | AP4M1 | 6 | 0.001025 | 152426 | 0.333017 |
| Down | AURKAIP1 | 6 | 3.28E-04 | 118548 | 0.330905 |
| Down | RPS6KA5 | 5 | 3.28E-04 | 419388 | 0.366108 |
| Down | MICU1 | 5 | 2.27E-04 | 82074 | 0.372588 |
| Down | RASSF7 | 5 | 4.97E-05 | 23858 | 0.271263 |
| Down | ZNHIT2 | 5 | 2.58E-05 | 32990 | 0.330594 |
| Down | KLF15 | 4 | 6.35E-04 | 443332 | 0.259098 |
| Down | TOM1 | 4 | 2.77E-04 | 711326 | 0.358318 |
| Down | MTMR14 | 4 | 9.82E-05 | 140346 | 0.373956 |
| Down | LDHD | 4 | 3.17E-04 | 259186 | 0.318328 |
| Down | NIT1 | 4 | 5.46E-05 | 50234 | 0.334392 |
| Down | MRPS18A | 3 | 0 | 0 | 0.375803 |
| Down | MATK | 3 | 3.54E-06 | 13652 | 0.324062 |
| Down | SPSB3 | 3 | 0 | 0 | 0.324846 |
| Down | HAL | 3 | 6.32E-06 | 11724 | 0.330975 |
| Down | VAMP5 | 3 | 6.72E-06 | 14676 | 0.333333 |
| Down | NAA38 | 3 | 1.30E-05 | 8096 | 0.28368 |
| Down | ACAT1 | 3 | 1.22E-04 | 216722 | 0.327185 |
| Down | NUDT1 | 3 | 0 | 0 | 0.295757 |
| Down | CEBPD | 3 | 1.38E-05 | 8852 | 0.281054 |
| Down | UBE2B | 3 | 8.58E-05 | 20824 | 0.269653 |
| Down | RAB40B | 2 | 8.66E-05 | 399168 | 0.347513 |
| Down | SPSB2 | 2 | 8.66E-05 | 399168 | 0.347513 |
| Down | ECI1 | 2 | 0 | 0 | 0.333562 |
| Down | CDKN2D | 2 | 8.79E-05 | 155226 | 0.333509 |
| Down | MPG | 2 | 1.57E-05 | 25806 | 0.330162 |
| Down | EPS8L2 | 2 | 8.00E-05 | 75018 | 0.33365 |
| Down | STYK1 | 2 | 0 | 0 | 0.317688 |
| Down | TNK1 | 2 | 0 | 0 | 0.317688 |
| Down | KSR1 | 2 | 0 | 0 | 0.317688 |
| Down | MAFK | 2 | 0 | 0 | 0.261296 |
| Down | CDKN1C | 2 | 0 | 0 | 0.292077 |
| Down | UCP2 | 2 | 0 | 0 | 0.298243 |
| Down | DYNLT3 | 2 | 0 | 0 | 0.296562 |
| Down | SLIRP | 2 | 8.11E-06 | 5184 | 0.295052 |
| Down | TMEM160 | 2 | 0 | 0 | 0.283438 |
| Down | RAPSN | 2 | 0 | 0 | 0.317688 |
| Down | TIGD5 | 2 | 1.37E-06 | 2552 | 0.33049 |
| Down | MRPL14 | 2 | 0 | 0 | 0.280106 |
| Down | STK32C | 2 | 0 | 0 | 0.317688 |
| Down | DEDD2 | 2 | 6.53E-06 | 3778 | 0.269814 |
| Down | CYSTM1 | 1 | 0 | 0 | 0.250228 |
| Down | RPL37 | 1 | 0 | 0 | 0.329266 |
| Down | RGS10 | 1 | 0 | 0 | 0.329266 |
| Down | PLEKHJ1 | 1 | 0 | 0 | 0.329266 |
| Down | MXI1 | 1 | 0 | 0 | 0.329266 |
| Down | MZT2A | 1 | 0 | 0 | 0.329266 |
| Down | C19orf33 | 1 | 0 | 0 | 0.329266 |
| Down | CDK2AP2 | 1 | 0 | 0 | 0.329266 |
| Down | CFDP1 | 1 | 0 | 0 | 0.329266 |
| Down | FN3KRP | 1 | 0 | 0 | 0.329266 |
| Down | ZSCAN16 | 1 | 0 | 0 | 0.329266 |
| Down | SSH3 | 1 | 0 | 0 | 0.329266 |
| Down | TMUB1 | 1 | 0 | 0 | 0.329266 |
| Down | GALE | 1 | 0 | 0 | 0.329266 |
| Down | IGSF10 | 1 | 0 | 0 | 0.329266 |
| Down | HINT2 | 1 | 0 | 0 | 0.329266 |
| Down | POMZP3 | 1 | 0 | 0 | 0.329266 |
| Down | ABHD11 | 1 | 0 | 0 | 0.329266 |
| Down | TGIF1 | 1 | 0 | 0 | 0.329266 |
| Down | PTGIS | 1 | 0 | 0 | 0.329266 |
| Down | YPEL3 | 1 | 0 | 0 | 0.329266 |
| Down | FAM50A | 1 | 0 | 0 | 0.329266 |
| Down | RAI2 | 1 | 0 | 0 | 0.329266 |
| Down | SIGIRR | 1 | 0 | 0 | 0.329266 |
| Down | GAL3ST1 | 1 | 0 | 0 | 0.329266 |
| Down | SNRNP35 | 1 | 0 | 0 | 0.329266 |
| Down | CLDN3 | 1 | 0 | 0 | 0.329266 |
| Down | HIGD2A | 1 | 0 | 0 | 0.329266 |
| Down | GTF3C6 | 1 | 0 | 0 | 0.329266 |
| Down | MIF | 1 | 0 | 0 | 0.31407 |
| Down | ESRRA | 1 | 0 | 0 | 0.272551 |
| Down | JAKMIP1 | 1 | 0 | 0 | 0.261556 |
| Down | CTU1 | 1 | 0 | 0 | 0.302936 |
| Down | F2 | 1 | 0 | 0 | 0.259098 |
| Down | VPS51 | 1 | 0 | 0 | 0.298892 |
| Down | SIK1 | 1 | 0 | 0 | 0.275776 |
| Down | IFI27 | 1 | 0 | 0 | 0.261318 |
| Down | ZNF787 | 1 | 0 | 0 | 0.253798 |
| Down | STYXL1 | 1 | 0 | 0 | 0.252549 |
| Down | MRPL54 | 1 | 0 | 0 | 0.279784 |
| Down | MEX3D | 1 | 0 | 0 | 0.264359 |
| Down | B3GALT4 | 1 | 0 | 0 | 0.271555 |
| Down | EIF4EBP3 | 1 | 0 | 0 | 0.283897 |
| Down | DDAH2 | 1 | 0 | 0 | 0.298892 |
| Down | ANG | 1 | 0 | 0 | 0.253798 |
| Down | EFNA1 | 1 | 0 | 0 | 0.260273 |
| Down | STIL | 1 | 0 | 0 | 0.330455 |
| Down | HS1BP3 | 1 | 0 | 0 | 0.292239 |
| Down | COMTD1 | 1 | 0 | 0 | 0.26248 |
| Down | EXD3 | 1 | 0 | 0 | 0.286681 |
| Down | CHCHD1 | 1 | 0 | 0 | 0.279784 |
| Down | CCDC124 | 1 | 0 | 0 | 0.283897 |
| Down | TRMT61A | 1 | 0 | 0 | 0.283897 |
| Down | ZFYVE19 | 1 | 0 | 0 | 0.257703 |
| Down | QTRT1 | 1 | 0 | 0 | 0.250486 |
| Down | ZNF768 | 1 | 0 | 0 | 0.253757 |
| Down | CAPN10 | 1 | 0 | 0 | 0.275849 |
| Down | ANAPC11 | 1 | 0 | 0 | 0.281543 |

### Construction of the miRNA-hub gene regulatory network

The miRNet database was used to anticipate and visualize the miRNA - hub gene regulatory network of hub genes. The miRNA - hub gene regulatory network consisted of 5139 (miRNA: 4751 and hub gene: 388) nodes and 73061 edges (Fig.5). CLTC was found to be regulated by 688 miRNAs (ex: hsa-miR-574-3p), CAND1 was found to be regulated by 604 miRNAs (ex: hsa-mir-382-5p), HSP90AB1 was found to be regulated by 548 miRNAs (ex: hsa-miR-218-1-3p), HSP90AA1 was found to be regulated by 538 miRNAs (ex: hsa-mir-548ad-5p), APP was found to be regulated by 511 miRNAs (ex: hsa-mir-31-5p), CDKN1A was found to be regulated by 919 miRNAs (ex: hsa-mir-208a-3p), RPS16 was found to be regulated by 222 miRNAs (ex: hsa-mir-216b-5p), GABARAP was found to be regulated by 205 miRNAs (ex: hsa-miR-218-5p), MAP1LC3B was found to be regulated by 199 miRNAs (ex: hsa-mir-6132) and SRC was found to be regulated by 155 miRNAs (ex: hsa-mir-205-5p) (Table 5).

**Fig. 5.**
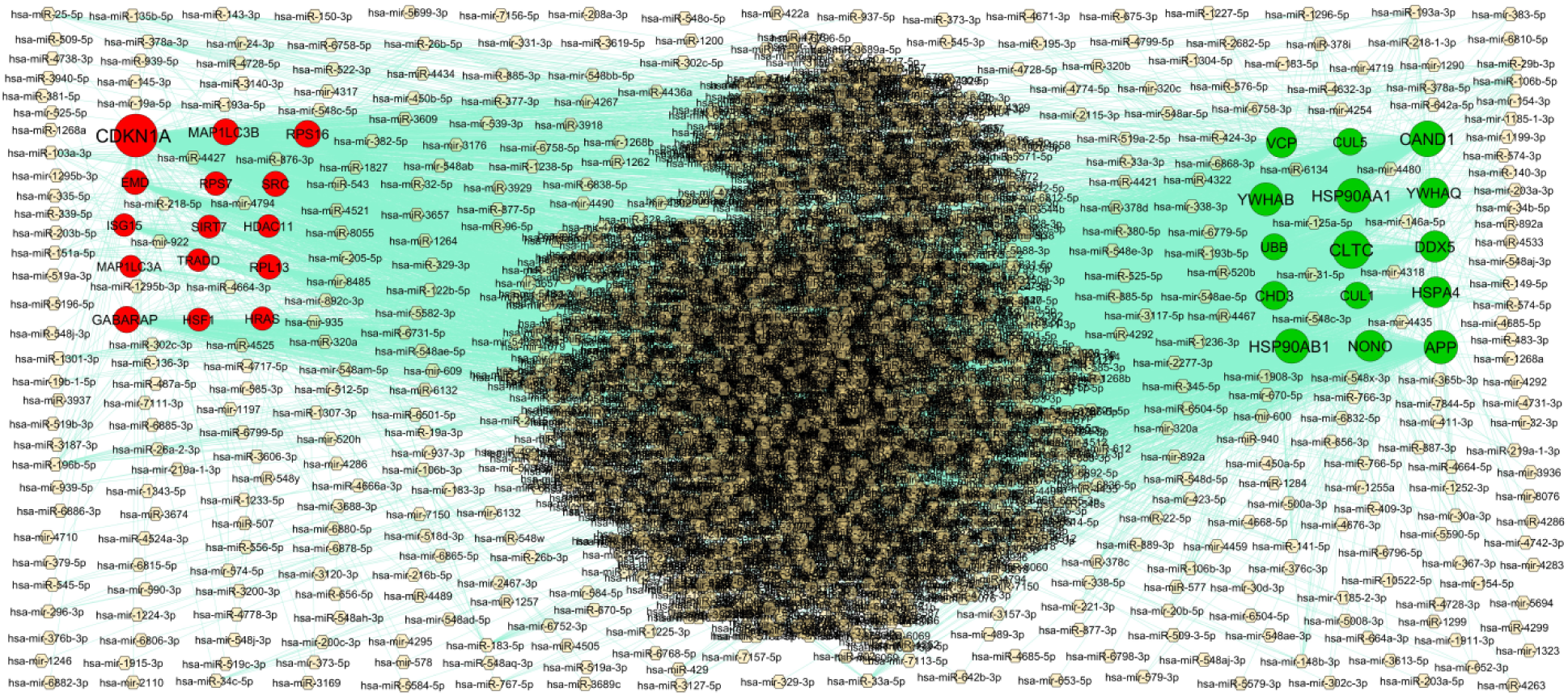
Hub gene - miRNA regulatory network. The light pink color diamond nodes represent the key miRNAs; up regulated genes are marked in green; down regulated genes are marked in red.

**Table 5.** MiRNA - hub gene and TF – hub gene topology table.

| Regulation | Hub Genes | Degree | MicroRNA | Regulation | Hub Genes | Degree | TF |
| --- | --- | --- | --- | --- | --- | --- | --- |
| Up | CLTC | 688 | hsa-miR-574-3p | Up | APP | 16 | SREBF1 |
| Up | CAND1 | 604 | hsa-mir-382-5p | Up | HSP90AA1 | 13 | ARID3A |
| Up | HSP90AB1 | 548 | hsa-miR-218-1-3p | Up | YWHAB | 10 | GATA3 |
| Up | HSP90AA1 | 538 | hsa-mir-548ad-5p | Up | CUL5 | 9 | IRF2 |
| Up | APP | 511 | hsa-mir-31-5p | Up | DDX5 | 8 | BRCA1 |
| Up | YWHAB | 490 | hsa-mir-6758-3p | Up | UBB | 8 | SOX10 |
| Up | DDX5 | 418 | hsa-miR-195-3p | Up | CLTC | 8 | USF2 |
| Up | NONO | 405 | hsa-miR-6504-5p | Up | CHD3 | 7 | STAT1 |
| Up | HSPA4 | 387 | hsa-mir-4286 | Up | HSP90AB1 | 6 | CREB1 |
| Up | VCP | 370 | hsa-miR-193a-3p | Up | VCP | 6 | NFIC |
| Up | YWHAQ | 351 | hsa-miR-940 | Up | HSPA4 | 5 | YY1 |
| Up | CHD3 | 290 | hsa-miR-4664-3p | Up | YWHAQ | 4 | POU2F2 |
| Up | CUL5 | 222 | hsa-miR-548d-5p | Up | CAND1 | 4 | FOXC1 |
| Up | CUL1 | 205 | hsa-miR-302c-3p | Up | CUL1 | 3 | GATA2 |
| Up | UBB | 196 | hsa-mir-1224-3p | Up | NONO | 3 | TP53 |
| Down | CDKN1A | 919 | hsa-mir-208a-3p | Down | CDKN1A | 14 | RELA |
| Down | RPS16 | 222 | hsa-mir-216b-5p | Down | MAP1LC3A | 14 | TFAP2C |
| Down | GABARAP | 205 | hsa-miR-218-5p | Down | HRAS | 14 | MAX |
| Down | MAP1LC3B | 199 | hsa-mir-6132 | Down | EMD | 10 | SPIB |
| Down | SRC | 155 | hsa-mir-205-5p | Down | ISG15 | 10 | NR2E3 |
| Down | EMD | 135 | hsa-miR-320b | Down | HSF1 | 9 | PDX1 |
| Down | RPL13 | 133 | hsa-miR-106b-5p | Down | MAP1LC3B | 8 | ZNF354C |
| Down | RPS7 | 90 | hsa-miR-1264 | Down | HDAC11 | 8 | JUN |
| Down | HRAS | 81 | hsa-miR-183-5p | Down | GABARAP | 8 | HINFP |
| Down | HSF1 | 64 | hsa-miR-96-5p | Down | RPS7 | 8 | PAX2 |
| Down | SIRT7 | 63 | hsa-miR-103a-3p | Down | RPL13 | 8 | ELK4 |
| Down | HDAC11 | 55 | hsa-miR-19a-3p | Down | SRC | 7 | E2F6 |
| Down | ISG15 | 51 | hsa-miR-148b-3p | Down | RPS16 | 7 | STAT3 |
| Down | TRADD | 44 | hsa-miR-22-5p | Down | TRADD | 6 | NFYA |
| Down | MAP1LC3A | 27 | hsa-miR-1284 | Down | SIRT7 | 3 | TFAP2A |

### Construction of the TF-hub gene regulatory network

The NetworkAnalyst database was used to anticipate and visualize the TF - hub gene regulatory network of hub genes. The TF - hub gene regulatory network consisted of 5139 (TF: 97 and hub gene: 386) nodes and 3209 edges (Fig.6). APP was found to be regulated by 16 TFs (ex: SREBF1), HSP90AA1 was found to be regulated by 13 TFs (ex: ARID3A), YWHAB was found to be regulated by 10 TFs (ex: GATA3), CUL5 was found to be regulated by 9 TFs (ex: IRF2), DDX5 was found to be regulated by 8 TFs (ex: BRCA1), CDKN1A was found to be regulated by 14 TFs (ex: RELA), MAP1LC3A was found to be regulated by 14 TFs (ex: TFAP2C), HRAS was found to be regulated by 14 TFs (ex: MAX), EMD was found to be regulated by 10 TFs (ex: SPIB) and ISG15 was found to be regulated by 10 TFs (ex: NR2E3) (Table 5).

**Fig. 6.**
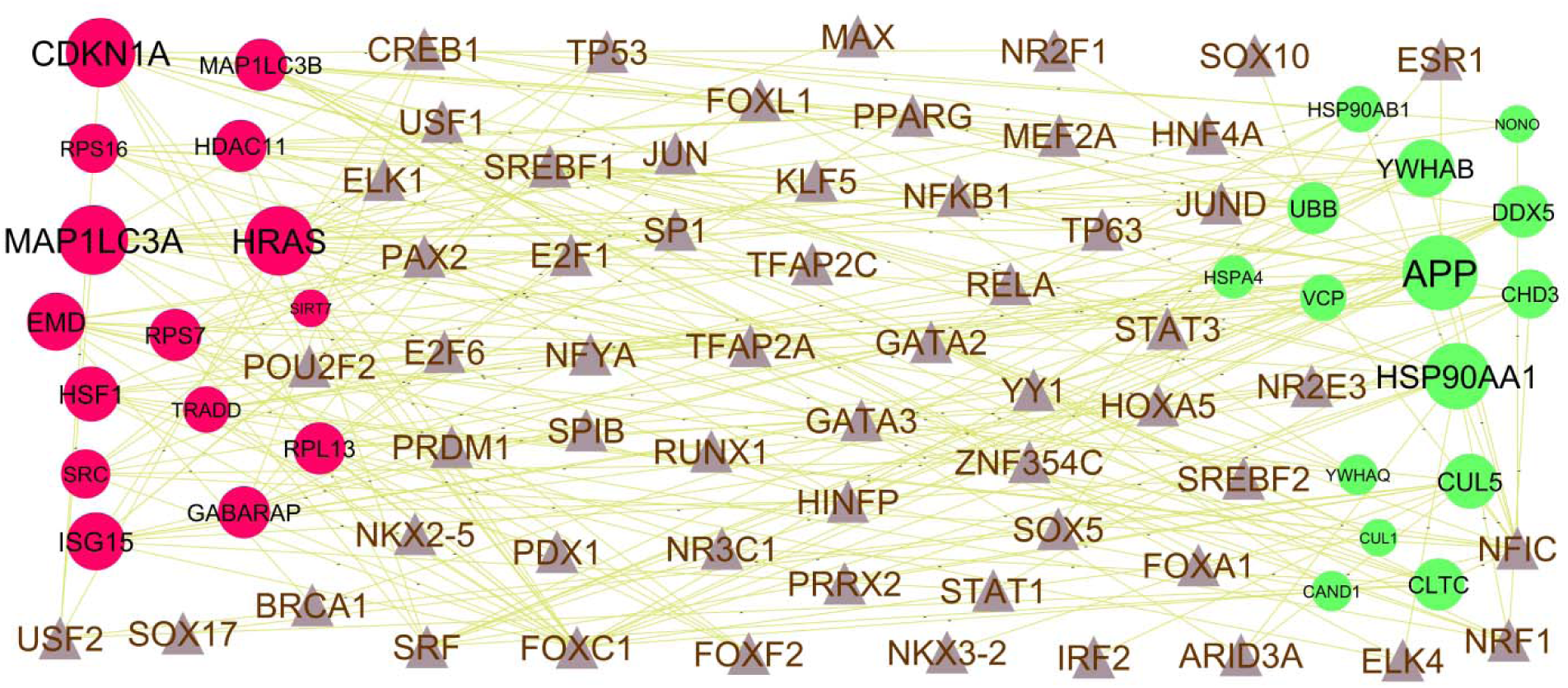
Hub gene - TF regulatory network. The violet color triangle nodes represent the key TFs; up regulated genes are marked in dark green; down regulated genes are marked in dark red.

### Receiver operating characteristic curve (ROC) analysis

ROC curves analysis were performed and the corresponding area under the ROC curve (AUC) was determined to validate the diagnostic value of these 10 hub genes obtained from the PPI network analysis. The larger the area under the ROC curve (AUC), the more the capability of the hub genes to diagnose RIF with excellent specificity and sensitivity. The diagnostic value of hub genes in RIF samples and normal control samples are as follow: APP (AUC:0.910), HSP90AA1 (AUC:0.923), CAND1 (AUC:0.907), CUL1 (AUC:0.901), HSP90AB1 (AUC:0.914), SIRT7 (AUC:0.926), SRC (AUC:0.944), CDKN1A (AUC:0.938), ISG15 (AUC:0.883) and RPS16 (AUC:0.938) (Fig.7).

**Fig. 7.**
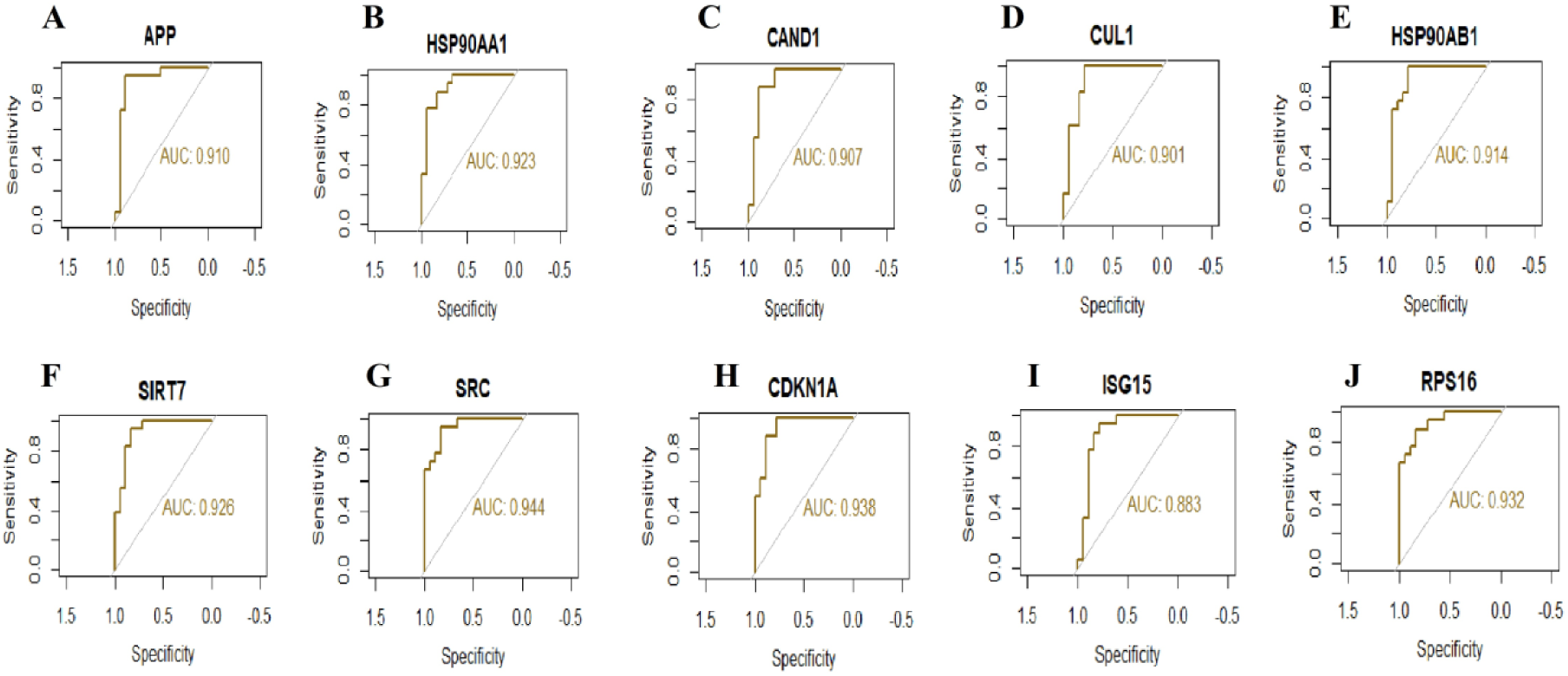
ROC curve analyses of hub genes. A) APP B) HSP90AA1 C) CAND1 D) CUL1 E) HSP90AB1 F) SIRT7 G) SRC H) CDKN1A I) ISG15 J) RPS16.

## Discussion

RIF is a collapse to accomplish a clinical pregnancy after transfer of at least four good-quality embryos in a woman under the age of 40 years [62]. Therefore, sensitive and specific biomarkers of RIF are crucially needed to be identified. In the current investigation, bioinformatics and NGS technology are promising methods to analyze the essential genes and signaling pathways, which might provide novel clues for diagnosis, therapy, and prognosis of RIF. Methodologies related to bioinformatics analyses were used to search the NGS dataset of RIF, and 958 (479 up regulated and 479 down regulated genes) DEGs between RIF and normal control samples were identified based on the GEO database. Recent studies have shown that HBA2 [63], HLA-DRB5 [64], DIO2 [65] and TNC (tenascin C) [66] play important roles in psychosocial consequences. HBA2 [67], FOLR1 [68], DIO2 [69], TNC (tenascin C) [70] and HAP1 [71] are altered expressed in viral infections. HBA2 [72], DIO2 [73], TNC (tenascin C) [74], GAST (gastrin) [75], C2CD4B [76] and CES1 [77] were shown to be involved in the oxidative stress. Several studies have shown that FOLR1 [78], MSLN (mesothelin) [78] and TNC (tenascin C) [79] might contribute to the pathogenesis of endometriosis and could be a potential biomarker for predicting disease progression and therapeutic effects in endometriosis patients. CALCR (calcitonin receptor) [80], DIO2 [81], TNC (tenascin C) [82], HAP1 [83] and CES1 [84] participates in the pathophysiological progression of inflammation. DIO2 [85] and S100P [86] might be implicated in the recurrent spontaneous abortion. DIO2 [87] might serve as molecular marker for insulin resistance. DIO2 [88], TNC (tenascin C) [89] and CES1 [90] have been reported its expression in the obesity. DIO2 [91], MSLN (mesothelin) [92] and S100P [93] plays a role in diagnosis of endometrial cancer. DIO2 [94], TNC (tenascin C) [95], GAST (gastrin) [96] and S100P [97] have been known to be involved in autoimmunity progression. TNC (tenascin C) [70] has a significant prognostic potential in preeclampsia. TNC (tenascin C) [98] is molecular marker for the diagnosis and prognosis of polycystic ovary syndrome. Altered expression of TNC (tenascin C) [99] and GAST (gastrin) [100] significantly correlates with immunomodulation. TNC (tenascin C) [101] might be a potential prognostic marker or therapeutic target for uterine fibroids. Recently, increasing evidence demonstrated that GAST (gastrin) [102] was over-expressed in hyperparathyroidism. Studies had shown that GAST (gastrin) [103] was associated with gestational diabetes. Abnormal regulation of HAP1 [104] is associated with chromosomal abnormalities. HAP1 [105] plays an indispensable role in DNA damage. The results of this investigation suggest that DEGs might play a key role in the pathogenesis of RIF.

GO term and REACTOME pathway enrichment analyses were used to explore the molecular mechanisms involved in the occurrence and development of RIF as well as to determine the potential role of DEGs in the occurrence and development of RIF. Signaling pathways include extracellular matrix organization [106], autophagy [107] and viral infections [21] were responsible for RIF. Recent studies have found that NRCAM (neuronal cell adhesion molecule) [108], JAG1 [109], CD4 [110], MMP2 [111], CDK6 [112], SEZ6L2 [113], SEMA4A [114], CDKN2C [115], EPHA4 [116], ADAM9 [117], ITPR1 [118], CAT (catalase) [119], JAK1 [120], IGF1R [121], PTEN (phosphatase and tensin homolog) [122], CHD4 [123], APP (amyloid beta precursor protein) [124], SRSF1 [125], LRP2 [126], NFAT5 [127], CLEC16A [128], CAPN5 [129], HPSE (heparanase) [130], CLDN4 [131], ANXA1 [132], ISG15 [133], CDKN1A [134], MIF (macrophage migration inhibitory factor) [135], ACE (angiotensin I converting enzyme) [136], GPX1 [137], UCP2 [138], TOB1 [139], KLF15 [140], FKBP1B [141], GDF15 [142], ADGRG3 [143], IRF7 [144], TOM1 [145], CXCR3 [146], IRF5 [147], SLIRP (SRA stem-loop interacting RNA binding protein) [148] and PIK3IP1 [149] have been proved to be related to the pathogenesis of autoimmunity. Studies have revealed that NRCAM (neuronal cell adhesion molecule) [150], LOXL1 [151], CCND1 [152], CYP1B1 [153], CD4 [154], TNFRSF21 [155], TUBB2B [156], PCDH10 [157], MMP2 [158], CDK6 [159], FOXF2 [160], PIAS1 [161], FURIN (furin, paired basic amino acid cleaving enzyme) [162], USP46 [163], ADAM9 [164], DDX5 [165], ABCA1 [166], SLC11A2 [167], JAK1 [168], RBFOX2 [169], IGF1R [170], PTEN (phosphatase and tensin homolog) [171], CHD4 [172], MSH3 [173], PCDHGB7 [174], KDM4A [175], IGF2R [176], PRKAR1A [177], YAP1 [178], SOS1 [179], ARID1A [180], CHD7 [181], EIF4G2 [182], ATM (ATM serine/threonine kinase) [183], PEG10 [184], FAM83B [185], FOLH1 [186], EPM2AIP1 [187], TRIM22 [188], HPSE (heparanase) [189], CLDN4 [190], NUPR1 [191], ISG15 [192], CDKN1A [193], ALDH2 [194], KSR1 [195], MIF (macrophage migration inhibitory factor) [196], ACE (angiotensin I converting enzyme) [197], GPX1 [91], PTGIS (prostaglandin I2 synthase) [198], ATP5F1D [199], RPIA (ribose 5-phosphate isomerase A) [200], BNIP3 [201], FLOT1 [202], HSF1 [203], SIRT7 [204], GDF15 [205], FXYD3 [206], MGST1 [207], MT2A [208], MT1A [208], ROMO1 [209], PFKFB3 [210], FOXA2 [211], SNHG25 [212], ABHD11 [213], CXCR3 [214], FAM83H [215], EMX2 [216], ESRRA (estrogen related receptor alpha) [217] and CCDC124 [218] can contribute to endometrial cancer. NRCAM (neuronal cell adhesion molecule) [219], BMP7 [220], SLC24A4 [221], KIF26B [222], ANPEP (alanylaminopeptidase, membrane) [223], HSD11B1 [224], ADCYAP1R1 [225], CD4 [226], NNAT (neuronatin) [227], TUBB2A [228], SYT7 [229], ADAM22 [230], LSAMP (limbic system associated membrane protein) [231], TNFRSF19 [232], TNFRSF21 [233], LTBP1 [234], MMP2 [235], SEZ6L2 [236], PIAS1 [237], ADGRL1 [238], HSP90AA1 [239], KCNC2 [240], SCARB2 [241], FURIN (furin, paired basic amino acid cleaving enzyme) [242], AGTR2 [243], SLC6A9 [244], EPHA4 [245], PSAP (prosaposin) [246], USP46 [247], UHMK1 [248], ADAM9 [249], SLC1A3 [250], ITPR1 [118], LRRC4C [251], CAT (catalase) [252], UNC13B [253], TUSC3 [254], ABCA1 [255], LRPAP1 [256], WNK3 [257], ADGRB3 [258], JAK1 [259], MAP6 [260], IGF1R [261], ATRN (attractin) [262], CLTC (clathrin heavy chain) [263], EPHA5 [264], MSH3 [265], BACE1 [266], AGO1 [267], APP (amyloid beta precursor protein) [268], CCDC88A [269], RBMX (RNA binding motif protein X-linked) [270], ATRX (ATRX chromatin remodeler) [271], PPP3CA [272], ALDH5A1 [273], MSI2 [274], PJA2 [275], YAP1 [276], HFE (homeostatic iron regulator) [277], KIDINS220 [278], SPAST (spastin) [279], AKAP11 [280], ARID1A [281], ARHGEF11 [282], ATM (ATM serine/threonine kinase) [283], PRSS12 [284], SCRN1 [285], MAPK10 [286], ALDH18A1 [287], NFAT5 [288], ITPR2 [289], GOLGB1 [290], LAMP2 [291], TSPAN5 [292], TTC3 [293], HSP90B1 [294], ASAH1 [295], VCP (valosin containing protein) [296], DLG3 [297], GPC5 [298], CLEC16A [128], M6PR [299], AADAT (aminoadipate aminotransferase) [300], HSPA4 [301], GFM1 [302], CAPN5 [303], ANG (angiogenin) [304], ANXA1 [305], RPS6KA5 [306], ALDH2 [307], TSPO (translocator protein) [308], MIF (macrophage migration inhibitory factor) [309], ACE (angiotensin I converting enzyme) [310], GPX1 [311], FBXL16 [312], UCP2 [313], CLCF1 [314], PRSS57 [315], DDAH2 [316], ARL6IP1 [317], MVP (major vault protein) [318], FLOT1 [319], STYXL1 [320], ATG4D [321], TNK1 [322], MRPL43 [323], CAPN10 [324], SHARPIN (SHANK associated RH domain interactor) [325], HINT2 [326], PEX10 [327], HSF1 [328], SIRT7 [329], ACAT1 [330], PPP3CC [331], ANG (angiogenin) [332], ADH1B [333], GDF15 [334], CEBPD (CCAAT enhancer binding protein delta) [335], RGS10 [336], ELK4 [337], IRF7 [338], TOM1 [339], CXCR3 [340], SNHG7 [341], ZNF787 [342], ARHGEF28 [343], HS1BP3 [344], GPR18 [345], UXT (ubiquitously expressed prefoldin like chaperone) [346], SNRPE (small nuclear ribonucleoprotein polypeptide E) [347], VPS51 [348], COQ8A [349], HDAC11 [350] and PCGF1 [351] are expected to be potential therapeutic targets for psychosocial consequences. BMP7 [352], HSD11B1 [353], CD4 [354], PIAS1 [355], ADAM9 [356], CAT (catalase) [357], ABCA1 [358], JAK1 [359], IGF1R [360], PTEN (phosphatase and tensin homolog) [361], BACE1 [362], APP (amyloid beta precursor protein) [363], IGF2R [364], HFE (homeostatic iron regulator) [365], ARHGEF11 [366], ATM (ATM serine/threonine kinase) [367], COL6A3 [368], NFAT5 [369], EPM2AIP1 [370], ARG2 [371], ALDH2 [372], TSPO (translocator protein) [373], MIF (macrophage migration inhibitory factor) [374], ACE (angiotensin I converting enzyme) [375], UCP2 [376], KLF15 [377], VEGFB (vascular endothelial growth factor B) [378], MSRB1 [379], CAPN10 [380], HSF1 [381], ADH1B [382], GDF15 [383], SOD3 [384], RGS10 [385], AGPAT2 [386], PFKFB3 [387], FOXA2 [388]. IRF7 [389] and CXCR3 [390] have been linked to the development of insulin resistance. Studies have shown that BMP7 [352], F13A1 [391], LOXL1 [392], HSD11B1 [393]. CCND1 [394], WNT16 [395], CYP1B1 [396], JAG1 [397], CD4 [354], NNAT (neuronatin) [398], UBB (ubiquitin B) [399], STRA6 [400], COL18A1 [401], MMP2 [402], CDK6 [403], PIAS1 [355], ADGRL1 [404], KCNC2 [405], SCARB2 [406], CDKN2C [407], RGN (regucalcin) [408], USP22 [409], LAMA1 [410], PRCP (prolylcarboxypeptidase) [411], CAT (catalase) [357], LDB1 [412], ABCA1 [358], EPHA3 [413], IGF1R [414], PTEN (phosphatase and tensin homolog) [415], ATRN (attractin) [416], AMPD2 [417], BACE1 [362], APP (amyloid beta precursor protein) [418], IGF2R [419], MSI2 [420], PRKAR1A [421], SRSF1 [422], YAP1 [423], SOS1 [424], KIDINS220 [425], ARID1A [426], ARHGEF11 [427], LRP2 [428], COL6A3 [429], NFAT5 [369], LAMP2 [430], ARHGAP21 [431], ARMCX3 [432], LANCL1 [433], CAP1 [434], ND2 [435], M6PR [436], GSTM4 [437], CAPN5 [438], HPSE (heparanase) [439], ARG2 [440], ANXA1 [441], ALDH2 [442], TSPO (translocator protein) [443], MIF (macrophage migration inhibitory factor) [374], ACE (angiotensin I converting enzyme) [444], GPX1 [445], MPST (mercaptopyruvatesulfurtransferase) [446], UCP2 [376], KLF15 [447], CDKN1C [448], ND3 [449], PFKP (phosphofructokinase, platelet) [450], HRAS (HRas proto-oncogene, GTPase) [451], VEGFB (vascular endothelial growth factor B) [452], MVP (major vault protein) [453], CAPN10 [454], HSF1 [455], SIRT7 [456], ACAT1 [457]. TKT (transketolase) [458], ADH1B [382], GDF15 [459], CEBPD (CCAAT enhancer binding protein delta) [460], SOD3 [384], PPP1R3G [461], AGPAT2 [462], PFKFB3 [387], FOXA2 [463], IRF7 [389], CXCR3 [390], IRF5 [464], PNPLA2 [465], EMD (emerin) [466], ESRRA (estrogen related receptor alpha) [467] and HDAC11 [468] plays an important role in the development of obesity. Findings have shown that BMP7 [469], HSD11B1 [470], MRC2 [471], CYP1B1 [472], CRISPLD2 [473], CD4 [474], PCDH10 [475], MMP2 [476], CDK6 [477], AGTR2 [478], CAT (catalase) [479], EPHA3 [480], IGF1R [481], PTEN (phosphatase and tensin homolog) [482], MSI2 [483], YAP1 [484], ARID1A [485], MAT2A [486], GSTM4 [487], ANG (angiogenin) [488], CLDN4 [489], ANXA1 [490], CDKN1A [491], ALDH2 [492], TSPO (translocator protein) [493], MIF (macrophage migration inhibitory factor) [494]. ACE (angiotensin I converting enzyme) [Kowalczyńska et al. 2014], GPX1 [495], PTGIS (prostaglandin I2 synthase) [496], KLF15 [497], HSF1 [498], GDF15 [499], PFKFB3 [500], FOXA2 [501], CXCR3 [502] and EMX2 [503] expression is regulated in endometriosis. Recently, increasing evidence demonstrated that BMP7 [504], F13A1 [505], HSD11B1 [506], CYP1B1 [507], CD4 [508], MMP2 [509], CORIN (corin, serine peptidase) [510], CAT (catalase) [511], ABCA1 [512], JAK1 [513], IGF1R [514], PTEN (phosphatase and tensin homolog) [515], ATRX (ATRX chromatin remodeler) [516], YAP1 [517], ARID1A [518], HSD17B6 [519], LIFR (LIF receptor subunit alpha) [520], MTM1 [521], CAPN5 [522], CDKN1A [523], TSPO (translocator protein) [524], MIF (macrophage migration inhibitory factor) [525], ACE (angiotensin I converting enzyme) [375], GPX1 [526], UCP2 [527], CDKN1C [528], BNIP3 [529], VEGFB (vascular endothelial growth factor B) [530], CAPN10 [531], SLC18A2 [532], GDF15 [533], MAP1LC3A [534], IRF7 [535] and SNHG7 [536] were alerted expressed in polycystic ovary syndrome. BMP7 [537], F13A1 [538], ANPEP (alanylaminopeptidase, membrane) [223], LOXL1 [539], HSD11B1 [540], WNT16 [541], CYP1B1 [542], CRISPLD2 [543], JAG1 [544], CD4 [545], NNAT (neuronatin) [546], TGFB3 [547], STRA6 [548], TNFRSF21 [233], CD109 [549], PIK3R3 [550], ABCC4 [551], MMP2 [552], MAFB (MAF bZIP transcription factor B) [553], VANGL2 [554], SEMA4A [555], PIAS1 [355], TNFSF11 [556], HSP90AA1 [557], RGN (regucalcin) [558], AGTR2 [559], USP22 [560], EPHA4 [561], HSP90AB1 [562], PSAP (prosaposin) [563], PRCP (prolylcarboxypeptidase) [564], NR2C2 [565], ADAM9 [566], ITPR1 [567], DDX5 [568], CAT (catalase) [569], ABCA1 [570], ZMPSTE24 [571], JAK1 [359], RRM2B [572], ITGB1 [573], IGF1R [574], SLC39A10 [575], ARHGAP24 [576], PTEN (phosphatase and tensin homolog) [577], UBR3 [578], CLTC (clathrin heavy chain) [579], MSH3 [580], TP53BP1 [581], BACE1 [582], APP (amyloid beta precursor protein) [124], ASAP1 [583], UFL1 [584], ATRX (ATRX chromatin remodeler) [585], FBN2 [586], MSI2 [420], YAP1 [587], HFE (homeostatic iron regulator) [588], SLC26A6 [589], KIDINS220 [590], ARID1A [591], PAG1 [592], EIF4G2 [593], COL6A3 [594], COL4A5 [595], NFAT5 [596], TTC3 [597], EXTL2 [598], ARMCX3 [599], STAT2 [600], CAP1 [601], VCP (valosin containing protein) [602], CUL1 [603], UBA1 [604], TNKS (tankyrase) [605], GPC5 [606], ST3GAL2 [607], MAT2A [608], TRIM22 [609], HSPA4 [301], CAPN5 [610], ANG (angiogenin) [611], HPSE (heparanase) [612], ARG2 [613], NUPR1 [614], ANXA1 [615], ISG15 [616], CDKN1A [134], ALDH2 [617], TSPO (translocator protein) [308], KSR1 [618], MIF (macrophage migration inhibitory factor) [619], ACE (angiotensin I converting enzyme) [620], GPX1 [621], MPST (mercaptopyruvatesulfurtransferase) [622], FBXL16 [312], UCP2 [623], TOB1 [624], KLF15 [625], NAPRT (nicotinatephosphoribosyltransferase) [626], BNIP3 [627], ARL6IP1 [317], MACROD1 [628], VEGFB (vascular endothelial growth factor B) [629], ABCA3 [630], MSRB1 [631], MVP (major vault protein) [632], TNK1 [633], RPL38 [634], DAPK3 [635], HINT2 [636], HSF1 [637], SIRT7 [638], ACAT1 [457], PPP3CC [639], GLTP (glycolipid transfer protein) [640], ANG (angiogenin) [641], CEBPD (CCAAT enhancer binding protein delta) [642], MAFF (MAF bZIP transcription factor F) [643], ADH1C [644], MT2A [645], SOD3 [384], ASCL2 [646], ROMO1 [647], RGS10 [648], ELK4 [337], JPH4 [649], PFKFB3 [650], FOXA2 [651], IRF7 [652], CXCR3 [390], IRF5 [653], SNHG7 [341], PNPLA2 [654], HMGB2 [655], TMUB1 [656], COX6C [657], FAM83H [658], TCFL5 [659], GPR18 [660], RNASEH2A [661], HDAC11 [662] and PCGF1 [351] be a potential therapeutic targets for inflammation. Studies had shown that BMP7 [663], JAG1 [664], ANXA1 [665] and KLF15 [497] were associated with endometrial receptivity. The dysregulation of cellular processes that involve genes include BMP7 [666], CD4 [667], ABCA1 [668], YAP1 [669], GOT2 [670], ARMH4 [671], ISG15 [672], TSPO (translocator protein) [673], MIF (macrophage migration inhibitory factor) [674], UCP2 [675], GDF15 [676], SOD3 [677] and CXCR3 [678] have been linked to the development of immunomodulation. A recent study also showed that BMP7 [679], CYP1B1 [680], CD4 [681], NNAT (neuronatin) [682], SYT7 [683], STRA6 [684], MMP2 [685], SEMA4A [686], PIAS1 [687], RGN (regucalcin) [688], USP22 [689], ADAM9 [690], CAT (catalase) [691], ABCA1 [692], JAK1 [693], RRM2B [572], ITGB1 [573], IGF1R [694], CHD4 [695], ATRN (attractin) [696], BACE1 [697], KDM4A [698], APP (amyloid beta precursor protein) [699], ARL6 [700], YAP1 [701], SOS1 [702], HFE (homeostatic iron regulator) [703], SLC26A6 [704], SPAST (spastin) [705], ARID1A [706], EIF4G2 [707], ATM (ATM serine/threonine kinase) [708], GPX8 [709], SCARA3 [710], LAMP2 [711], FSCN1 [712], AIG1 [713], TTC3 [597], LANCL1 [714], ASAH1 [715], CAP1 [716], GRK3 [717], VCP (valosin containing protein) [718], MAT2A [719], STARD4 [720], CHD3 [721], ANG (angiogenin) [722], ARG2 [723], ANXA1 [724], ISG15 [725], ALDH2 [726], MIF (macrophage migration inhibitory factor) [727], ACE (angiotensin I converting enzyme) [728], GPX1 [621], UCP2 [729], KLF15 [730], CDKN1C [731], CDKN2D [732], BNIP3 [733], MSRB1 [734], NOL3 [735], NUDT1 [736], TXK (TXK tyrosine kinase) [737], MSRB2 [738], ATG4D [739], DAPK3 [740], HINT2 [741], HSF1 [742], SIRT7 [743], TKT (transketolase) [744], ANG (angiogenin) [745], GDF15 [746], CEBPD (CCAAT enhancer binding protein delta) [747], PAGE4 [748], MAFF (MAF bZIP transcription factor F) [749], MT2A [750], MGST1 [751], SOD3 [752], MT1A [753], ROMO1 [647], RGS10 [754], AGPAT2 [755], NOXA1 [756], PFKFB3 [757], FOXA2 [758], IRF7 [759], CXCR3 [760], PNPLA2 [761], HMGB2 [762], NOL3 [763], EMD (emerin) [764], COX6C [657], SLC25A23 [765] and HDAC11 [662] improves oxidative stress.

Several studies suggested that F13A1 [766], CCND1 [767], JAG1 [768], CD4 [226], MMP2 [769], CDK6 [770], RARB (retinoic acid receptor beta) [771], PIAS1 [772], HSP90AA1 [773], SCARB2 [774], FURIN (furin, paired basic amino acid cleaving enzyme) [775], AGTR2 [776], USP22 [777], ADAR (adenosine deaminase RNA specific) [778], HSP90AB1 [562], PRCP (prolylcarboxypeptidase) [779], ADAM9 [780], SLC1A3 [781], DDX5 [782], CAT (catalase) [569], ABCA1 [783], RAB14 [784], ZMPSTE24 [785], JAK1 [786], IGF1R [787], PTEN (phosphatase and tensin homolog) [788], CLTC (clathrin heavy chain) [789], MSH3 [790], MOV10 [791], APP (amyloid beta precursor protein) [792], CCDC88A [793], RBMX (RNA binding motif protein X-linked) [794], UFL1 [795], ATRX (ATRX chromatin remodeler) [796], SRSF1 [797], YAP1 [798], ARID1A [799], ATM (ATM serine/threonine kinase) [800], HACD3 [801], LAMP2 [802], STAT2 [803], ANO6 [804], VCP (valosin containing protein) [805], HCFC2 [806], CUL1 [807], SLC39A9 [808], GOPC (golgi associated PDZ and coiled-coil motif containing) [809], MAP1LC3C [810], API5 [811], DDX24 [812], LARP4 [813], M6PR [814], ST3GAL2 [607], CAND1 [815], MAT2A [816], TRIM22 [817], TRIM5 [818], ARL5B [819], CAPN5 [820], ANG (angiogenin) [821], HPSE (heparanase) [822], TMPRSS13 [823], ARG2 [824], CLDN4 [825], IFI27 [826], ANXA1 [827], ISG15 [828], CDKN1A [829], TSPO (translocator protein) [830], MIF (macrophage migration inhibitory factor) [831], ACE (angiotensin I converting enzyme) [832], GPX1 [833], SPSB2 [834], PRSS8 [835,] KLF15 [836], RILP (Rab interacting lysosomal protein) [837], BNIP3 [838], CAMLG (calcium modulating ligand) [839], ABCA3 [840], MVP (major vault protein) [841], AGO4 [842], DAPK3 [843], HSPBP1 [844], HSF1 [845], ANG (angiogenin) [846], GDF15 [847], SOD3 [848], ZNF683 [849], RGS10 [850], IRF7 [338], CXCR3 [851], SLA2 [852], IRF5 [853], HMGB2 [854], RHEBL1 [855], MAF1 [856], AADAC (arylacetamidedeacetylase) [857], GPR18 [345] and HDAC11 [858] might serve as therapeutic targets for viral infections. F13A1 [859], CYP1B1 [860], CD4 [861], MMP2 [862], FURIN (furin, paired basic amino acid cleaving enzyme) [863], CAT (catalase) [864], ROBO1 [865], JAK1 [866], IGF1R [867], PTEN (phosphatase and tensin homolog)) [868], AGO1 [869], IGF2R [870], CPLANE1 [871], FBN2 [872], YAP1 [873], KIDINS220 [874], PEG10 [875], CUL1 [876], CDKN1A [877], MIF (macrophage migration inhibitory factor) [878], ACE (angiotensin I converting enzyme) [879], GPX1 [880], F2 [881], UCP2 [882], CDKN1C [883], BNIP3 [884], FN3KRP [885], GDF15 [886], ROMO1 [887], CXCR3 [888], IRF5 [889], SNHG7 [890] and TMUB1 [656] might have a significant role in the development of recurrent spontaneous abortion. Recent research indicated that HSD11B1 [891], CD4 [892], ITGA9 [893], MMP2 [894], SLC31A1 [895], GNA12 [896], HSF2 [897], TNFSF11 [898], HSP90AA1 [899], CORIN (corin, serine peptidase) [900], AGTR2 [901], USP22 [902], PRCP (prolylcarboxypeptidase) [903], ADAM9 [902], ABCA1 [904], ITGB1 [905], RBFOX2 [906], IGF1R [907], PTEN (phosphatase and tensin homolog) [908], SLC23A2 [909], MOV10 [910], APP (amyloid beta precursor protein) [911], MYLIP (myosin regulatory light chain interacting protein) [912], YAP1 [908], ARID1A [913], PEG10 [914], NFAT5 [915], VCP (valosin containing protein) [916], ANG (angiogenin) [917], HPSE (heparanase) [918], ARG2 [919], CLDN3 [920], ANXA1 [921], ISG15 [922], CDKN1A [923], MIF (macrophage migration inhibitory factor) [924], ACE (angiotensin I converting enzyme) [925], GPX1 [926], UCP2 [927], CDKN1C [928], BNIP3 [733], DDAH2 [929], RPL39 [930], CAPN10 [931], COPS9 [932], HSF1 [933], ANG (angiogenin) [934], GDF15 [935], SOD3 [936], PPP1R3G [937], TMBIM4 [938], SNHG7 [939] and ANKRD37 [940] might be considered as a pathogenic genetic factors for preeclampsia. The CCND1 [941], CD4 [942], USP22 [943], ADAM9 [944], CAT (catalase) [945], IGF1R [946], KDM4A [947], ATRX (ATRX chromatin remodeler) [271], DNAJC9 [948], CUL1 [949] and HPSE (heparanase) [950] plays a crucial role in establishing chromosomal abnormalities. CYP1B1 [951], CD4 [952], STRA6 [953], PLA2R1 [954], HSP90AA1 [955], DDX5 [956], CAT (catalase) [957], ZMPSTE24 [958], ITGB1 [959], IGF1R [960], PTEN (phosphatase and tensin homolog) [961], CHD4 [962], KDM4A [963], APP (amyloid beta precursor protein) [964], RBMX (RNA binding motif protein X-linked) [965], ATRX (ATRX chromatin remodeler) [966], MSI2 [967], SRSF1 [968], DHX36 [969], YAP1 [970], ARID1A [971], ATM (ATM serine/threonine kinase) [972], HSD17B6 [973], STAT2 [974], VCP (valosin containing protein) [975], UBA1 [976], TNKS (tankyrase) [977], MAT2A [978], CHD3 [979], HPSE (heparanase) [980], CLDN4 [981], ISG15 [982], CDKN1A [983], ALDH2 [984], KSR1 [985], MIF (macrophage migration inhibitory factor) [986], ACE (angiotensin I converting enzyme) [987], GPX1 [988], BNIP3 [989], MVP (major vault protein) [990], FLOT1 [991], NUDT1 [992], COPS9 [993], HSF1 [994], SIRT7 [995], MOB2 [996], TKT (transketolase) [997], ADH1B [998], CEBPD (CCAAT enhancer binding protein delta) [999], SOD3 [1000], IRF5 [1001] and PIERCE1 [1002] were recently found to be a promoter of DNA damage. CD4 [1003], TGFB3 [1004], STRA6 [548], CORIN (corin, serine peptidase) [900], CAT (catalase) [1005], ABCA1 [1006], IGF1R [1007], PTEN (phosphatase and tensin homolog) [1008], IGF2R [1009], SOS1 [1010], SLC26A6 [1011], M6PR [1012], ANG (angiogenin) [1013], CLDN4 [1014], ANXA1 [1015], MIF (macrophage migration inhibitory factor) [1016], ACE (angiotensin I converting enzyme) [1017], CAPN10 [1018], SIRT7 [1019], ANG (angiogenin) [1020] and GDF15 [1021] were found to be linked with gestational diabetes mellitus. TGFB3 [1022], HSF2 [1023], HSP90AA1 [1024], ITPR1 [1025], TAF7L [1026], CAT (catalase) [1027], ABCA1 [1028], PTEN (phosphatase and tensin homolog) [1029], IGF2R [1030], PRKAR1A [1031], DHX36 [1032], YAP1 [1033], HFE (homeostatic iron regulator) [1034], CHD7 [1035], CCDC188 [1036], DCAF17 [1037], ALDH2 [1038], ACE (angiotensin I converting enzyme) [1039], GPX1 [1027], ND3 [1040], TPST2 [1041], ATG4D [1042], UBE2B [1043], MT1A [753] and MTMR14 [1044] could potentially play a role in the onset of male infertility. CDKN2C [1045], SLC26A6 [1046] and ACE (angiotensin I converting enzyme) [1047] have been identified the involvement in hyperparathyroidism. Altered level of IGF1R [1048], PTEN (phosphatase and tensin homolog) [1049], AGO1 [1050], ND2 [1051], MAT2A [1052], HPSE (heparanase) [1053], CLDN4 [1054] and CLDN3[1054] can facilitate RIF. These research findings concur with those from this investigation, which suggest that enriched genes might have a role in the pathophysiology of RIF, and the enrichment analysis of GO terms and pathways also suggests that enriched genes might participate in the autoimmunity, endometrial cancer, psychosocial consequences, insulin resistance, obesity, endometriosis, polycystic ovary syndrome, inflammation, endometrial receptivity, immunomodulation, oxidative stress, viral infections, recurrent spontaneous abortion, preeclampsia, chromosomal abnormalities, DNA damage, gestational diabetes mellitus, male infertility and hyperparathyroidism.

PPI networks of the DEGs in RIF were analyzed by using IMex interactome database. These analyses could help to find some key factors involved in the regulation of RIF and its associated complications. We identified hub genes from the PPI networks. Several hub genes have been reported to be related to RIF and its associated complications. APP (amyloid beta precursor protein) [124], CDKN1A [134] and ISG15 [133] might serve as genetic markers of autoimmunity. Expression of APP (amyloid beta precursor protein) [124], HSP90AA1 [557], CUL1 [603], HSP90AB1 [562], SIRT7 [638], CDKN1A [134], ISG15 [616], HSPA4 [301], CUL5 [1055] and VCP (valosin containing protein) [602] promotes the development of inflammation. APP (amyloid beta precursor protein) [268], HSP90AA1 [239], SIRT7 [329], HSPA4 [301] and VCP (valosin containing protein) [296] are associated to the risk of psychosocial consequences. APP (amyloid beta precursor protein) [363] is found to be associated with insulin resistance. Some studies have shown that APP (amyloid beta precursor protein) [418], SIRT7 [456] and UBB (ubiquitin B) [399] plays a key role in obesity. Altered expression of APP (amyloid beta precursor protein) [699], SIRT7 [743], ISG15 [725] and VCP (valosin containing protein) [718] promotes oxidative stress. A previous study reported that the APP (amyloid beta precursor protein) [792], HSP90AA1 [773], CAND1 [815], CUL1 [807], HSP90AB1 [562], CDKN1A [829], ISG15 [828], CUL5 [1056] and VCP (valosin containing protein) [805] genes were associated with viral infections. Altered expression of APP (amyloid beta precursor protein) [911], HSP90AA1 [899], CDKN1A [923], ISG15 [922], VCP (valosin containing protein) [916] and RBFOX2 [906] are associated with prognosis in patients with preeclampsia. APP (amyloid beta precursor protein) [964], HSP90AA1 [955], SIRT7 [995], CDKN1A [983], ISG15 [982] and VCP (valosin containing protein) [975] are involved in DNA damage progression. HSP90AA1 [1024] gene is a potential marker for the detection and prognosis of male infertility at an early age. A previous study reported that CUL1 [876] and CDKN1A [877] are altered expression in recurrent spontaneous abortion. Research has revealed that CUL1 [949] is expressed in chromosomal abnormalities. SIRT7 [204], CDKN1A [193], ISG15 [192], CUL5 [1057] and RBFOX2 [169] have been found to have a strong association with endometrial cancer. SIRT7 [1019] was an important target gene of gestational diabetes mellitus. CDKN1A [491] has been reported to be expressed in endometriosis. The expression pattern of CDKN1A [523] might offer useful information for treating polycystic ovary syndrome. ISG15 [672] gene is associated with immunomodulation. New biomarkers associated with diagnosis were identified in this investigation: SRC (SRC proto-oncogene, non-receptor tyrosine kinase). RPS16, POLR2B, YWHAQ (tyrosine 3-monooxygenase/tryptophan 5-monooxygenase activation protein theta), YWHAB (tyrosine 3-monooxygenase/tryptophan 5-monooxygenase activation protein beta), RPL13, RPS27L, RPS7 and RPS21. These hub genes might contribute to a better understanding of the molecular mechanisms underlying the pathogenesis of RIF and its associated complications.

We constructed a miRNA-hub gene regulatory network and TF-hub gene regulatory network for these hub genes and screened for a top-scoring miRNAs and TFs. Meanwhile, we found that miRNAs and TFs were involved in the whole process of RIF progression. CLTC (clathrin heavy chain) [263], HSP90AA1 [239], APP (amyloid beta precursor protein) [268], GABARAP (GABA type A receptor-associated protein) [1058], hsa-miR-574-3p [1059], hsa-mir-208a-3p [1060], hsa-miR-218-5p [1061], SREBF1 [1062], BRCA1 [1063], MAX (MYC associated factor X) [1064] and NR2E3 [1065] are known to be involved in the development of psychosocial consequences. Studies have indicated that CLTC (clathrin heavy chain) [579], HSP90AB1 [562], HSP90AA1 [557], APP (amyloid beta precursor protein) [124], CDKN1A [134], GABARAP (GABA type A receptor-associated protein) [1066], CUL5 [1055], DDX5 [568], ISG15 [616], SREBF1 [1067], IRF2 [1068], BRCA1 [1069], MAX (MYC associated factor X) [1070] and NR2E3 [1071] plays a substantial role in inflammation. CLTC (clathrin heavy chain) [789], CAND1 [815], HSP90AB1 [562], HSP90AA1 [773], APP (amyloid beta precursor protein) [792], CDKN1A [829], CUL5 [1056], DDX5 [782], ISG15 [828], GATA3 [1072], IRF2 [1073] and BRCA1 [1074] are associated with the progression of viral infections. HSP90AA1 [899], APP (amyloid beta precursor protein) [911], CDKN1A [923], ISG15 [922], hsa-mir-31-5p [1075], hsa-miR-218-5p [1076], hsa-mir-205-5p [1077] and GATA3 [1078] are a critical biomarkers and are widely involved in preeclampsia. HSP90AA1 [955], APP (amyloid beta precursor protein) [964], CDKN1A [983], DDX5 [956], ISG15 [982], SREBF1 [1079], ARID3A [1080] and GATA3 [1081] were identified to be closely associated with DNA damage, HSP90AA1 [1024] and BRCA1 [1082] can be used as an important therapeutic targets for male infertility. APP (amyloid beta precursor protein) [124], CDKN1A [134], ISG15 [133], hsa-mir-216b-5p [1083], ARID3A [1084], GATA3 [1085], IRF2 [1086] and BRCA1 [1087] were found to be altered expression in autoimmunity patients. APP (amyloid beta precursor protein) [363], SREBF1 [1088] and GATA3 [1089] were significantly associated with insulin resistance. APP (amyloid beta precursor protein) [418], HRAS (HRas proto-oncogene, GTPase) [451], EMD (emerin) [466], hsa-mir-205-5p [1090], SREBF1 [1091], GATA3 [1089] and BRCA1 [1092] are a therapeutic targets for obesity. The altered expression of APP (amyloid beta precursor protein) [699], EMD (emerin) [764], ISG15 [725], SREBF1 [1093], IRF2 [1094], BRCA1 [1092] and NR2E3 [1095] are associated with oxidative stress. CDKN1A [877], GATA3 [1096] and MAX (MYC associated factor X) [1097] have become a potential biomarker for recurrent spontaneous abortion. CDKN1A [193], CUL5 [1057], DDX5 [165], ISG15 [192], SREBF1 [1098], GATA3 [1099], BRCA1 [1100] and MAX (MYC associated factor X) [1101] promotes the initialization and development of endometrial cancer. CDKN1A [491], hsa-miR-574-3p [1102], GATA3 [1103] and BRCA1 [1104] are a potential targets for endometriosis therapy. CDKN1A [523], MAP1LC3A [534], hsa-mir-208a-3p [1105], SREBF1 [1106] and BRCA1 [1107] play important roles in polycystic ovary syndrome growth and progression. ISG15 [672], GATA3 [1089] and BRCA1 [1108] are expected to become a predictive biomarkers for a immunomodulation. hsa-miR-218-5p [1109] and SREBF1 [1110] were significantly associated with prognosis of gestational diabetes mellitus. hsa-mir-205-5p [1111] and SREBF1 [1112] makes a vital regulatory role in endometrial receptivity. Some studies have reported that upregulated GATA3 [1113] and MAX (MYC associated factor X) [1114] correlates with prognosis in hyperparathyroidism. BRCA1 [1115] vital for the development of chromosomal abnormalities. There were intersected to finally obtain novel biomarkers related to RIP and its associated complication, which included the following: RPS16, MAP1LC3B, SRC (SRC proto-oncogene, non-receptor tyrosine kinase), YWHAB (tyrosine 3-monooxygenase/tryptophan 5-monooxygenase activation protein beta), hsa-mir-382-5p, miR-218-1-3p, hsa-mir-548ad-5p, hsa-mir-6132, RELA (RELA proto-oncogene, NF-kB subunit), TFAP2C and SPIB (Spi-B transcription factor). In short, focusing on miRNAs and TFs regulations could provide potential approaches for the treatment of RIP and its associated complication.

### Conclusions

In the current investigation, we identified 958 significant DEGs in RIF. Therefore, the results in this investigation provided reliable key genes and signaling pathways for RIP and its associated complication, which will be useful for RIF molecular mechanisms, diagnosis and candidate targeted treatment. However, further investigations are needed to investigate the potential impact of these genes and signaling pathways on RIP progression, in order to fully validate their role in RIP.

## Acknowledgement

I thanks very much to Fan Y, Shi C, Huang N, Fang F, Tian L, Wang J, Peking University People’s Hospital, Beijing, China, the authors who deposited their NGS dataset GSE243550, into the public GEO database.

## Ethical approval

This article does not contain any studies with human participants or animals performed by any of the authors.

## Informed consent

No informed consent because this study does not contain human or animals participants.

## Availability of data and materials

The datasets supporting the conclusions of this article are available in the GEO (Gene Expression Omnibus) (https://www.ncbi.nlm.nih.gov/geo/) repository. [(GSE243550) https://www.ncbi.nlm.nih.gov/geo/query/acc.cgi?acc=GSE243550]

## Consent for publication

Not applicable.

## Author Contributions

B. V. - Writing original draft, and review and editing

C. V. - Software and investigation

